# Interactome Analysis of Visceral Adipose Tissue Elucidates Gene Regulatory Networks and Novel Gene Candidates in Obesity

**DOI:** 10.1101/2023.12.21.572734

**Authors:** Lijin Wang, Pratap Veerabrahma Sesachalam, Ruiming Chua, Sujoy Ghosh

**Author notes:** CONTACT INFO: Sujoy Ghosh, Laboratory of Bioinformatics and Computational Biology, Pennington Biomedical Research Center, Baton Rouge, LA 70808, USA. DISCLOSURE: The authors declared no conflict of interest.

## Abstract

**Objective:** Visceral adiposity is associated with increased proinflammatory activity, insulin resistance, diabetes risk and mortality rate. Numerous individual genes have been associated with obesity, but studies investigating gene-regulatory networks in human visceral obesity are lacking.

**Methods:** We analyzed gene-regulatory networks in human visceral adipose tissue (VAT) from 48 obese and 11 non-obese Chinese subjects using gene co-expression and network construction with RNA-sequencing data. We also conducted RNA interference-based tests on selected genes for adipocyte differentiation effects.

**Results:** A scale-free gene co-expression network was constructed from 360 differentially expressed genes between obese and non-obese VAT (absolute log fold-change >1, FDR<0.05) with edge probability >0.8. Gene regulatory network analysis identified candidate transcription factors associated with differentially expressed genes. Fifteen subnetworks (communities) displayed altered connectivity patterns between obese and non-obese networks. Genes in pro-inflammatory pathways showed increased network connectivities in obese VAT whereas the oxidative phosphorylation pathway displayed reduced connections (enrichment FDR<0.05). Functional screening via RNA interference identified *SOX30* and *OSBPL3* as potential network-derived gene candidates influencing adipocyte differentiation.

**Conclusions:** This interactome-based approach highlights the network architecture, identifies novel candidate genes, and leads to new hypotheses regarding network-assisted gene regulation in obese vs. non-obese VAT.

*What is already known about this subject?:*

- Visceral adipose tissue (VAT) is associated with increased levels of proinflammatory activity, insulin resistance, diabetes risk and mortality rate.
- Gene expression studies have identified candidate genes associated with proinflammatory function in VAT.

*What are the new findings in your manuscript?:*

- Using integrative network-science, we identified co-expression and gene regulatory networks that are differentially regulated in VAT samples from subjects with and without obesity
- We used functional testing (adipocyte differentiation) to validate a subset of novel candidate genes with minimal prior reported associations to obesity

*How might your results change the direction of research or the focus of clinical practice:*

- Network biology-based investigation provides a new avenue to our understanding of gene function in visceral adiposity
- Functional validation screen allows for the identification of novel gene candidates that may be targeted for the treatment of adipose tissue dysfunction in obesity

## INTRODUCTION

Although obesity’s role in cardiovascular and metabolic diseases is well recognized, not all obesity forms are associated with similar risks of disease (1). Some individuals with high body fat appear to be still protected from obesity-related complications, that can be partly explained by associating disease with the topography of fat distribution and adipose tissue quality, rather than body fat percentage or total body weight (2–4). Specifically, visceral adipose tissue (VAT) accumulation poses significant health risks, including inflammation, insulin resistance, diabetes, dyslipidemia, hypertension, atherosclerosis, and increased mortality (5, 6). Factors such as ethnicity, age, gender, genetics, and the gut microbiome etiologically influence VAT variation (7, 8). Despite these known associations, the molecular mechanisms linking visceral fat to health risks are poorly understood. Genetic studies have linked several loci with visceral fat indices such as waist-to-hip ratio (WHR) or waist circumference (WC) (9–11), but the biological mechanisms linking these loci to VAT function remain largely unclear. Transcriptomic studies on human visceral fat are fewer than those on subcutaneous fat, and while such studies have generated important insights into gene function in VAT, the full scope of transcriptomic impact has not been fully explored (12–14).

Network biology, analyzing binary interactions among biomolecules (genes, proteins, or metabolites), provides a complementary approach to understanding cellular mechanisms, and can provide important information on biological states even when individual network components may not show statistically significant changes (15). A related approach known as pathway enrichment analysis also provides collective information based on well-defined biological processes, but lacks the topological association among genes within and between pathways. Additionally, over 30% of genes lack pathway associations, further limiting the scope of discoveries (16, 17). In contrast, network analysis considers gene topologies regardless of pathway membership, offers a broader view of the interactome, and can span multiple pathways. Network topology variations are also disease-relevant, such as that reported for type 1 and 2 diabetes, obesity and several cancers (18).

Network analysis studies typically involves creating condition-specific networks from experimental data, such as gene co-expression networks implying co-expression with possible functional relatedness, or via mapping genes to external networks (e.g., protein-protein interactions or gene regulatory networks) for augmented biological predictions (19, 20). Our study employs both methods to understand obesity-related changes in the VAT transcriptome. We aim to (i) explore how gene-gene associations and affected biological processes change with obesity in VAT, and (ii) examine alterations in transcription factor-gene interactions (gene regulatory networks) and their impact on network community structures in obese and non-obese VAT. We further prioritize genes and transcription factors identified through these methods, and employ literature and disease database searches to identify putatively novel gene candidates associated with visceral obesity. Selected candidates are further tested via an RNA-interference mediated functional screening assay for lipid-accumulation in human adipocytes.

## RESEARCH DESIGN AND METHODS

### Ethics Statement

The project was approved by the ethical committee of the First Affiliated Hospital of Nanjing Medical University (2016-SR-220). Informed written consent was obtained before gastric bypass surgery.

### Study population

Visceral adipose tissue was collected from 60 Chinese subjects undergoing gastric bypass surgery at the First Affiliated Hospital of Nanjing Medical University. Samples were stored at –80℃ until further use. Demographics data and metabolic traits for all subjects are summarized in **Table 1**, and reported in further detail in a previous publication (14).

**Table 1.** Demographic and clinical characteristics of the study cohort.

| Clinical parameter | Group |  | P-value |
| --- | --- | --- | --- |
|  | Obese<br>(F/M; 27/21) | Non-obese<br>(F/M; 9/2) | ObesevsNon-obese |
| Age (years) | 28.29±5.87 | 41.18±11.35 | <0.001 |
| Waist (cm) | 129.87±17.67 | - | - |
| Hip (cm) | 130.31±13.03 | - | - |
| WHR | 0.99±0.07 | - | - |
| BMI (kg/m<sup>2</sup>) | 41.92±7.06 | 22.60±1.97 | <0.001 |
| Systolic pressure (mmHg) | 136.81±18.45 | 117.09±9.99 | <0.001 |
| Diastolic pressure (mmHg) | 82.54±13.91 | 75.91±9.26 | 1.98E-01 |
| HbA1c (%) | 6.84±2.16 | - | - |
| Total cholesterol (mmol/L) | 4.47±0.74 | 4.62±0.98 | 9.46E-01 |
| TAG (mmol/L) | 1.73±1.13 | 0.86±0.17 | 1.00E-03 |
| HDL (mmol/L) | 1.03±0.20 | 1.38±0.43 | 1.30E-02 |
| LDL (mmol/L) | 3.30±0.62 | 3.13±0.63 | 3.55E-01 |
| Fasting glucose (mmol/l) | 7.18±3.32 | 5.04±0.41 | 9.00E-03 |
| Uric acid (umol/L) | 409.63±139.14 | 284.51±65.15 | 1.00E-03 |
| Leptin (ug/L) | 6.33±4.53 | - | - |
| C-reactive protein (mg/L) | 8.44±5.73 | - | - |
| 25-OHVitD (nmol/L) | 33.73±16.93 | - | - |
| HOMA-IR | 8.54±5.53 | - | - |
| HOMA-B | 247.63±191.36 | - | - |

Value of clinical parameter are given as means ± SD. *P*-value is calculated by Mann-Whitney test.

Based on the recommendations for obesity in Chinese adults (21), we classified participants with BMI >= 28 as subjects with obesity, whereas subjects with BMI < 28 were considered as non-obese.

### Sample preparation and RNA-seq

The Qiagen(Qiagen, Hilden, Germany) miRNeasy kit was used for extraction of RNAs of visceral adipose tissues. After quality evaluation of all samples, one sample with low quality RNA in the non-obese group was dropped. RNA-seq libraries were prepared using NEBNext® UltraTM RNA Library Prep Kit from Illumina (San Diego, CA, USA). RNA sequencing was carried out on the Illumina Hiseq 4000 sequencer.

### Gene Expression Profiling

Quality of the paired-end RNA sequencing reads was evaluated via FastQC (v0.11.5) (https://www.bioinformatics.babraham.ac.uk/projects/fastqc/). STAR (v2.5.3) was applied to align the 150bp paired-end reads of 59 subjects to the human genome (GRCh37) (22).

Aligned reads were further sorted via Samtools (23) and read counts were summarized at the gene level via FeatureCounts (v1.5.0) (24).

### Identification of differentially expressed gene

Differentially expressed genes (DEGs) between obese and non-obese VAT were identified using the R packages *edgeR* (version 3.14.0) and *limma* (version 3.37.7) (25, 26) after adjustments for age and sex. Raw read counts were TMM-normalized in *edgeR*, and genes with >1 counts-per-million in at least 20% of the samples was retained for further analysis. Sample outliers were investigated through principal components analysis via the *stats* package in R (version 3.6.2). Multiple testing correction was performed via estimation of the false discovery rate (FDR). Genes with absolute log2 (fold change) > 1 and FDR < 0.05 were considered as differentially expressed.

### Gene Co-expression network analysis

Gene co-expression networks were constructed through the Graphical Gaussian model-based GeneNet package via partial correlations (27). GeneNet employs a computationally efficient learning algorithm to estimate correlation networks reliably from small sample numbers, making it suitable for the present study. In the network, each gene corresponds to a node and the edges portray direct dependencies between the genes. Subnetworks with probability of an edge being present > 0.8 were selected and further analyzed in Cytoscape (28). Two key network centrality indices, degree and betweenness, were calculated via Centiscape (29). The degree of a network node refers to the number of nodes connected to that node, with high-degree nodes functioning as essential hubs and regulating multiple processes (30). Betweenness of a node quantifies the number of shortest paths between all pairs of nodes passing through that node, and is often associated with genes regulating the flow of information through the network (31). The scale-free architecture of the gene co-expression network was tested via the power-law function in the R package *igraph* (https://igraph.org). A Kolmogorov-Smirnov test p-value <0.05 was used to reject the hypothesis that the observed data could have been drawn from the fitted power-law distribution (32).

### Regulatory network construction

Gene-transcription factor (TF) and gene-gene interactions were constructed via PANDA (Passing Attributes between Networks for Data Assimilation) (33), involving an integrative message-passing procedure to reconstruct gene regulatory networks. Instead of relying only on TF-gene co-expression values, PANDA investigates shared patterns of co-expression between common targets of a transcription factor to derive edge weights. The PANDA-informed gene-regulatory networks were ultimately constructed from three sources of inputs – a user-defined gene-list, protein-protein interaction data, and transcription factor binding motifs (TFBMs), the latter two being obtained from VAT specific data available in GRAND (https://grand.networkmedicine.org/tissues/Adipose_visceral_tissue/) (34). Through this approach, we generated two types of gene regulatory networks. In one version (DEG-PANDA), a TF-gene regulatory network was constructed based on significantly differentially expressed genes (DEGs) between obese and non-obese VAT (|log2 (fold change)| > 1 and adj.P-value < 0.05). In another version, we constructed separate obese and non-obese gene regulatory networks based on all expressed genes in each group (OL-PANDA). The edge weights of the PANDA derived networks reflected the likelihood of a gene’s regulation by its associated TF (only positive edge weights were considered). A total of 644 transcription factors was used for deriving the networks in both cases.

### Differential network analysis via comparison of community structures

Biological networks are characterized by their modularity, often comprising of closely linked groups of genes responsible for similar cellular functions (communities). We investigated community structures in the obese and non-obese networks (OL-PANDA) via the ALPACA application (ALtered Partitions Across Community Architectures) as implemented in netZooR v 1.2.2 (35). Through ALPACA we identified separate gene modules that displayed significantly different within-group connectivities between obese and non-obese networks.

### Pathway enrichment analysis in network communities

The pathway over-representation analysis tool Enrichr was used to perform enrichment analyses on gene subsets identified through PANDA and ALPACA (36). Pathway enrichment was ascertained on gene-sets from the Hallmark repository. Pathways containing >=5 genes and an enrichment FDR<=0.05 were considered as significantly enriched.

### Candidate gene selection

From each of the individual analyses described above, we selected genes with the top 10 scores, resulting in 80 genes (some genes ranked in the top 10 in multiple analyses). A total of 9 analysis categories were considered including (1) differential gene expression (limma) p-values, (2) maximum absolute partial correlaton scores (GeneNet), (3) median absolute partial correlation scores (GeneNet), (4) degree distribution (Centiscape), (5) betweenness distribution (Centiscape), (6) top genes identified in DEG-PANDA (from a robust ranking of the frequencies of gene-gene and gene-TF associations and median estimates for weighted edge scores of gene-gene and gene-TF associations), (7) top transcription factors identified by DEG-PANDA (from a robust ranking of the frequencies of TF-gene and TF-TF associations and median estimates for weighted edge scores of TF-gene and TF-TF associations), (8) ALPACA-based gene-level modularity scores, and (9) Alpaca-based TF-level modularity scores. Robust ranking of PANDA-based gene and transcription factor candidates was performed on the ranks of the individual analyses via the *Robustrankaggreg* package (version 1.2.1) in R.

### Literature, disease database and GWA study searches

The 80 candidate genes were used as input for querying literature, gene-disease associations, and genetic association studies (GWAS) to identify known associations to obesity. Gene selection was performed heuristically, by focusing on genes with minimal prior association to adiposity and with similar representations for transcription factors and non-TF categories. For querying the published literature, we used the latest version of Pubmed (https://pubmed.ncbi.nlm.nih.gov/) with the following keywords as search terms: *((“Obesity”[Title/Abstract] OR “Adiposity”[Title/Abstract] OR “Fat”[Title/Abstract] OR “Body Mass Index”[Title/Abstract] OR “Adipose”[Title/Abstract] OR “Adipocyte”[Title/Abstract] OR “BMI”[Title/Abstract] OR “Obese”[Title/Abstract]) AND “gene-symbol”[Title/Abstract]) AND (2010/1/1:3000/12/12[pdat])*

For each gene, the number of Pubmed records satisfying the search terms was obtained along with an indication of any ambiguity in the search results, such as the confounding of a gene symbol with abbreviation for an unrelated biological process (e.g. the FEV gene confounded with ‘forced expiratory volume’ in the Pubmed abstracts). In addition to the literature-based Pubmed searches, we also queried the DisGenet database (www.disgenet.org) (37) to identify the number of diseases associated with each query gene. For each gene, we tallied the number of diseases linked to that gene and estimated a ‘disease-specificity index’ (DSI) for each gene, defined as;

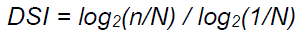

where n = number of disease associated genes and N = total number of diseases in Disgenet. A larger DSI indicated a relatively lesser number of gene-associated diseases and vice versa. We further identified the number of obesity-related diseases associated with the query genes by selecting [*\*obesity\**] as Disease category in the Disgenet output (wildcarded by the ‘*’ character). Finally, we also downloaded and directly queried the GWAS catalog (https://www.ebi.ac.uk/gwas) for evidence of single nucleotide polymorphisms (SNPs) within or near the query genes for their association to adiposity-related phenotypes. Only SNPs with GWAS-association P-value <=10E-05 were considered for this analysis.

### Functional analysis in human adipocytes

Human adipose-derived stem cells were isolated from WAT of patients undergoing bariatric surgery (approved by the Domain Specific Review Board at National Healthcare Group, Singapore) according to methods described previously (38). ASCs were differentiated in 96-well gelatin-coated plates, beginning at 48 hrs after 100% confluence (Day 0), with an adipogenic cocktail consisting of 0.5mM isobutylmethylxanthine (IBMX, Sigma, MO, USA), 1uM dexamethasone (Sigma), 172 nM insulin (Gibco) and 100 uM indomethacin (Sigma), in the presence of differentiation media (low glucose DMEM, 10% FBS, 5% Pen-strep, 5% NEAA, all obtained from Thermo Fisher Scientific, MA, USA) until day 6, with refreshment of media on day 3. On Day 6, the adipogenic cocktail was changed to 1uM dexamethasone (Sigma), 172 nM insulin (Gibco) together with the same differentiation media until Day 12 with refreshment of media on Day 9. Gene knockdown in differentiating adipocytes was performed via siRNA transfection on Days 0, 3, 6 and 9 of differentiation. Briefly, siRNAs were designed against selected genes via the SMARTpool platform and transfected into cells via Accell siRNA technology (Dharmacon, Colorado, USA) through an in-house developed custom protocol. A control (mock) transfection with non-targeting Accell-NT siRNA was also performed. Intracellular lipid accumulation was quantified by the Adipored adipogenesis assay reagent (Lonza, Basel, Switzerland). On day 12, cells were washed once in Hanks Balanced Salt Solution (HBSS), and then incubated in the dark with Adipored (pre-diluted in HBSS at a ratio of 40:1) for 30 minutes at room temperature. Dye fluorescence was quantified by a Tecan 200 spectrofluorometer with an excitation wavelength of 485nm and emission wavelength of 572nm. Cell nuclei were quantified by incubation with the Hoeschst 33342 ds-DNA binding stain (BD Biosciences, NJ, USA). For nuclear staining, Adipored dye was removed from the cells, followed by incubation with Hoechst 33342, diluted 1:2500 in HBSS for 30 minutes in the dark at room temperature without washing. Hoechst 33342 florescence was measured in the Tecan 200 spectrofluorometer with an excitation wavelength at 368nm and emission wavelength at 465nm. The average amount of lipid content per cell was estimated by dividing the Adipored based value per well by the Hoechst 33342 derived value for the same well. The time-course of expression of selected genes throughout adipocyte differentiation was performed via quantitative RT-PCR on total RNA isolated from adipocytes from Day 0 to Day 12 of differentiation. Gene expression was quantified via SybrGreen (Biorad, USA) and normalized to expression of peptidylprolyl isomerase B (PPIB or S-Cyclophilin) as a housekeeping gene. Changes in gene expression during adipocyte differentiation were quantified with respect to gene expression on Day 0 by the ΔΔCt method (39). The qPCR primer sequences for all genes are provided in Supplementary **Table S1.**

### Data analysis

All data analysis related to differential gene expression, gene-co-expression network construction, bipartite gene-gene and TF-gene regulatory network analysis, network community identification and candidate gene prioritization were carried out through application-specific software packages written in the R statistical computing language (version 4.3.2, https://www.R-project.org/). For functional testing experiments, data was analyzed either in Microsoft Excel (2019) or through the *tidyverse* package (version 2.0) in R. Results are expressed as mean(±SD) unless otherwise indicated.

## RESULTS

### Gene co-expression networks in visceral fat from individuals with or without obesity

A large-scale undirected gene-co-expression network was constructed based on 410 genes (identified by limma), showing significant expression differences between obese and non-obese samples (absolute log fold-change >=1, FDR <= 0.05). The network, pruned to only retain high-likelihood gene-gene partial correlations (probability >0.8), resulted in 2489 binary interactions involving 360 genes, with 1562 positive associations indicating similar expression directions. Power-law analysis was consistent with a scale-free network architecture (KS p-value = 0.82). **(Fig. 1A)**. A map of the top 100 interacting genes (based on absolute pair-wise partial correlations) is shown in **Fig. 1B** (full list available in Table S2). Some of the strongest partial correlations (r >=0.2) were observed between *MUC20-MUC4, LINC01230-DMRT2*, and *AL049634.2-SIRPB1* gene pairs. We then analyzed centrality indices (degree and betweenness) for the network nodes. There was a positive association between degree and betweenness centrality **(Fig. S1)**. The top 5 genes with the highest connectivity were *IGHD, IGFN1, PKP3, ATP8A2,* and *PHACTR3* (degree ranging between 47-57), while *IGFN1, IGHD, PHACTR3, PKP3*, and *CLDN9* had the top 5 highest betweenness scores (3859–5721). Among lincRNAs, *LINC01956* and *LINC00942* showed high degree (>30) and betweenness (>1900), with *LINC01956* additionally connected to ATP8A2 and IGHD (identified earlier as highly connected nodes). The full centrality measures for all genes are shown in **Table S3**.

**Figure 1.**
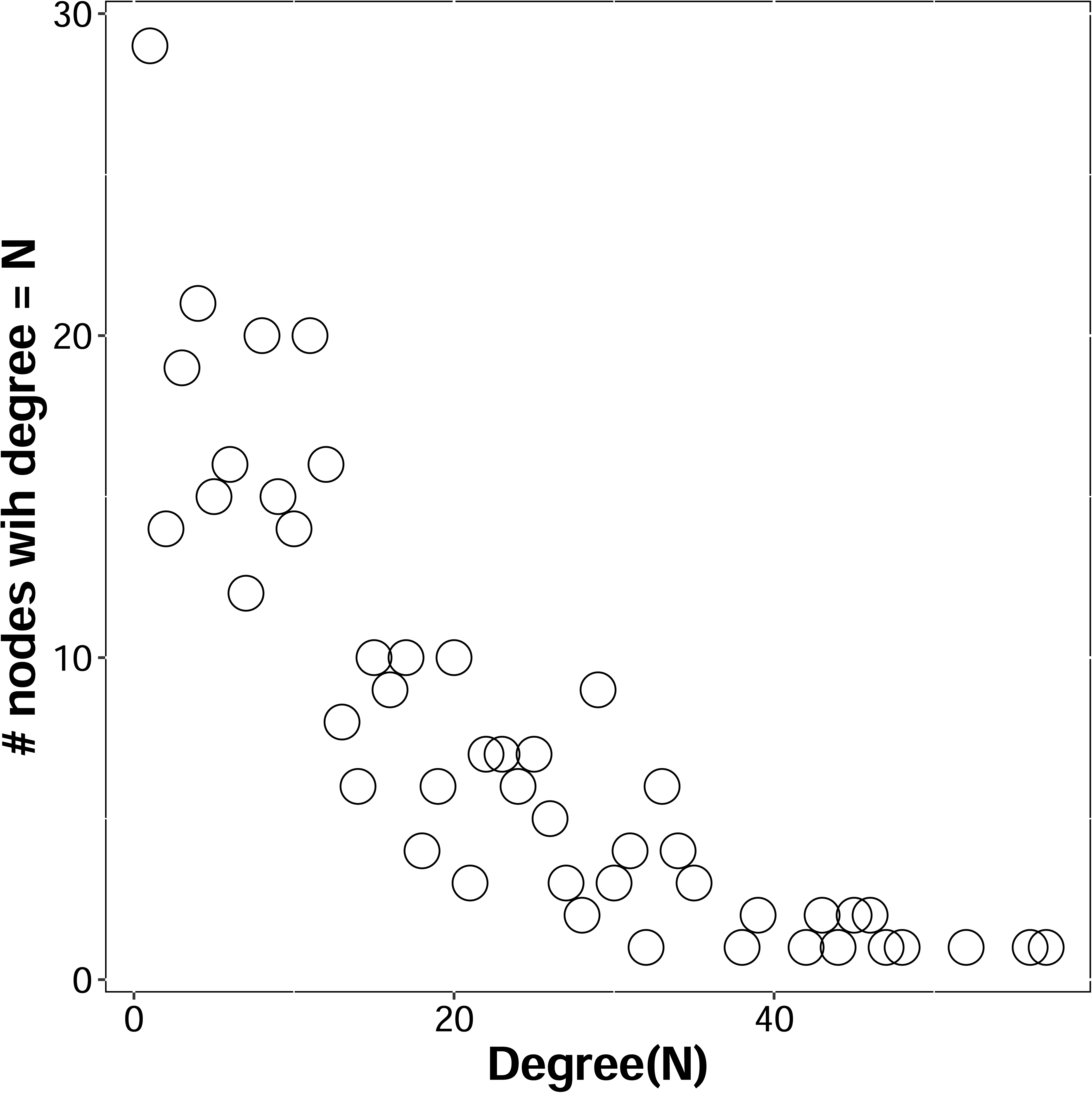
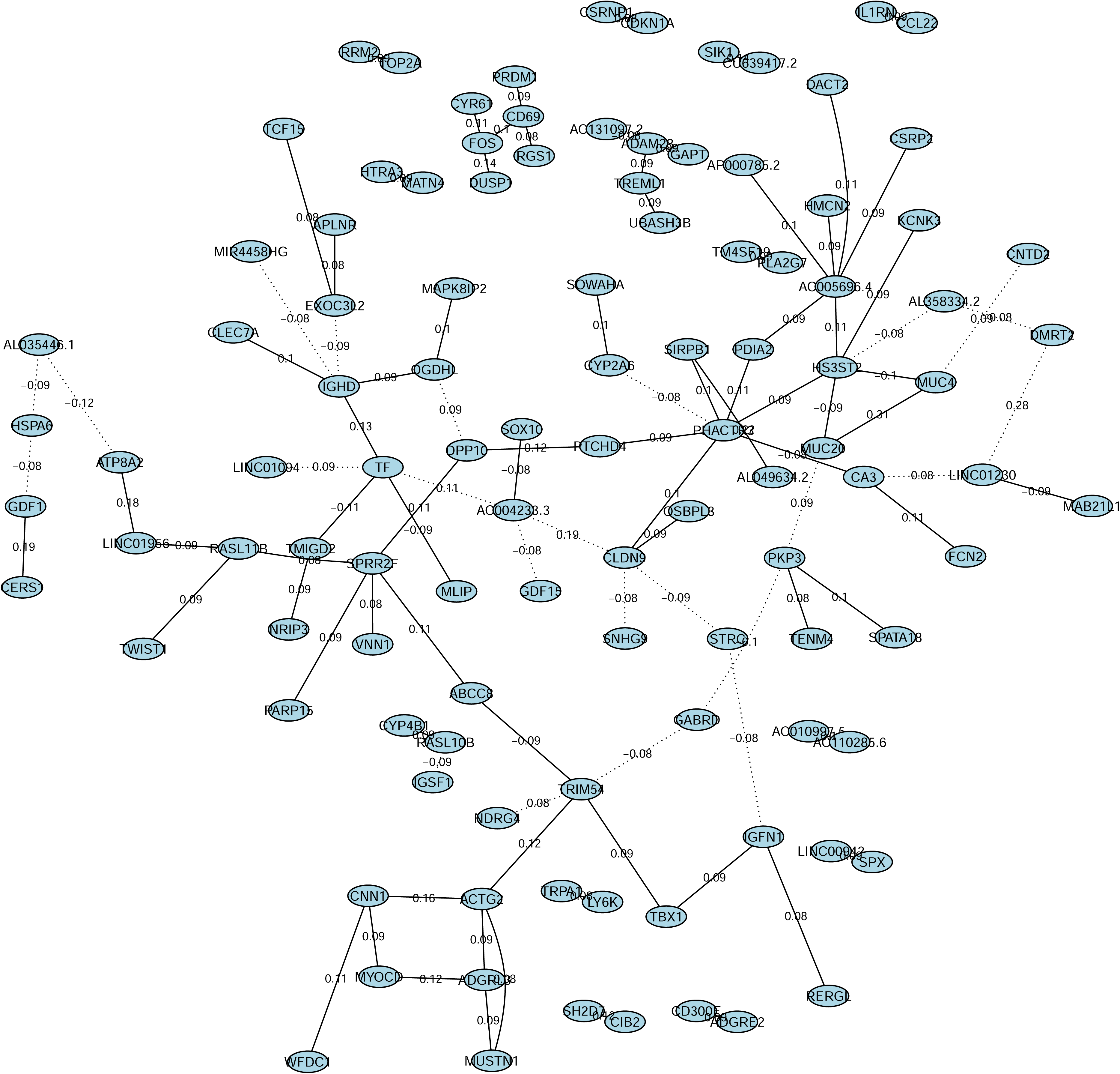
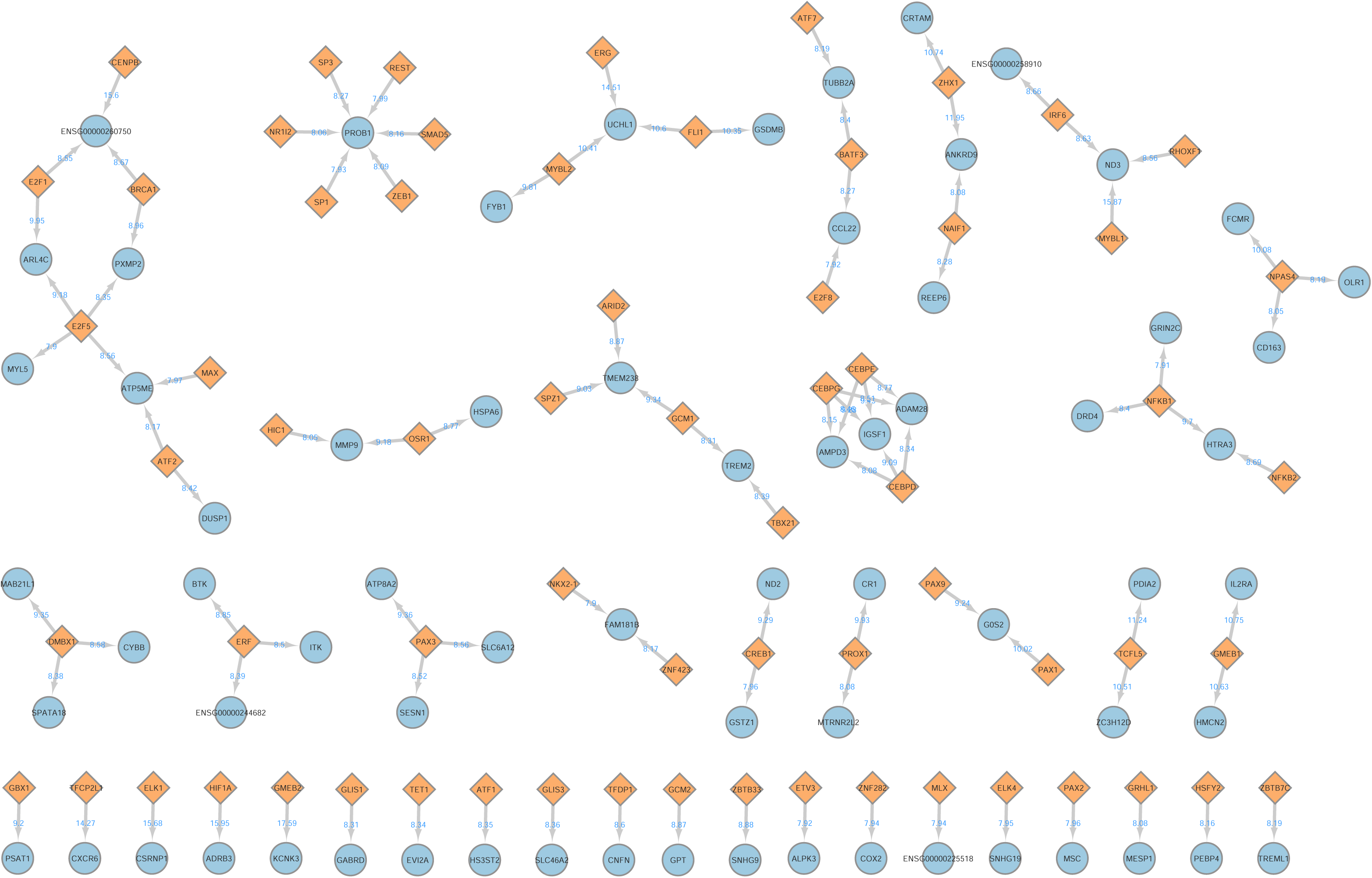
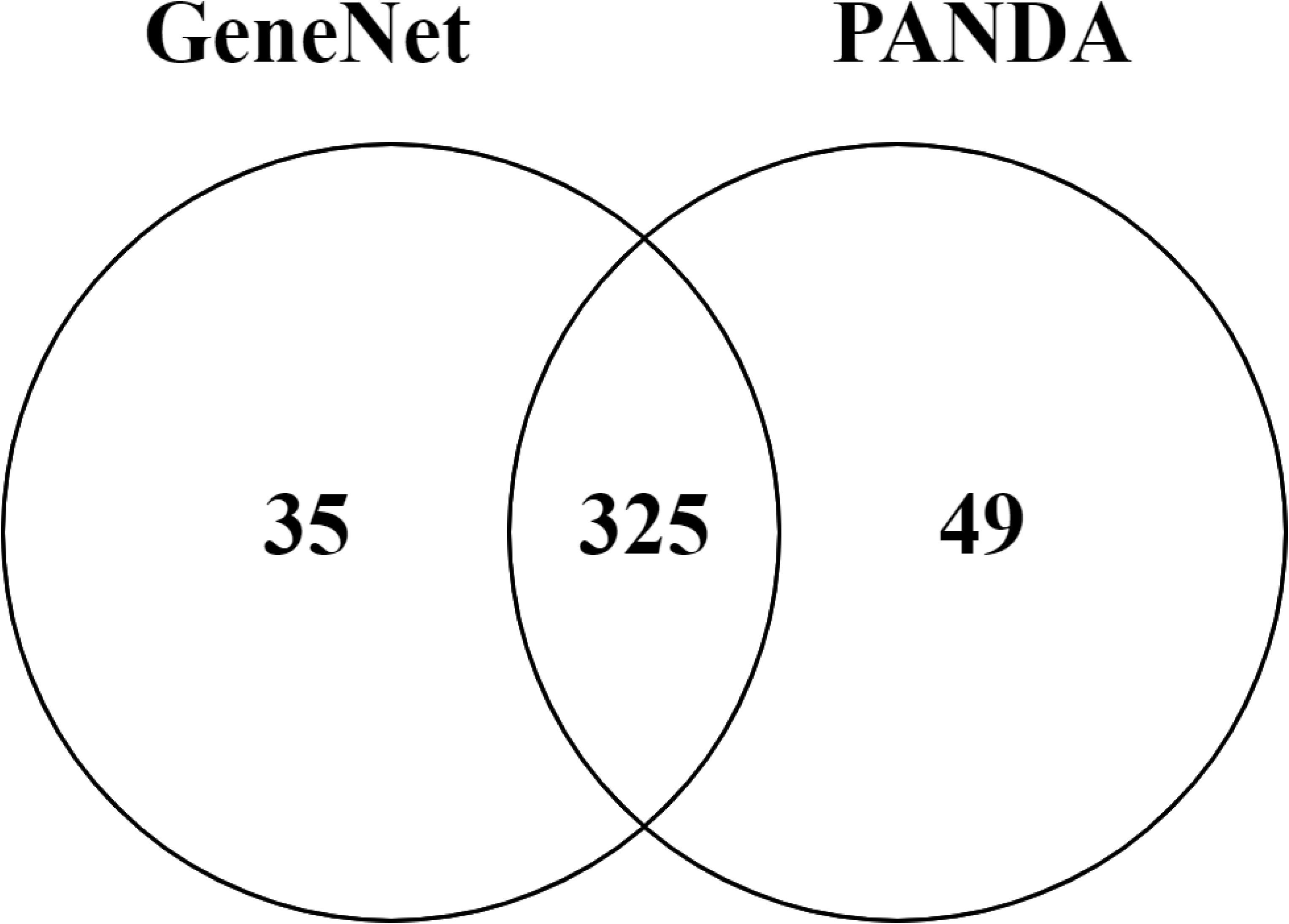
Co-expression networks from genes differentially expressed between obese and non-obese VAT. (A) Power-law fit of network degree (x-axis) vs. degree distribution (y-axis), to test for scale-free architecture of network. **(B)** Subset of the co-expression network constructed in GeneNet from the top 100 edge weights (absolute partial correlations. In the undirected network, nodes are annotated with gene names and the partial correlation between node pairs are indicated on the edges. **(C)** Subset of the directed gene regulatory network consisting of the top 100 transcription factor to gene (TF-gene) interactions, as derived by PANDA analysis of differentially expressed genes. TFs are represented by purple triangles, and target genes by blue circles. **(D)** Overlap among significant interactors in networks derived from co-expression (GeneNet) and TF-gene association (PANDA). For GeneNet network, 360 genes with edge probability >0.8 were considered. For TF-gene regulatory network, 374 genes with PANDA-based interaction score >0 were included.

### Gene-gene and TF-gene connectivities in non-obese and obese VAT networks

We utilized PANDA to create directed transcriptional regulatory network models, integrating transcription factor (TF) motif data, protein-protein interactions, and gene expression results. DEG-PANDA identified top TFs like *FOXP1, FOXP4*, *ARID3A, BPTA* and *IRF5* as the top 5 TFs with the greatest number of DEG interactions, considering both up– and down-regulated genes. For genes upregulated only in obese VAT, *IRF3* and *PRDM1* displayed the highest gene connectiities, while *SP1* was linked to the largest number of downregulated genes. (full list of TF-DEG and DEG-DEG connectivities are presented in **Table S4**). The top 100 TF-gene interactions from DEG-PANDA are shown in **Figure 1C**. Comparing DEG-PANDA’s top connected DEGs with the top genes from GeneNet’s gene-co-expression network revealed significant overlap, with 325 genes in common from 360 GeneNet and 374 DEG-PANDA genes **(Fig. 1D)**. In addition to gene regulatory networks with DEGs, we also constructed similar TF-based networks separately for genes expressed in obese and non-obese VAT. This OL-PANDA analysis identified *IRF8, PAX8, ARNT, KDM2B* and *ATF4* as the top 5 TFs with the greatest gains in gene associations in obesity, while *YY1*, *MEF2D*, *STAT3*, *GSC* and *ELF2* were the top 5 TFs with the largest losses.

We next investigated how the regulatory networks identified in our study compared to similar networks constructed from reference VAT transcriptomes in the general population available from the Genotype-Tissue Expression project (GTEx, version 8). We hypothesized that the number of gene-gene, gene-TF, TF-gene and TF-TF connections will be different in the obese and non-obese samples when compared to the population average represented by the GTEx samples. Indeed, a comparison of the gene-TF and TF-gene interactions between our dataset and GTEx showed increased connectivities for both obese and non-obese interactomes (indicated by the shift of data points off the diagonals, and towards the obese and non-obese axis), with the changes were more pronounced for obese samples (**Fig. 2A-D**). Conversely, we observed a reduced number of TF-TF connections in obese samples compared to GTEx, whereas the number of TF-TF interactions between GTEx and the non-obese cohort were more similar (**Fig S2a-b**).

**Figure 2.**
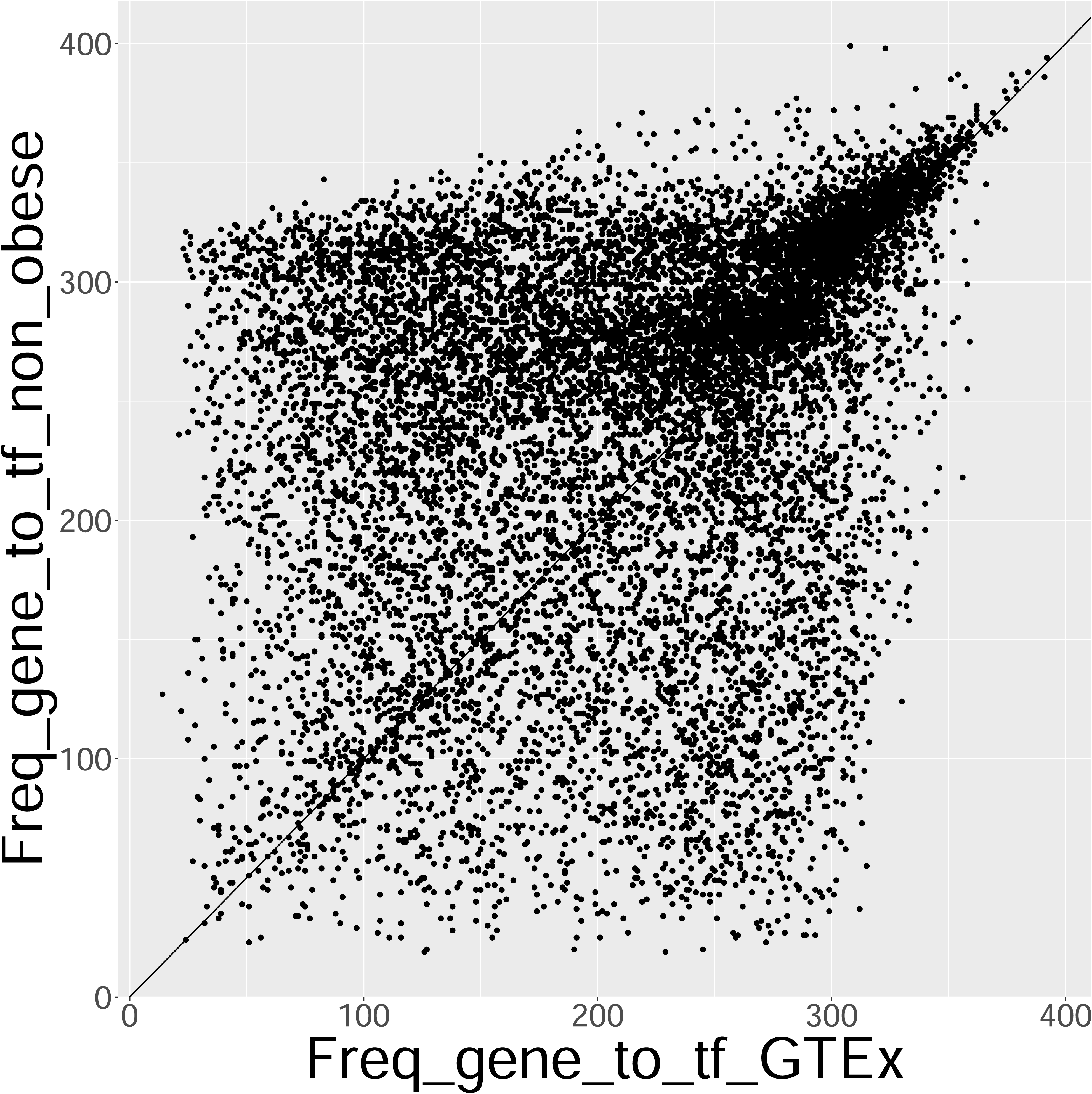
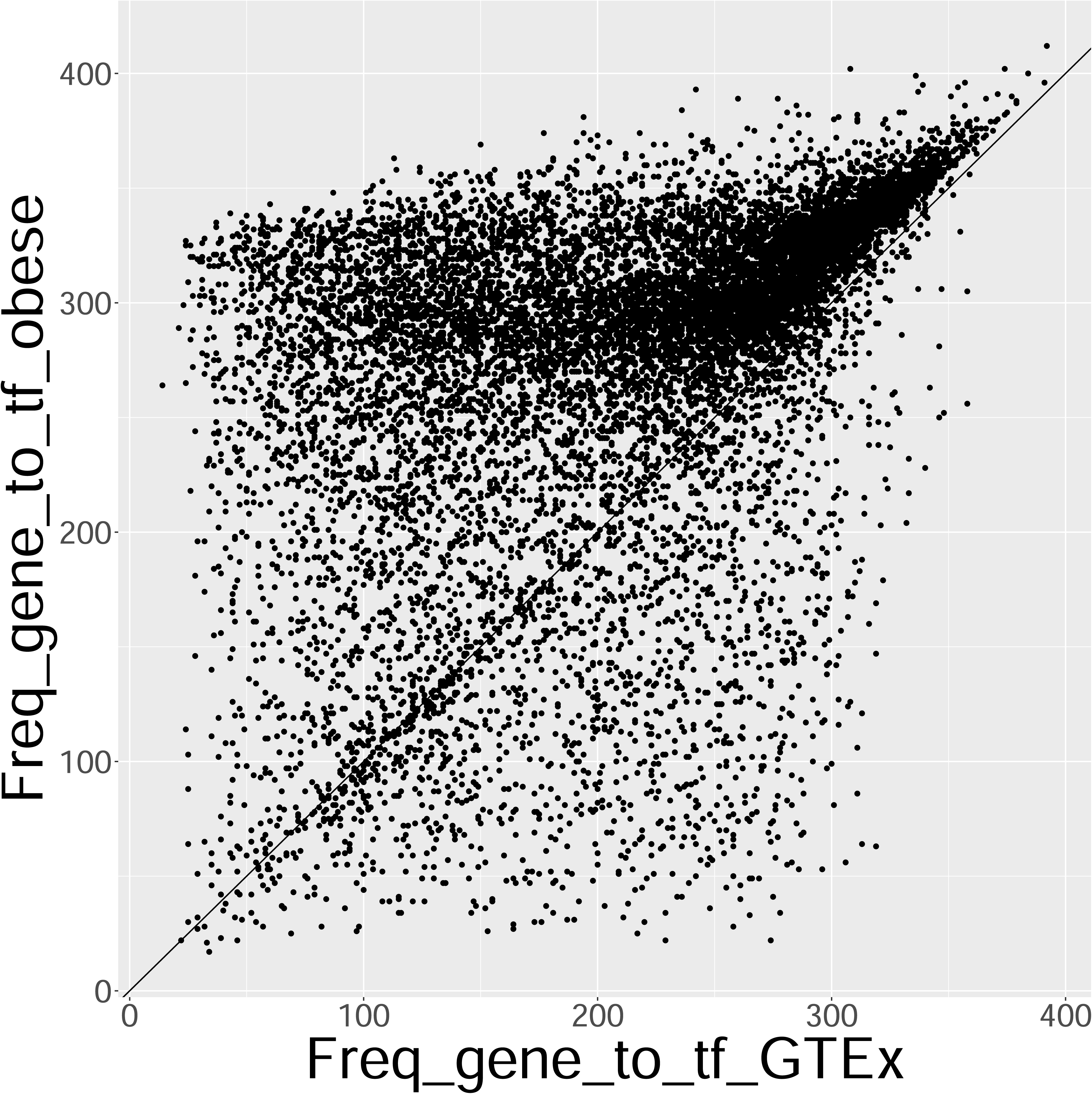
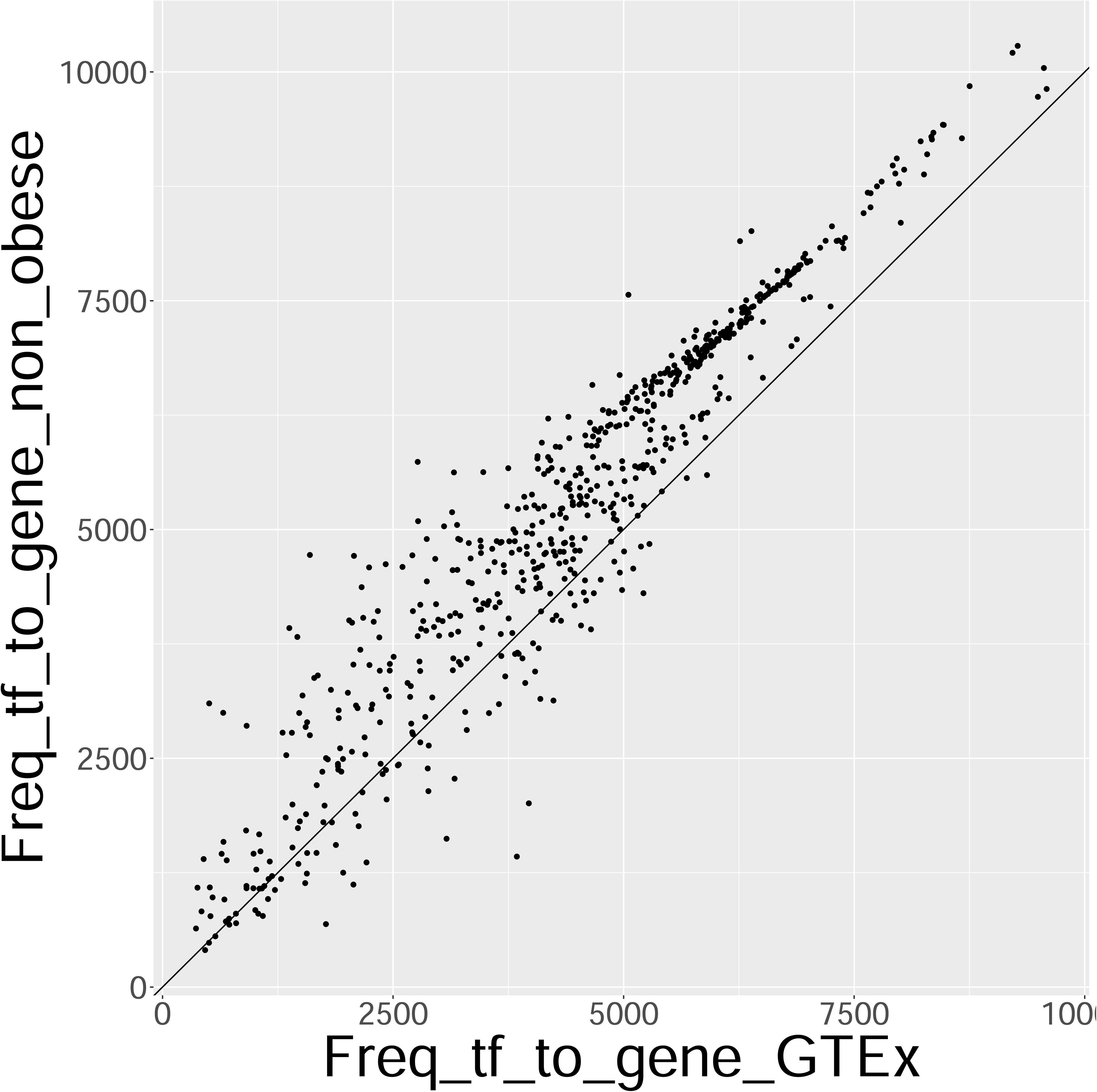
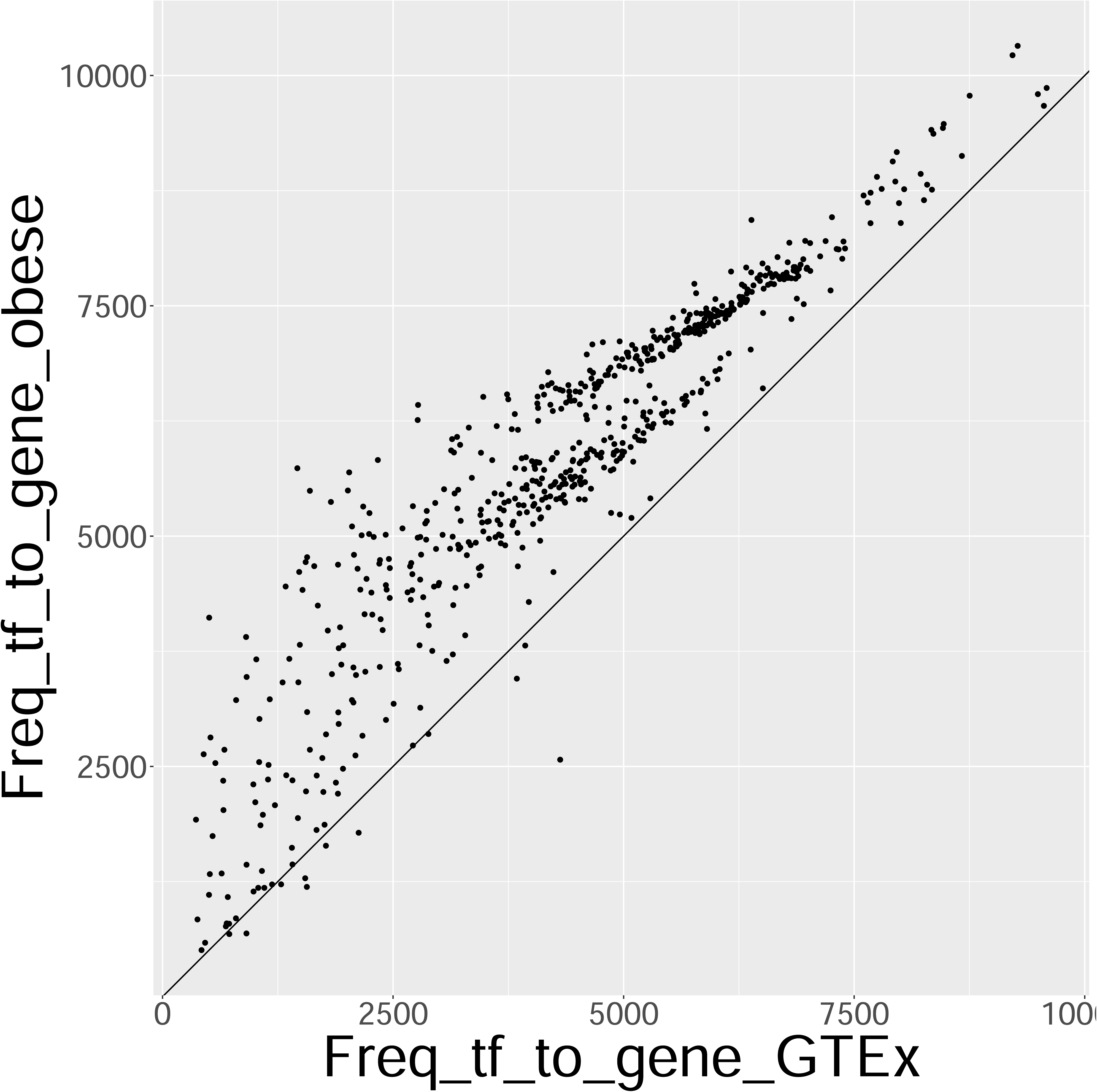
Comparison of obese and non-obese regulatory networks in current study to similar networks constructed from VAT transcriptomic data from GTEx. (A,B) Comparison of the number of associations between genes and transcription factors (gene-TF) in GTEx (x-axis) to non-obese **(A)** or obese **(B)** samples. Each dot represents a gene and the number represents the number of TFs the gene is predicted to interact with, according to PANDA. The diagonal represents an equal level of interaction between the dataset pairs. **(C,D)** Comparison of the number of associations between transcription factors and genes (TF-gene) in GTEx (x-axis) to non-obese **(C)** or obese **(D)** samples. Each dot represents a TF and the number represents the number of genes it is predicted to interact with, according to PANDA. The diagonal represents an equal level of interaction between the dataset pairs.

Pathway enrichment analysis on genes with over 1000 interaction changes revealed that genes with increased connectivities in obesity were enriched in inflammatory pathways like TNF-alpha and IL2/STAT5 signaling **(Fig. 3A)**. In contrast, genes with large connection losses in obesity were enriched in pathways like oxidative phosphorylation and adipogenesis (Fig. 3B).

**Figure 3.**
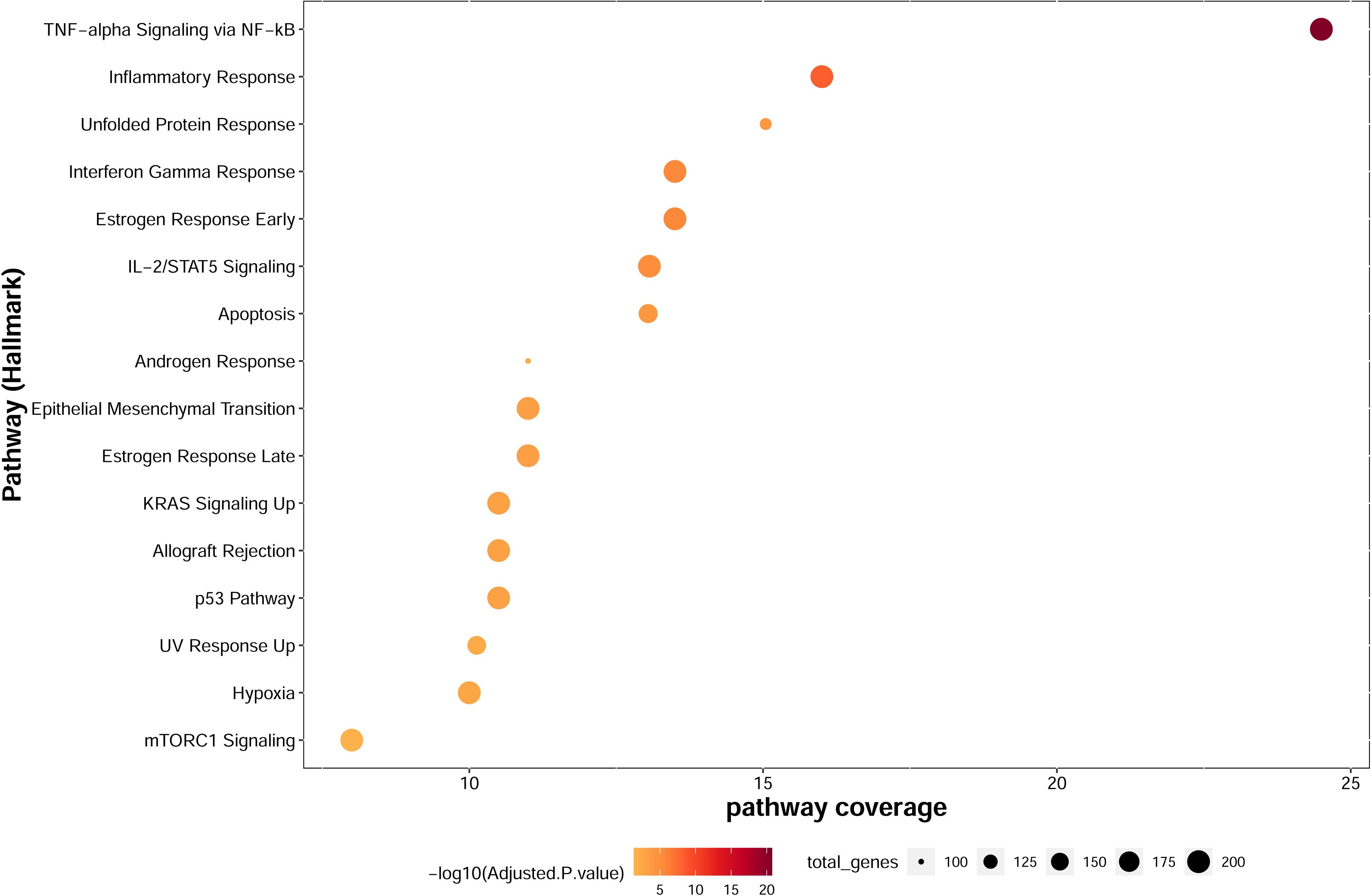
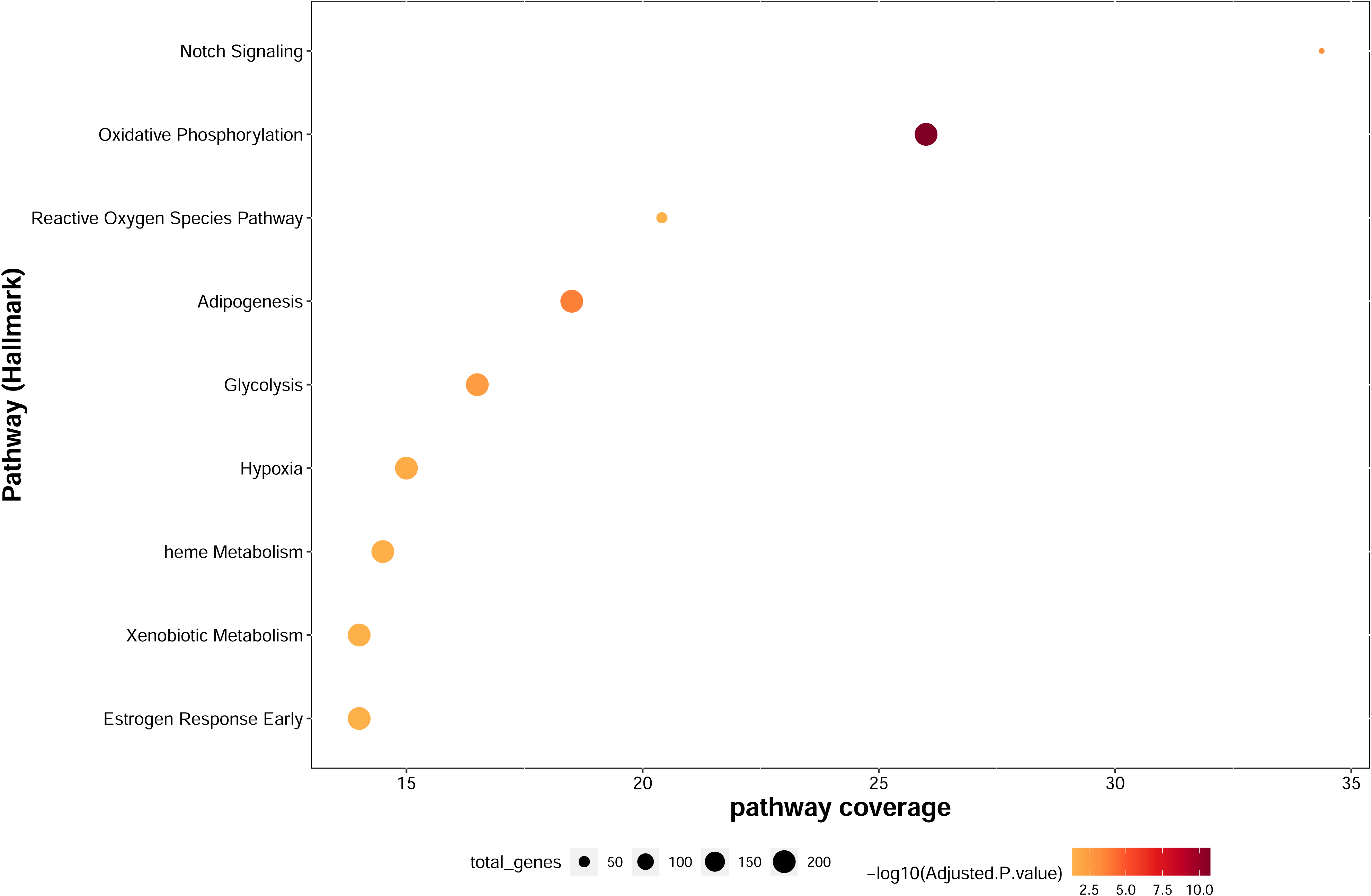
Enrichr-based pathway over-representation analysis on genes showing differences in connectivities (a change of >=1000 interactions) between obese and non-obese regulatory networks. (A, B) Over-represented Hallmark pathways in genes displaying a gain in connections in obesity **(A)**, or reduced connections in the obese state compared to the non-obese **(B)**. Pathways with FDR<0.05 and 10 or more genes in overlap are considered. The x-axis depicts pathway coverage (percent pathway in overlap with query gene list) and the y-axis shows the pathway names. Pathways are colored by –log(FDR) and sized by number of genes in pathway

### Altered network communities in non-obese and obese VAT

Using ALPACA, we identified 15 condition-specific gene modules (communities) in TF-gene regulatory networks, showing significant connectivity differences between obese and non-obese samples, with the largest community including 4670 genes and 270 TFs. Differences in modularity scores, depicted in **Fig. 4A**, quantified each gene’s and TF’s changing contributions to the community architecture in obese and non-obese samples. The top 5 differences in modularity scores were observed for TFs *PAX9, REL, PAX1, RELA* and *PAX2*, impacting modules 15 and 9. The *ARRDC2, NFIL3, EIF2B3, RASSF9* and *SEC61A1* genes displayed the top 5 modularity differences and influenced modules 15, 11, and 9. Pathway enrichment analysis of communities with >=80 members revealed obesity-associated enrichment in pathways involved in adipogenesis and TNF-alpha signaling (FDR<0.05) (**Fig. 4B**).

**Figure 4.**
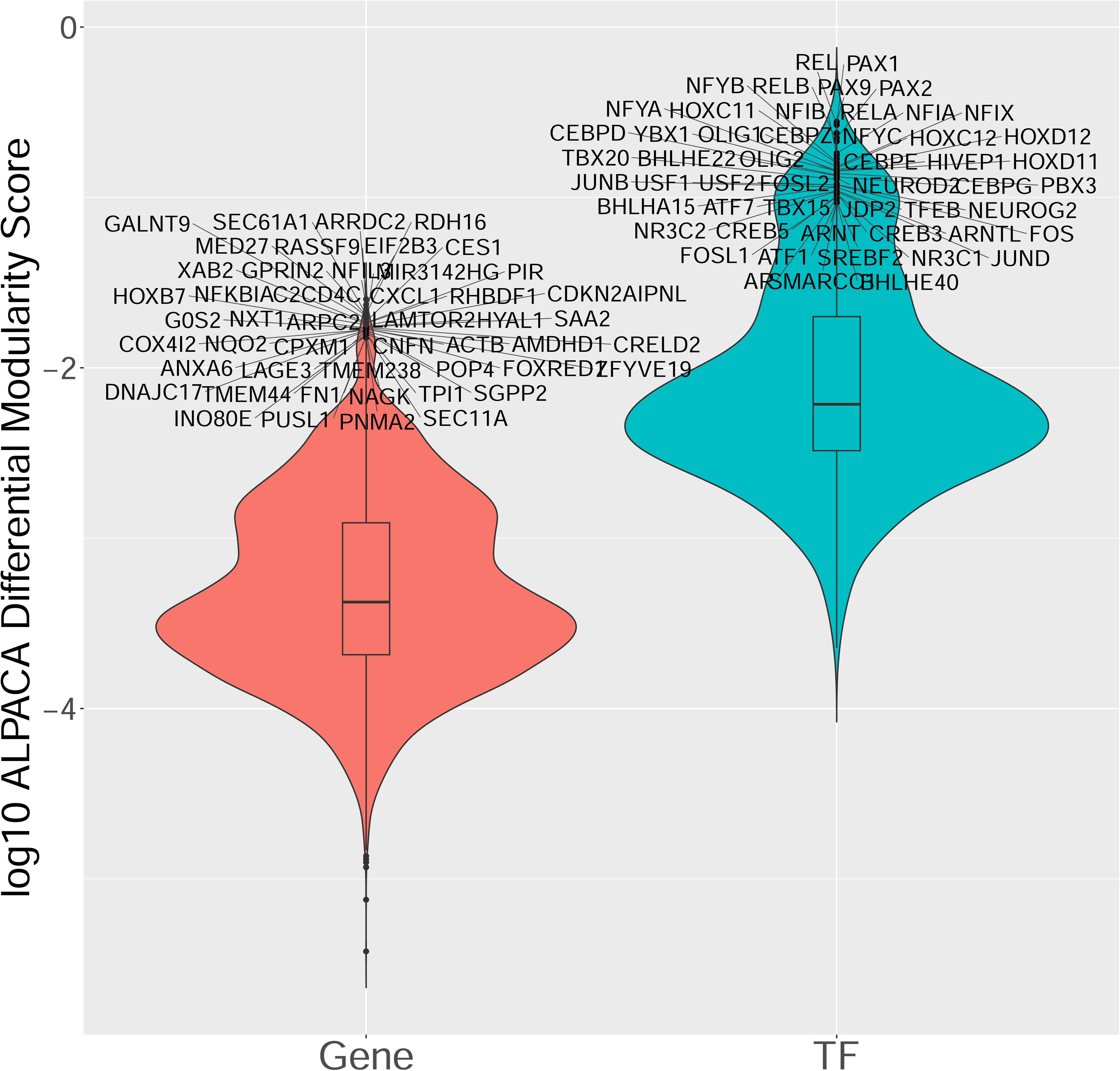
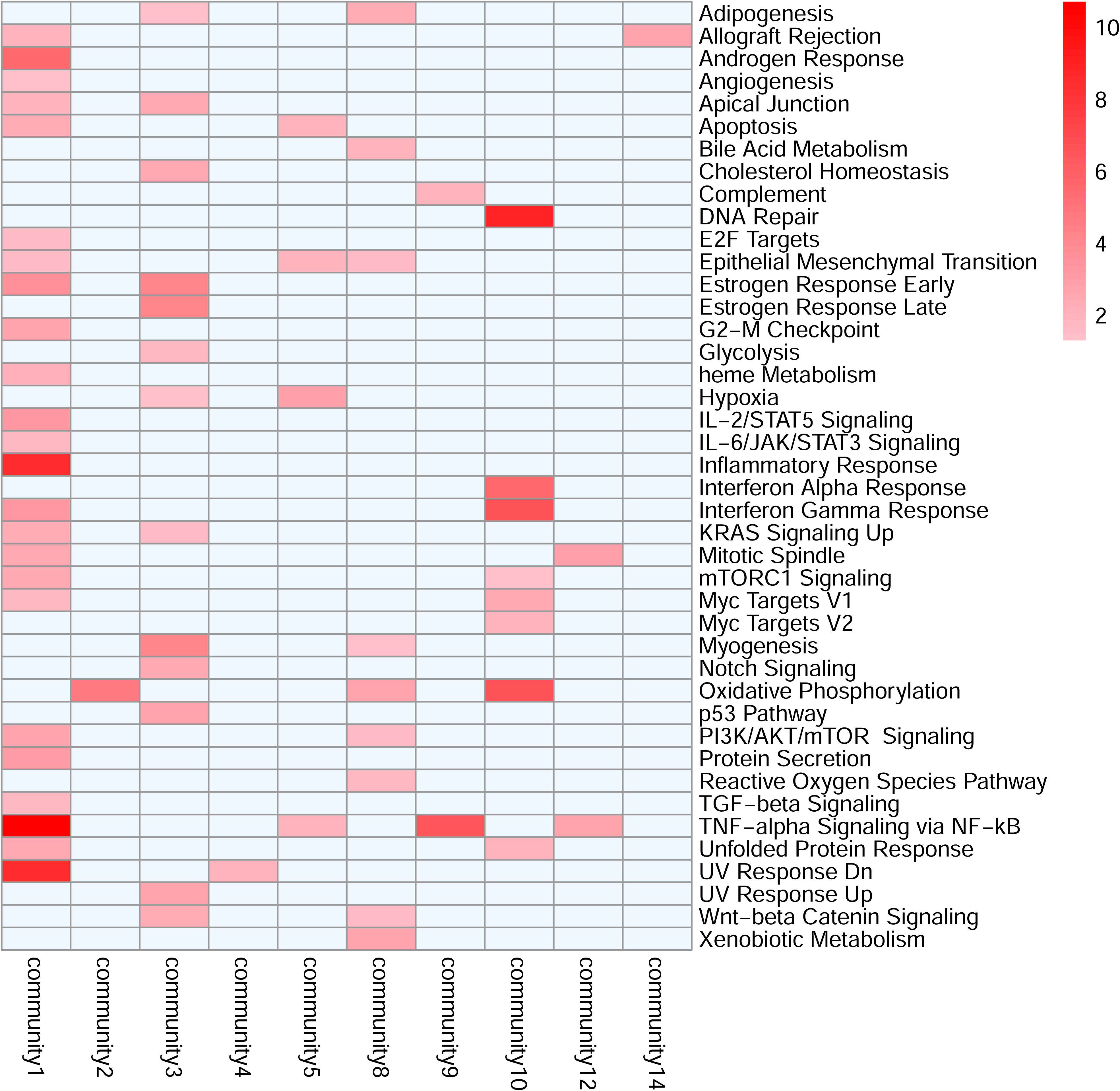
Analysis of network community structures in obese and non-obese VAT via ALPACA. (A) Violin plots showing the distribution of the difference in modularity scores (in log10 scale) for genes (red) and transcription factors (TF) between the regulatory networks in obese vs. non-obese samples. Genes and TFs showing the highest differences are annotated. **(B)** Enrichr-based pathway over-representation analysis for gene sets belonging to communities showing significantly different inter-node connectivities between obese and non-obese VAT regulatory networks (determined by ALPACA). Only communities with >80 members were considered for pathway enrichment analysis. Heatmap columns indicate community identifiers, while rows indicate Hallmark pathways. Heatmap is color coded from pink to red by –log10 of pathway enrichment adjusted p-value (only pathways with adjusted p-value <0.005 are shown). Light blue cells indicate absence of pathway data.

### Evidence for gene-disease association from external sources

We selected 80 candidate genes as described under Methods (**Supplementary Table S5**), and tested them for evidence of prior association to adiposity and related traits, based on literature (Pubmed), disease-gene databases (Disgenet), and genetic association studies (GWAS catalog). Pubmed searches yielded results for 61 of these genes, with 6 having ambiguous associations, and gene-adiposity associations ranging from 1-88. Disgenet searches implicated 70 genes as associated with cardiovascular diseases, with disease associations per gene ranging from 1-483 and DSI scores from 0.27-1 (**Fig. S4B and Table S7**). Notably, none of the candidate genes were linked to specific obesity phenotypes including *‘moderate obesity’*, *‘obesity’*, ‘*obesity, metabolically benign*’, ‘*obesity, morbid’*, ‘*obesity, visceral’*, ‘*obesity-associated insulin resitance*’ and *‘pediatric obesity’*. GWAS catalog lookup results encompassed gene-phenotype associations for 60 genes, with associations ranging from 1-120 traits (**Fig. S4C and Table S8**). This comprehensive analysis helped identify genes with minimal prior evidence of association with adiposity, marking them as novel candidates for functional testing.

### Gene ranking and selection for functional testing on adipocyte differentiation

We selected a set of 10 genes (including 4 transcription factors) for functional testing from an initial list of 80 genes, as described under Methods. Gene selection was performed heuristically and emphasized the relative novelty of the genes with respect to their known association to obesity (less than 15 GWAS-trait associations, less than 10 Pubmed obesity-related citations, and less than 50 Disgenet gene-disease associations). The higher threshold for Disgenet was required to accommodate transcription factors, which have a higher disease-association score compared to other gene categories, in our list of candidate genes.

### Functional testing in adipocytes

We conducted siRNA-based knockdowns in differentiating human adipocytes, and also quantified gene expression under untreated, mock treated and siRNA treated conditions via quantitative RT-PCR (**Figure 5A**). Distinct expression patterns were observed, e.g. *MUC20, SOX30,* and *OSBPL3* were induced early in differentiation, while *NFYC, FAM78B*, and *PPARG* increased later, with *PPARG* showing the highest induction (∼20-fold). *DDTL, DMRT2,* and *MEF2B* appeared to peak specifically on Day 6. We assessed the impact of gene knockdown on adipocyte differentiation by measuring lipid accumulation on Day 12 (**Figure 5B**). The level of adipocyte differentiation was essentially unchanged between the untransfected and mock-transfected cells (cells transfected with a scrambled siRNA construct), suggesting that observed changes in lipid accumulation are unlikely to arise from the procedures for nucleic acid transfection alone. PPARG knockdown most significantly reduced differentiation (∼40% compared to mock), while MEF2B, OSBPL3, SIRPB1, and SOX30 knockdowns led to over 25% reduction. Other genes showed 14-24% reductions (**Figure 5C**).

**Figure 5.**
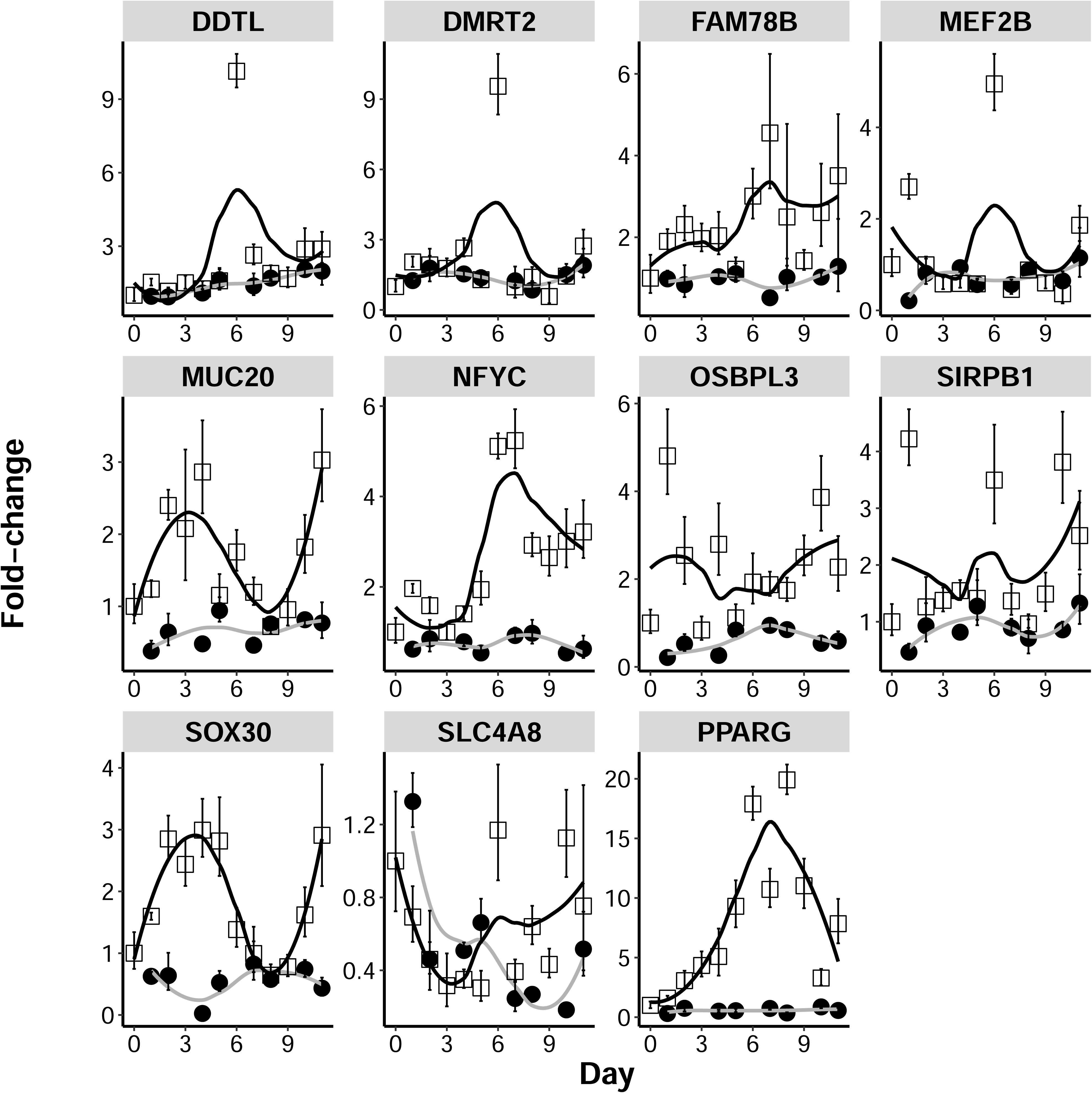
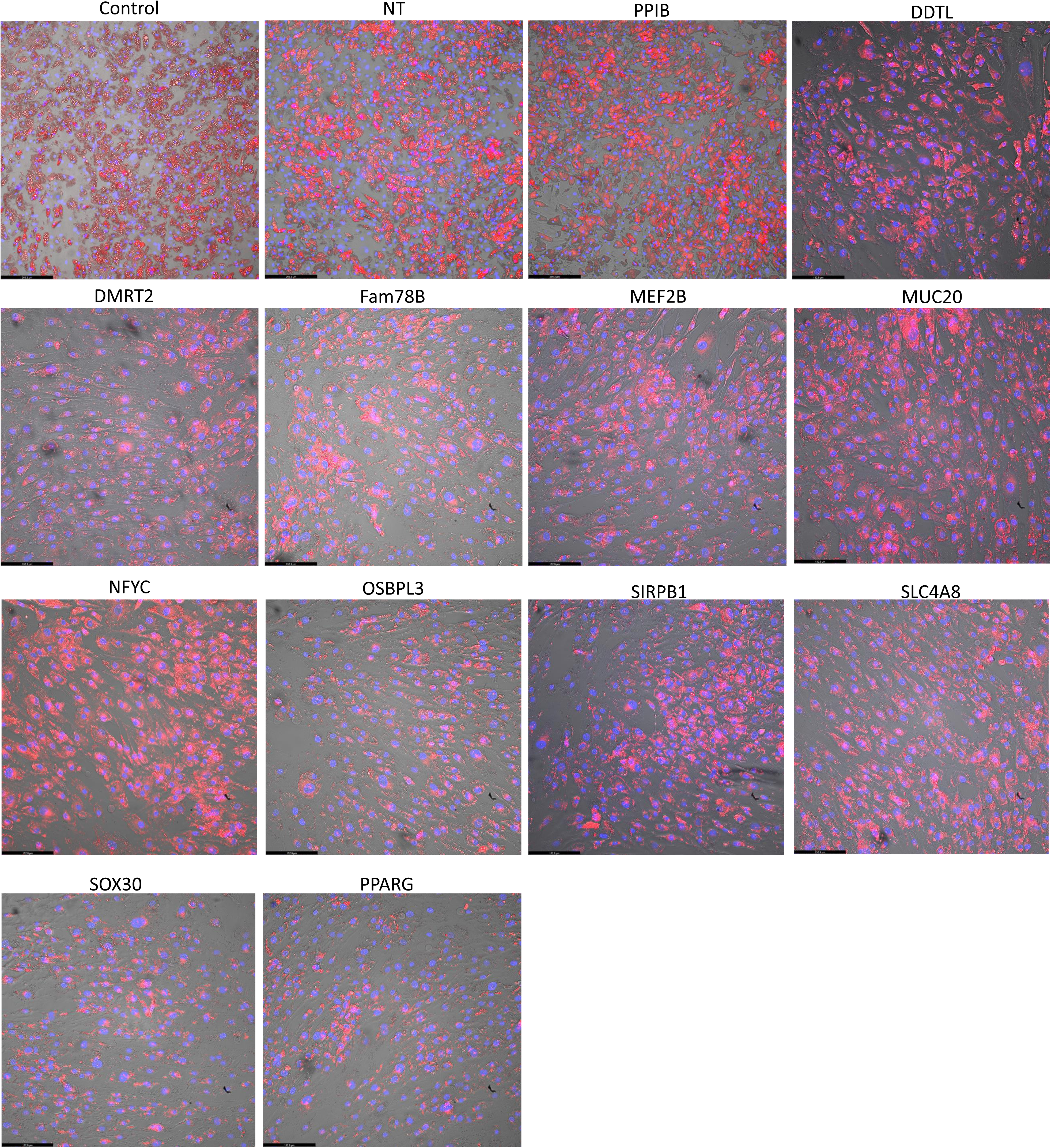
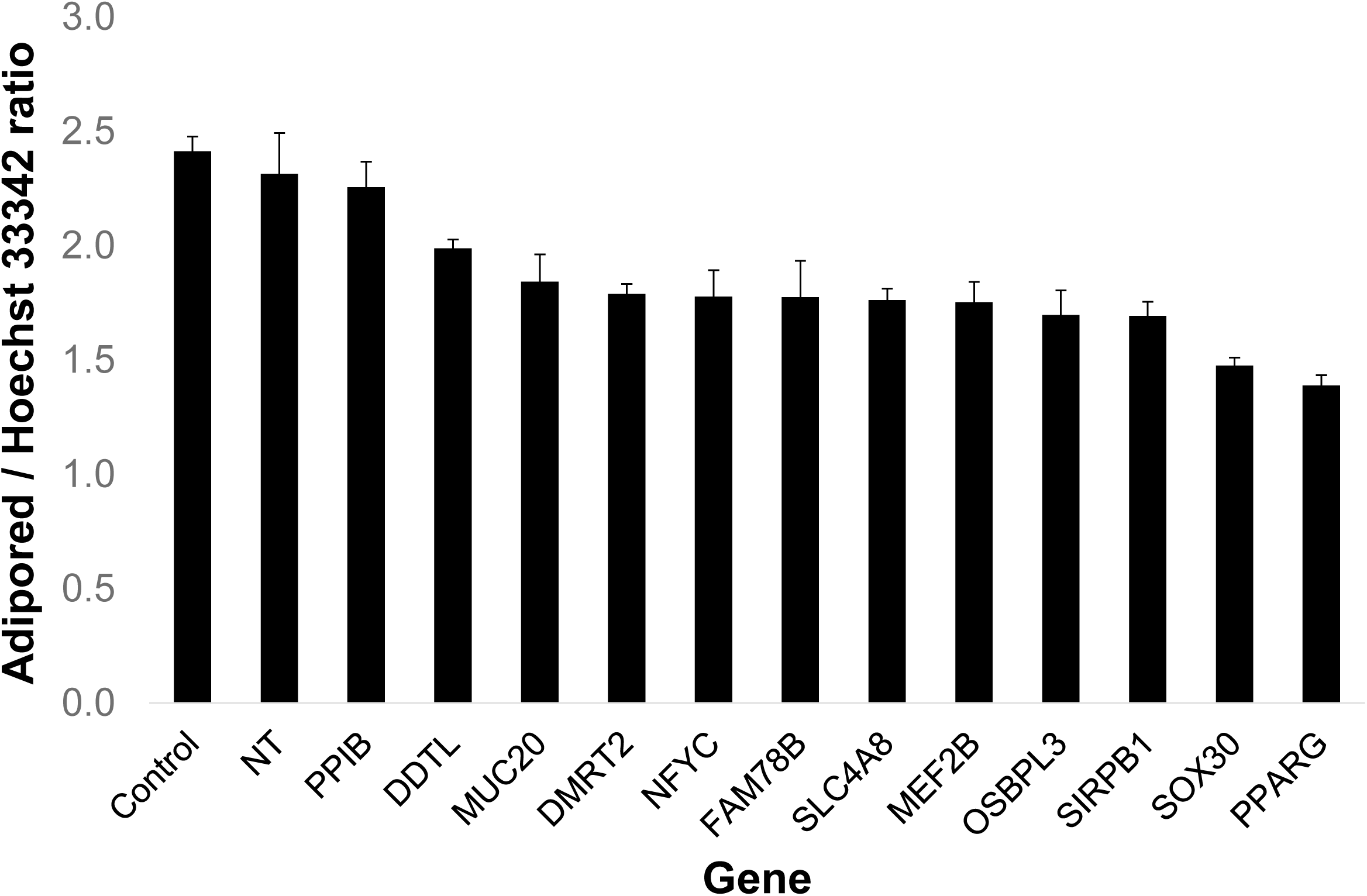
Functional screening of candidate genes via RNA interference. (A) Ten candidate genes, plus *PPARG*, were assayed for mRNA expression (open circles) and for the effects of siRNA mediated gene knockdown (closed circles) throughout the timecourse of differentiation of cultured human adipocytes. Gene expression in each day is expressed as fold-change compared to expression at Day 0. The extent of gene knockdown for a given day is expressed as the fold-change of expression compared to transfection with a non-targeting siRNA for the same day. The timecourse of differentiation is expressed along the x-axis and the fold-change in gene expression along the y-axis. **(B)** Estimation of lipid accumulation in adipocytes on Day 12 of differentiation via Adipored. Each subplot refers to the knockdown of the gene indicated at the top of the plot (NT=non-targeting siRNA, PPIB=loading control gene for qPCR). **(C)** Quantification of lipid content per cell (Adipored/Hoechst 33342 ratio) based on the staining of Day 12 adipocytes with Adipored (lipid accumulation) and Hoechst 33342 (cell nuclei). Experiments were performed in triplicates and results are expressed as means(±SD).

To investigate the *SOX30* gene-regulatory network in greater details, we constructed separate *SOX30* dependent sub-networks in obese and non-obese samples, based on OL-PANDA data as shown in **Figure 6A**. Most *SOX30* interacting genes (908) were common to both groups, with 121 and 94 genes unique to obese and non-obese samples, respectively. Pathway analysis demonstrated that *SOX30* interactors in obese samples were enriched in inflammation pathways, while those in non-obese samples were associated with oxidative phosphorylation. Additionally, genes interacting with SOX30 in both groups were linked to cell division and fatty acid metabolism (**Figure 6B**).

**Figure 6.**
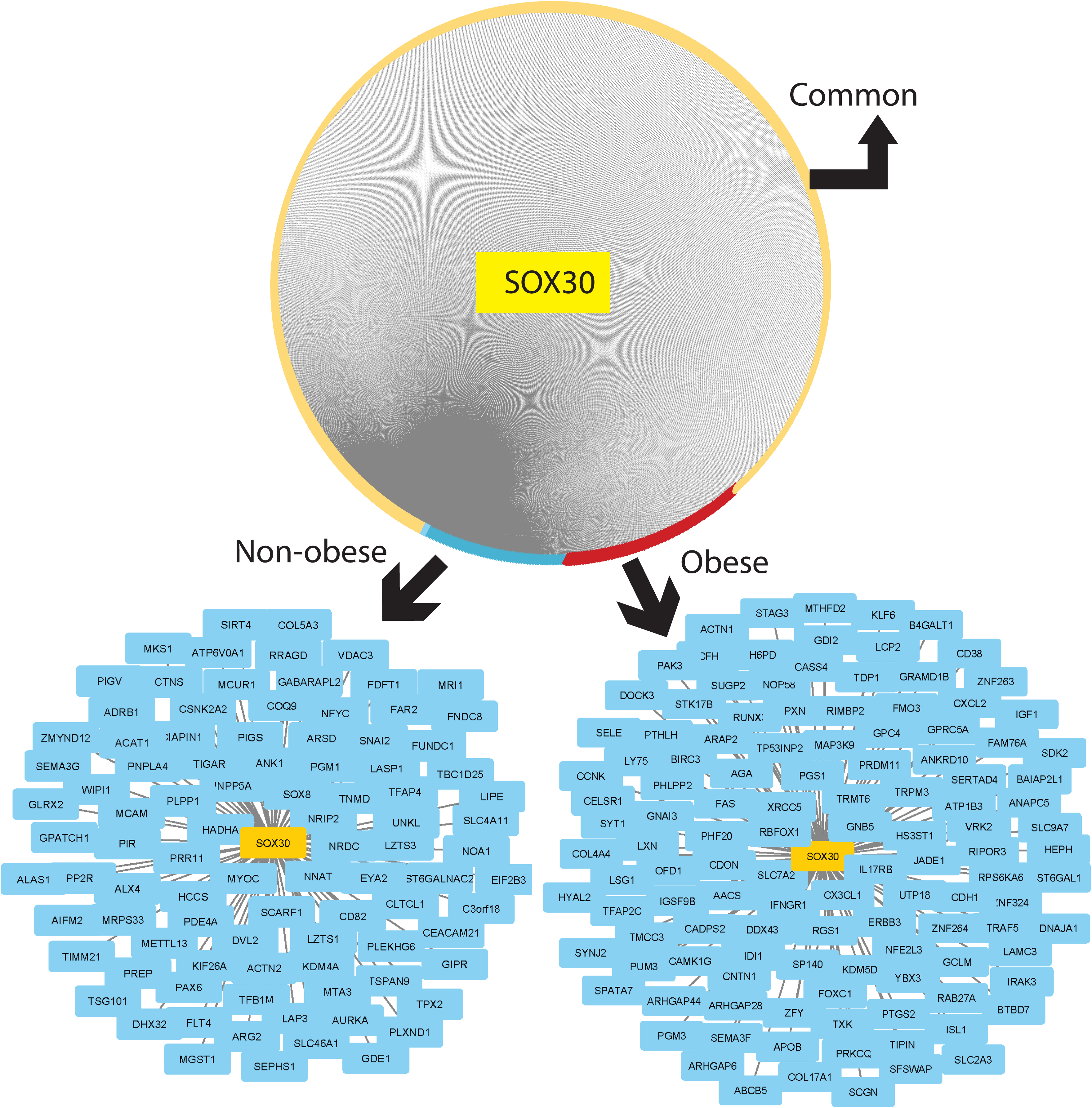
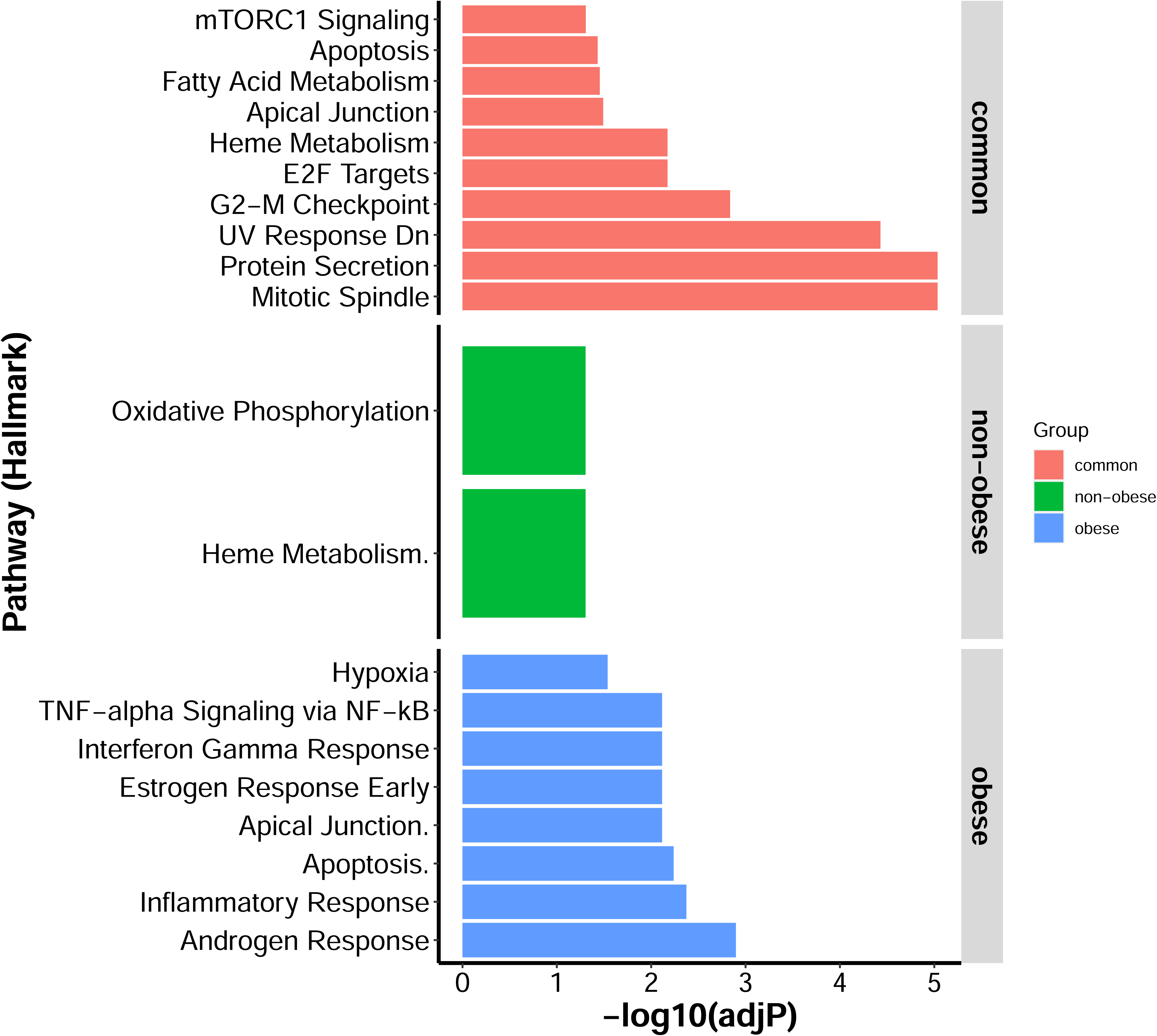
Gene regulatory network for SOX30 in obese and non-obese samples. SOX30 interacting genes were identified in PANDA. Network consists of SOX30 interacting genes in both obese and non-obese groups, (green part of circle), as well as genes that show interactions only in the obese (red) or non-obese (blue) groups. The obese and non-obese specific interactors are shown in the expanded network to the right. SOX30 in indicated in yellow and its interacting genes are shown in blue. **(B)** Pathway overrepresentation analysis of SOX30 interacting genes. Genes interacting with SOX30 either uniquely in obese or non-obese samples, or in common, were interrogated for enrichment of biological processes through the Hallmark pathway repository. The x-axis refers to the enrichment adjusted p-value (in –log10 scale) and the y-axis lists the Hallmark pathway names. Results are color-coded by the source of the genes.

## Discussion

In this paper, we employed a network-centric approach to understand gene transcription differences in visceral adipose tissue (VAT) from individuals with and without obesity. Our method combined gene co-expression and gene regulatory network analysis to interrogated key transcriptome-driven alterations in visceral obesity. Of the various methods available for gene co-expression (40), and differential gene co-expression analysis (41), we employed GeneNet for gene co-expression analysis, and two module-based approaches (PANDA and ALPACA), including transcription factor motif information and protein-protein interactions, to identify gene-regulatory networks and subnetworks (communities) in obese and non-obese VAT. We analyzed gene-gene and TF-gene networks through 9 different approaches and identified the top 10 genes from each analysis for investigation of prior association to obesity-related phenotypes based on genetic association, literature citation, and disease-gene database searches. Our goal was to identify genes not previously associated with multiple diseases or adiposity traits, considering them as potentially novel regulators of adipose tissue function with therapeutic potential. A subset of these genes was functionally tested through adipogenesis assays in human adipocytes. This constituted our overall strategy for employing network science for identifying and validating novel gene targets in VAT biology.

In the co-expression-based gene network, strong partial correlations were observed between *MUC20-MUC4, LINC01230-DMRT2*, and *AL049634.2-SIRPB1* genes. Of these, the long non-coding RNA *LINC01230* is proximally located to *DMRT2*, is regulated by *PPARG*, and is a modifier of endothelial function (42). The *AL049634.2-SIRPB1* association is also notable due to the association of *SIRPB1* to impulsivity and feeding behavior in obesity (43, 44). From PANDA-derived networks, both *SOX30* (SRY-box transcription factor 30) and *OSBPL3* (oxysterol-binding protein like 3) reduced lipid accumulation by >25% in our RNA-interference based functional screens of adipocyte differentiation. Not much is known regarding the role of *SOX30* in adiposity traits although *Sox30* knockout mouse displays altered fasting glucose levels (according to Mouse Genome Informatics (MGI) database, https://www.informatics.jax.org/). *OSBPL3* is primarily known for tumor growth regulation, but has recently been linked to fatty liver disease via *PPARG*-dependent mechanisms (45, 46). Interestingly, *Osbpl3* knockout mouse were found to display decreased body fat and increased lean body mass (according to MGI database search). In addition to individual genes, we also identified multiple network communities with altered connectivities between obese and non-obese VAT. These communities represented distinct biological functions related to inflammation, substrate and energy metabolism and cell division, as identified through pathway over-representation analysis.

Although our network-based approach identified several potentially novel gene candidates with functionally demonstrable effects on adipocyte lipid accumulation, some limitations of the study are now discussed. The generalizability of our results to VAT from other ethnicities remains unknown due to lack of comparable VAT transcriptomic data. Also, while the classification of obesity and non-obesity based on a BMI cutoff could be seen as arbitrary, we retained this classification to maintain sufficient sample numbers in each category and, consequently, maintain statistical power. We recognize that the functional screening employed in this study may not reflect the full scope of possible functions for the tested genes, but our choices were constrained by the non-availability of other tests with similar throughput. Finally, as our transcriptome analysis was performed at the adipose tissue level, the adipose-resident key cell type(s) where these genes function remains an area for future investigation.

In conclusion, our analysis identified distinct differences in gene regulatory networks in VAT from subjects with and without obesity, with further effects on network community structures. Functional testing of a subset of genes further identified potential novel regulators of adipocyte function. We hope that the approach presented here serves as a template for the application of network science for characterizing phenotypic states and identifying novel candidate gene effectors.

