## Supplemental Files for "Interactome Analysis of Visceral Adipose Tissue Elucidates Gene Regulatory Networks and Novel Gene Candidates in Obesity"

**CONTACT INFO:**

Sujoy Ghosh, Laboratory of Bioinformatics and Computational Biology,
Pennington Biomedical Research Center, Baton Rouge, LA 70808, USA.


### Supplementary File contents

**Supplementary Figure S1:** Relationship between degree and betweenness in gene coexpression network

**Supplementary Figure S2:** Comparison of TF-TF and gene-gene associations in PANDA-derived gene regulatory networks in obese and non-obese VAT to networks constructed from GTEx VAT

**Supplementary Figure S3:** Histograms of the distribution of candidate genes' associations observed in different query databases

**Supplementary Table S1:** QPCR primers used for candidate gene expression analysis in cultured adipocytes

**Supplementary Table S2:** List of binary interactions for coexpression network generated using GeneNet (p>0.8)

**Supplementary Table S3:** Network centrality measures for gene coexpression network constructed in Genenet

**Supplementary Table S4:** Output from PANDA analysis of differentially expressed genes showing connectivities to genes and TFs

**Supplementary Table S5:** List of 80 candidate genes, their ranks and the analysis datasets

**Supplementary Table S6:** Results from Pubmed analysis of 80 candidate genes

**Supplementary Table S7:** Results of Disgenet analysis of 80 candidate genes

**Supplementary Table S8:** Results from GWAS Catalog analysis of 80 candidate genes

Supplementary Figure S1. Relationship between degree and betweenness in gene coexpression network. A partial-correlation based gene coexpression network was constructed via GeneNet based on genes differentially expressed between obese and non-obese VAT samples. The degree and betweenness network centrality measures were computed via Centiscape. Plot shows the general positive relationship between degree (x-axis) and betweenness (y-axis) scores for network genes.

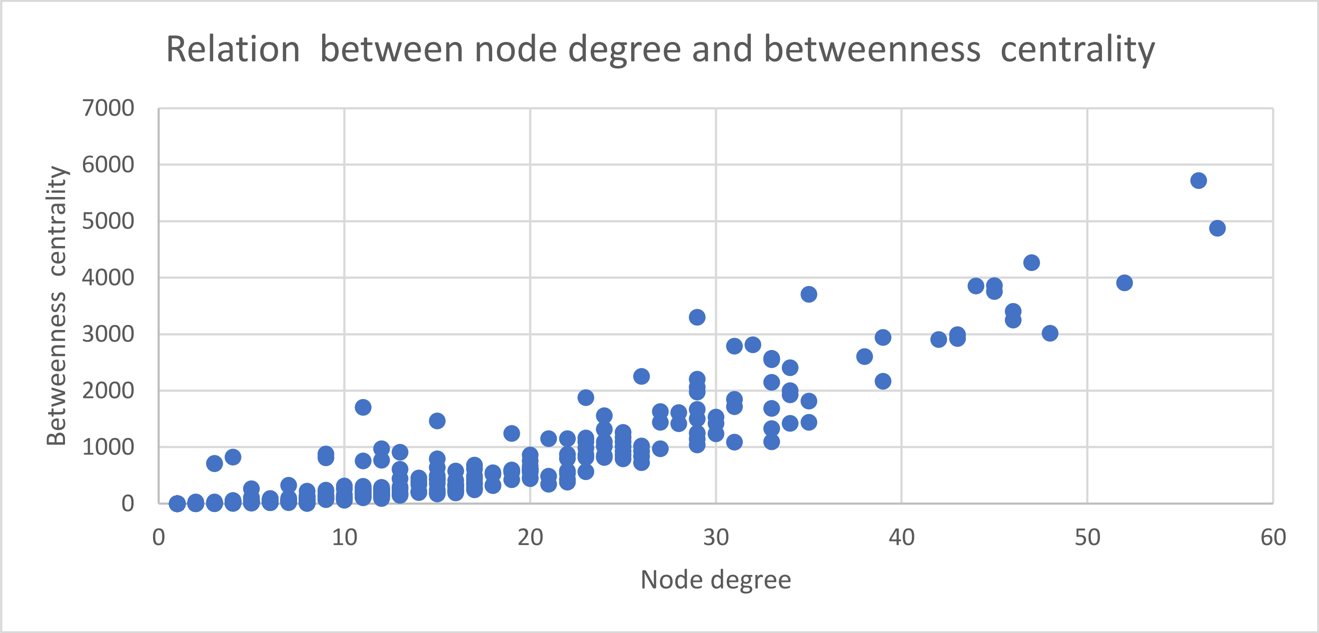

Supplementary Figure S2. Comparison of TF-TF and gene-gene associations in PANDA-derived gene regulatory networks in obese and non-obese VAT to networks constructed from GTEx VAT. (A) comparison of the number of associations in the TF-TF networks in non-obese VAT vs. TF-TF network in GTEx. (B) comparison between TF-TF network in obese VAT vs. GTEx. (C) comparison of the number of gene-gene interactions in non-obese VAT vs. gene-gene interactions in GTEx. (D) comparison of gene-gene interactions in obese VAT vs. those in GTEx.

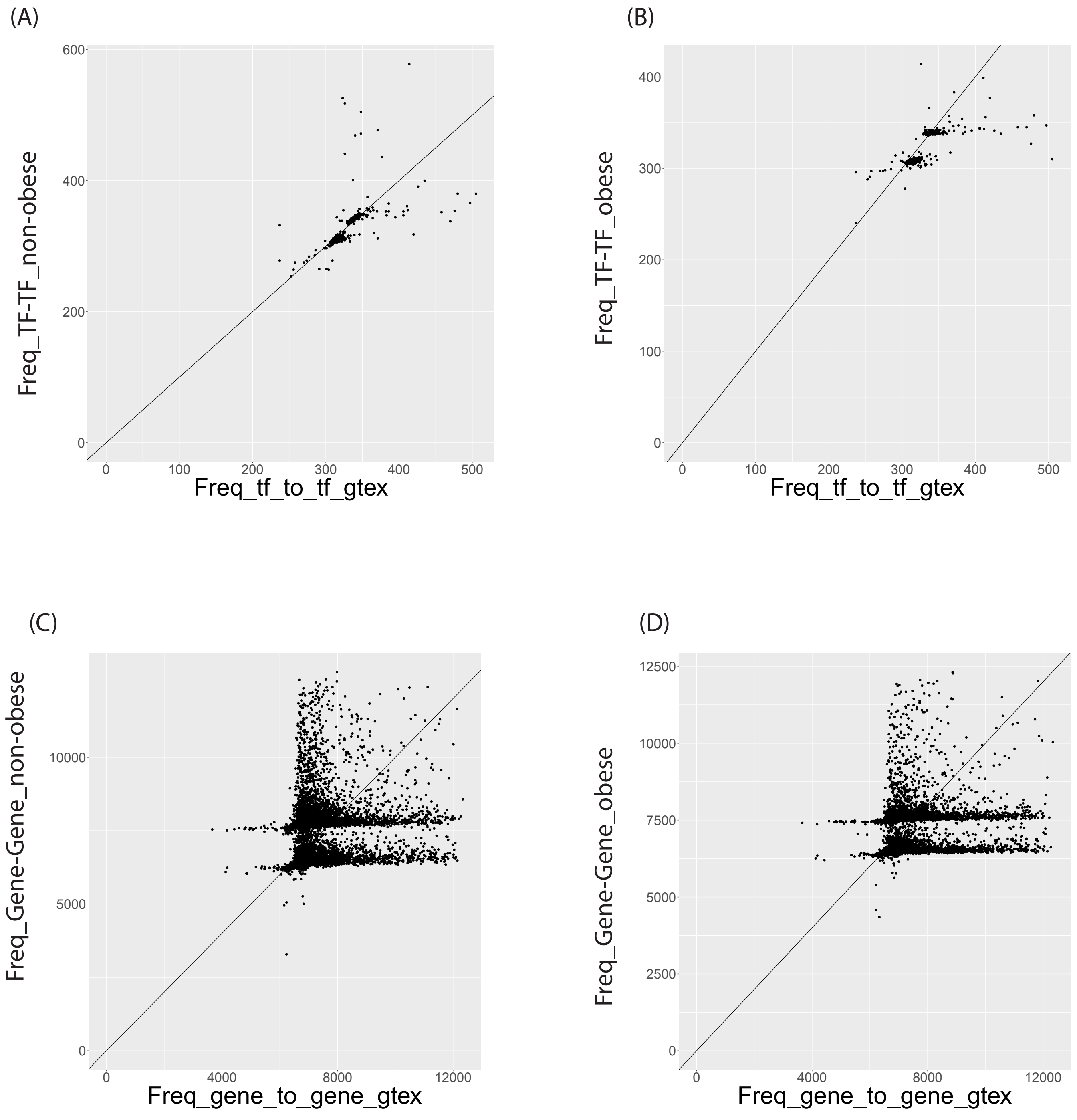

Supplementary Figure S3. Histograms of the distribution of candidate genes' associations observed in different query databases. (A) histogram of 80 candidate gene associations to 'obesity'-related publications in Pubmed. (B) histogram of candidate gene associations to diseases based on Disgenet database searches. (C) histogram of candidate gene associations to traits in GWAS catalog. Query details for all 3 searches have been described under Methods.

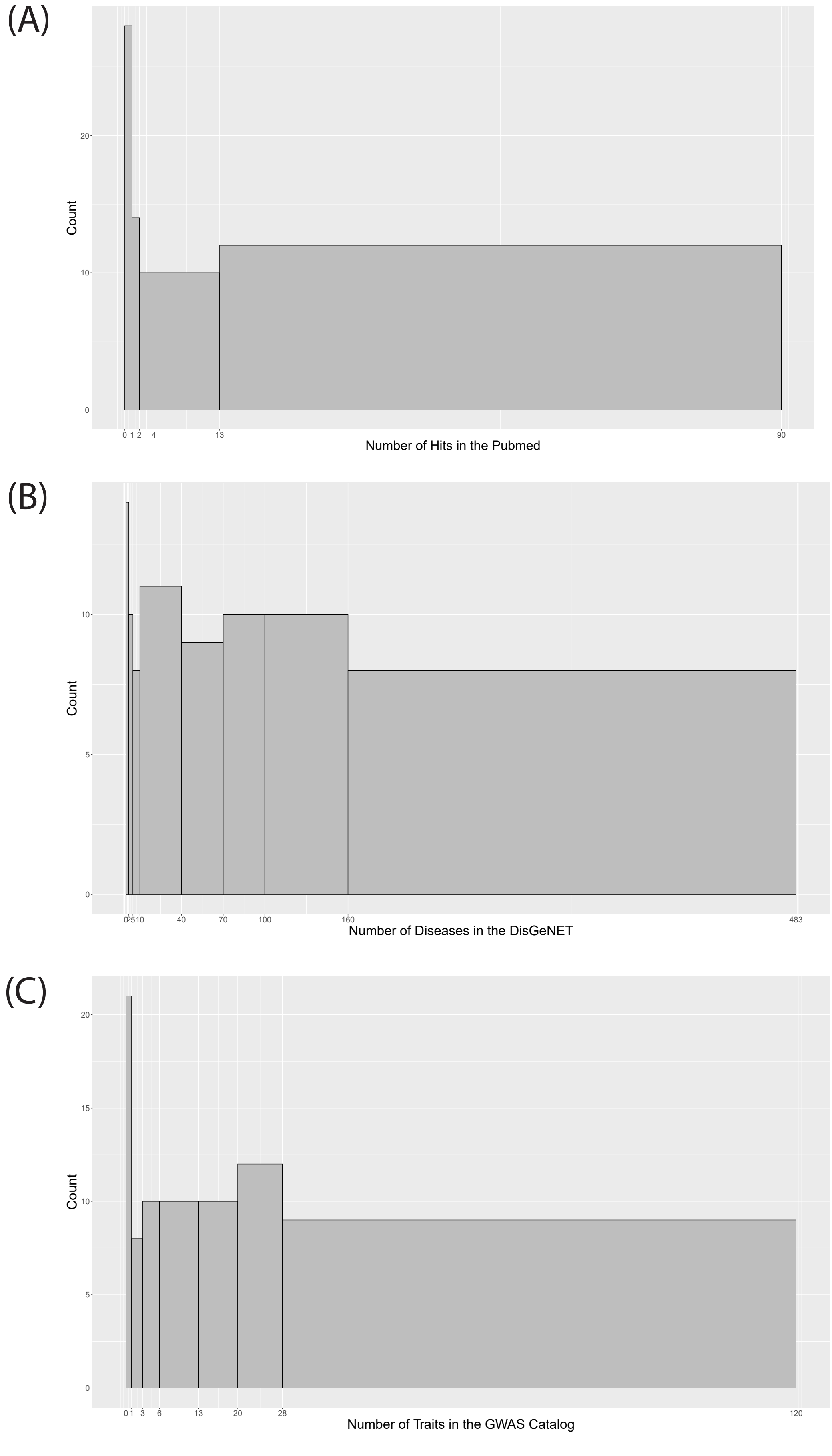

Table S1. QPCR primers used for candidate gene expression analysis in cultured adipocytes

| No | Species | Gene | Forward_primer | Reverse_primer |
|---|---|---|---|---|
| 1 | HU | CEBPG | 5' ACTCCAGGGGTGAACGGAAT 3' | 5' CATGGGCGAACTCTTTTTGCT 3' |
| 2 | HU | DDTL | 5' CATCTCCTCCATCGGCGTAGTG 3' | 5' GGCTGGTGGATAAGACCGTAGG 3' |
| 3 | HU | DMRT2 | 5' TTTAGAAGGCTATCGCCCCAT 3' | 5' TCCAGCATAATGTTCTCCAACTC 3' |
| 4 | HU | FAM78B | 5' CGCGAGAACATCGTGGTGTA 3' | 5' GGTCGCTGTAGGTGTTGAAGA 3' |
| 5 | HU | FGD2 | 5' CATCGGTGACGTGATCCAGAA 3' | 5' AGCCGCTCGCTCAAAGTTC 3' |
| 6 | HU | MEF2B | 5' ATGGACCGTGTGCTGCTGAAGT 3' | 5' TCCGAAACTTCTCTCCTGGCTC 3' |
| 7 | HU | MUC20 | 5' ATGACAACGGACGACACAGAA 3' | 5' TCAGCGTTTGAGTTTCCAGAG 3' |
| 8 | HU | NFYC | 5' GGAGGATTTGGTGGTACTAGCA 3' | 5' GCACTCGGAAGTCTTTCACTG 3' |
| 9 | HU | GLIS1 | 5' CGTCTCTGGTCACCTGTGTAA 3' | 5' CTCATGGCTGTCCGTCGAT 3' |
| 10 | HU | OSBPL3 | 5' TTGGTGTGTCCCAAAAATTGGT 3' | 5' TCCTGGGTGTAATTCATCTCCC 3' |
| 11 | HU | CEBPG | 5' ACTCCAGGGGTGAACGGAAT 3' | 5' CATGGGCGAACTCTTTTTGCT 3' |
| 12 | HU | SIRPB1 | 5' TGGCCCCACCTTCCTAGTC 3' | 5' AGCTCCTCTGAACCACTGGA 3' |
| 13 | HU | SLC4A8 | 5' CATCGTGACAGCAGAAGTCCAC 3' | 5' CATCGTGACAGCAGAAGTCCAC 3' |
| 14 | HU | SOX30 | 5' TGAGAGACTCGATGCAAGGC 3' | 5' GTGTGGAGCGTCAAAGGGA 3' |
| 15 | HU | PPARG | 5' GGGATCAGCTCCGTGGATCT 3' | 5' TGCACTTTGGTACTCTTGAAGTT 3' |
| 16 | HU | PPIB | 5' AAGTCACCGTCAAGGTGTATTTT 3' | 5' TGCTGTTTTTGTAGCCAAATCCT 3' |

Table S2. List of binary interactions for network generated using GeneNet (p>0.8)

| Nodes | Partial Correlation |
|---|---|
| DPP10~PTCHD4 | -0.11997 |
| AL035446.1~ATP8A2 | -0.11613 |
| TF~TMIGD2 | -0.11289 |
| MUC4~HS3ST2 | -0.09921 |
| GABRD~PKP3 | -0.09809 |
| LINC01230~MAB21L1 | -0.09443 |
| IGHD~EXOC3L2 | -0.09418 |
| HSPA6~AL035446.1 | -0.09246 |
| RASL10B~IGSF1 | -0.09154 |
| STRC~CLDN9 | -0.08876 |
| MUC20~HS3ST2 | -0.08723 |
| TF~MLIP | -0.0862 |
| TRIM54~ABCC8 | -0.08582 |
| CYP2A6~PHACTR3 | -0.08488 |
| AC004233.3~SOX10 | -0.08484 |
| AC131097.2~ADAM28 | -0.08472 |
| IGFN1~STRC | -0.08459 |
| GABRD~TRIM54 | -0.08441 |
| MIR4458HG~IGHD | -0.08405 |
| AL358334.2~HS3ST2 | -0.08396 |
| SNHG9~CLDN9 | -0.08391 |
| HSPA6~GDF1 | -0.08332 |
| DMRT2~AL358334.2 | -0.08329 |
| CA3~PHACTR3 | -0.0824 |
| AC004233.3~GDF15 | -0.08147 |
| MLIP~PKP3 | -0.08085 |
| GABRD~ACTG2 | -0.08031 |
| SOWAHA~SGCA | -0.08021 |
| ADAM28~SLC4A8 | -0.07984 |
| MUC20~LY6K | -0.07937 |
| WFDC1~ANKRD24 | -0.0791 |
| KCNK3~GLYAT | -0.07855 |
| HEY2~LY6K | -0.07835 |
| CCL22~AC005696.4 | -0.07831 |
| PKP3~SGCA | -0.07775 |
| HSPA6~CERS1 | -0.07762 |
| GABRD~RCOR2 | -0.07758 |
| CHIT1~PKP3 | -0.0769 |
| DMRT2~MAB21L1 | -0.07643 |
| LINC01230~AL358334.2 | -0.07623 |
| FCGR1A~HMCN2 | -0.07559 |
| TRPA1~IGHD | -0.07548 |
| HSPA6~ADRB3 | -0.07492 |
| SPRR2F~AMPD3 | -0.07491 |
| TRPA1~MATN4 | -0.07475 |
| FCGR2C~IGHD | -0.0744 |

| | |
|---|---|
| AC110285.6~AL049634.2 | -0.07427 |
| LINC01094~HEY2 | -0.07419 |
| IGFN1~TREML1 | -0.07395 |
| FCGR1A~SPATA18 | -0.07383 |
| HSPA6~AP000785.2 | -0.07366 |
| LINC00968~IGHD | -0.07324 |
| TF~RANBP3L | -0.07314 |
| TRIM54~RP11-592B15.9 | -0.07293 |
| PTCHD4~PKP3 | -0.0729 |
| ADRB3~ABCC8 | -0.07285 |
| PRR7~AC005696.4 | -0.07276 |
| STRC~GDF15 | -0.07227 |
| ADRB3~GRIN2C | -0.07211 |
| GAS2L2~SGCA | -0.0721 |
| CSRP2~AC092720.1 | -0.07188 |
| IGFN1~POTEF | -0.07182 |
| DACT2~CECR2 | -0.07177 |
| LY6K~CRTAM | -0.07161 |
| GABRD~IGSF6 | -0.07159 |
| RASL11B~MIR4458HG | -0.07155 |
| PKP3~SOX10 | -0.07153 |
| IGFN1~SPATA18 | -0.07132 |
| C6~IGSF1 | -0.07132 |
| IGHD~ADGRE2 | -0.07118 |
| RP11-592B15.9~IGHD | -0.0711 |
| SLC46A2~SPX | -0.07096 |
| PTGDR2~AL049634.2 | -0.07088 |
| NEXN~CD69 | -0.07073 |
| SPRR2F~LINC01094 | -0.07048 |
| ADRB3~SLC46A2 | -0.0704 |
| SOWAHA~CCL22 | -0.07036 |
| PKP3~PHACTR3 | -0.07028 |
| THBS1~AC005696.4 | -0.06945 |
| CA3~ABCC8 | -0.06918 |
| MUC20~SPATA18 | -0.06917 |
| MUC4~SPATA18 | -0.06876 |
| RASL11B~AC004233.3 | -0.06874 |
| POTEF~PTCHD4 | -0.06855 |
| G0S2~ALOX15B | -0.06852 |
| TRIM55~OXT | -0.06849 |
| PRDM1~AC110285.6 | -0.06841 |
| GABRD~LUCAT1 | -0.06838 |
| AC131097.2~TENM4 | -0.06825 |
| DHRS9~SOX10 | -0.06818 |
| LY6K~FOS | -0.06802 |
| ADAM28~HMCN2 | -0.06793 |
| CHIT1~TGFBR3L | -0.06764 |
| CLDN9~CYP2A6 | -0.06757 |

| | |
|---|---|
| HOXA9~NRTN | -0.06752 |
| CCL18~PHACTR3 | -0.06737 |
| CHIT1~LINC00942 | -0.06731 |
| CD28~TNFAIP3 | -0.06725 |
| FCGR1A~SOX10 | -0.06708 |
| GAS2L2~IGSF1 | -0.06699 |
| LY6K~ANKRD24 | -0.06694 |
| AL358334.2~IGHD | -0.06694 |
| MUC4~HOXA9 | -0.0669 |
| CCL22~SOX10 | -0.06647 |
| MLIP~SOX10 | -0.06637 |
| MLIP~TWIST1 | -0.06626 |
| TNFAIP3~SGCA | -0.06624 |
| HMCN2~TCF15 | -0.06618 |
| LRRC73~ATP8A2 | -0.06616 |
| MTRNR2L12~CD69 | -0.06612 |
| FCGR1A~AL035446.1 | -0.06607 |
| ADRB3~AC005696.4 | -0.06607 |
| MYCL~PKP3 | -0.06599 |
| SPATA18~RCOR2 | -0.06595 |
| ADRB3~TENM4 | -0.06585 |
| SPX~GFAP | -0.06578 |
| DPP10~ATP8A2 | -0.06577 |
| AC005696.4~CD300E | -0.06562 |
| TMIGD3~SGCA | -0.06555 |
| ADRB3~LINC01230 | -0.06546 |
| DPP10~AC005696.4 | -0.06533 |
| GABRD~ATP8A2 | -0.06528 |
| FCGR2C~DMRT2 | -0.06527 |
| CTLA4~PHACTR3 | -0.06527 |
| ADRB3~OLR1 | -0.06524 |
| LINC01956~LY6K | -0.06522 |
| MLIP~AC107976.1 | -0.0649 |
| KCNA2~C20orf204 | -0.06487 |
| MYCL~AC131097.2 | -0.06479 |
| TREML1~CLDN9 | -0.06476 |
| IGHD~SNHG9 | -0.06476 |
| TIGAR~CYP2A6 | -0.06473 |
| HTRA3~COL9A1 | -0.06471 |
| FGF22~IGSF1 | -0.06469 |
| LINC01770~DPP10 | -0.06468 |
| POTEF~CD69 | -0.06466 |
| LY6K~PKP3 | -0.06452 |
| KCNA2~GAS2L2 | -0.06441 |
| DPP10~APLNR | -0.06431 |
| ALPK3~IGSF1 | -0.06431 |
| ID4~FHOD3 | -0.06422 |
| FOSL2~LINC01956 | -0.0641 |

| | |
|---|---|
| TFCP2L1~TMIGD2 | -0.06403 |
| SPX~CD300E | -0.06379 |
| TUBB2A~OXT | -0.06374 |
| LINC01230~PHACTR3 | -0.06368 |
| FCN2~RCOR2 | -0.06363 |
| GAPT~LINC00942 | -0.06359 |
| STRC~AL049634.2 | -0.06355 |
| RASL11B~FOS | -0.0633 |
| CYR61~CLDN9 | -0.06324 |
| CRTAM~STRC | -0.06298 |
| FCGR1A~MUC4 | -0.06294 |
| HSPA6~NDRG4 | -0.06293 |
| FCGR1A~FHOD3 | -0.06292 |
| ELFN1~GAS2L2 | -0.06289 |
| NRIP3~F2RL3 | -0.06288 |
| IGFN1~GALNT6 | -0.06282 |
| IGHD~MATN4 | -0.06265 |
| MIR4458HG~FAM181B | -0.06242 |
| FCGR2C~LINC01230 | -0.06237 |
| OGDHL~EXOC3L2 | -0.06228 |
| NRIP3~PHACTR3 | -0.06215 |
| TRIM54~LINC00942 | -0.06196 |
| PDIA2~HS3ST2 | -0.06167 |
| SPRR2F~CA3 | -0.06157 |
| PRR7~LINC00942 | -0.06155 |
| TUBB2A~MAB21L1 | -0.06138 |
| TNFAIP3~AC010997.5 | -0.06138 |
| ALOX5AP~SGCA | -0.06136 |
| FCN2~CRTAM | -0.06134 |
| IL1RN~STRC | -0.06128 |
| ACTG2~MUC20 | -0.06126 |
| FCGR2C~CHIT1 | -0.06124 |
| CD69~IGHD | -0.06118 |
| ACTG2~ANKRD24 | -0.06105 |
| HSPA6~PTCHD4 | -0.06103 |
| PEBP4~GLYAT | -0.061 |
| CD69~ANKRD24 | -0.06093 |
| SPATA18~ADGRE2 | -0.06091 |
| GABRD~RORB | -0.06086 |
| CD52~CLDN9 | -0.06084 |
| TFCP2L1~LINC01230 | -0.06079 |
| MIR4458HG~CD69 | -0.06075 |
| AC131097.2~GFAP | -0.06071 |
| GAS2L2~GDF15 | -0.06064 |
| DPP10~TWIST1 | -0.06051 |
| HOXA9~AC004233.3 | -0.06038 |
| SDS~TBX1 | -0.06038 |
| BDH1~IGSF1 | -0.06034 |

| | |
|---|---|
| FCGR1A~AC005696.4 | -0.06031 |
| FKBP5~RCOR2 | -0.06029 |
| LINC01956~CD69 | -0.0602 |
| AL358334.2~LILRB3 | -0.06013 |
| MUC20~LUCAT1 | -0.06012 |
| IGFN1~FHOD3 | -0.06005 |
| TRIM54~AC092720.1 | -0.06002 |
| ATP8A2~RASL10B | -0.05974 |
| SLAMF8~PKP3 | -0.05947 |
| SPRR2F~SOWAHA | -0.05938 |
| AC004233.3~GRIN2C | -0.05938 |
| HTRA3~HS3ST2 | -0.05925 |
| RANBP3L~SGCA | -0.05909 |
| HOXA9~TGFBR3L | -0.05908 |
| SOWAHA~AL035446.1 | -0.05904 |
| NRIP3~CLDN9 | -0.05897 |
| SPRR2F~AC004233.3 | -0.05896 |
| GRIN2C~CNN1 | -0.05892 |
| IGHD~THBS1 | -0.0589 |
| MLIP~SUSD1 | -0.05866 |
| HEY2~F2RL3 | -0.05865 |
| ATP8A2~HS3ST2 | -0.05865 |
| TFCP2L1~TWIST1 | -0.05856 |
| IGFN1~OLR1 | -0.05853 |
| IGFN1~TF | -0.05848 |
| FAM181B~THBS1 | -0.05848 |
| C6~PSAT1 | -0.05845 |
| FCGR1A~MATN4 | -0.05844 |
| P2RY12~EXOC3L2 | -0.05842 |
| BDH1~RASSF10 | -0.05836 |
| MAB21L1~AL049634.2 | -0.05836 |
| AC131097.2~SMOC1 | -0.05821 |
| ADAM28~MS4A6E | -0.05816 |
| SPATA18~RASSF10 | -0.05806 |
| DPP10~TMIGD2 | -0.05803 |
| TWIST1~RASSF10 | -0.05803 |
| AC107976.1~AC004233.3 | -0.05802 |
| CD28~LINC00942 | -0.05799 |
| RASL11B~C5orf58 | -0.05799 |
| RASL11B~GAS2L2 | -0.05791 |
| OLR1~MATN4 | -0.05788 |
| IL1RN~F2RL3 | -0.05783 |
| HPCA~IGFN1 | -0.05773 |
| COL9A1~PKP3 | -0.05772 |
| POTEF~HOXA9 | -0.05771 |
| FCMR~HOXA9 | -0.05754 |
| TRIM54~ID4 | -0.05753 |
| NEXN~MAB21L1 | -0.05745 |

| | |
|---|---|
| ACTG2~MUC4 | -0.05745 |
| FAT2~EXOC3L2 | -0.05745 |
| C6~IGHD | -0.05726 |
| ANKRD24~PHACTR3 | -0.05725 |
| TFCP2L1~OXT | -0.05724 |
| LINC01770~ADGRL3 | -0.05721 |
| CTLA4~IGSF1 | -0.05721 |
| CXCR6~FOS | -0.05721 |
| RASL11B~AL035446.1 | -0.05719 |
| ACTG2~F2RL3 | -0.05718 |
| VNN1~ATP8A2 | -0.05718 |
| CSRNP1~ADAM28 | -0.05713 |
| IGFN1~AL035446.1 | -0.05702 |
| HSPA6~ABCC8 | -0.057 |
| HTRA3~CLDN9 | -0.05698 |
| AC092720.1~CCL18 | -0.05696 |
| LINC01956~LDHD | -0.05692 |
| ID4~AL358334.2 | -0.05689 |
| TWIST1~SGCA | -0.05682 |
| CLEC7A~CLDN9 | -0.05681 |
| LINC01703~HS3ST2 | -0.05679 |
| DCSTAMP~CYP2A6 | -0.05675 |
| PHACTR3~Z99774.1 | -0.05657 |
| GDF15~CYP2A6 | -0.05655 |
| HSPA6~LINC00942 | -0.05652 |
| AC107976.1~AC005696.4 | -0.05652 |
| AC131097.2~SLC46A2 | -0.05651 |
| GALNT6~ATP8A2 | -0.05649 |
| ARL4C~SPP1 | -0.05648 |
| LINC01956~MMP9 | -0.05642 |
| PROB1~CA3 | -0.05631 |
| AL358334.2~OXT | -0.0562 |
| SPATA18~ADRB3 | -0.05617 |
| UCHL1~MLIP | -0.05616 |
| LINC00942~CASS4 | -0.05614 |
| AKR1B15~CD69 | -0.05613 |
| TFCP2L1~SGCA | -0.05611 |
| HMCN2~OGDHL | -0.05611 |
| RASL11B~IGSF6 | -0.05589 |
| SUSD1~CLDN9 | -0.05588 |
| DMRT2~ART4 | -0.05584 |
| AC131097.2~PHLDA1 | -0.05583 |
| UBASH3B~IGHD | -0.05579 |
| CYP4B1~OSBPL3 | -0.05576 |
| SPRR2F~SOX10 | -0.05576 |
| RASL10B~ABCC3 | -0.05576 |
| HSPA6~PHLDA1 | -0.05571 |
| MLIP~HCN2 | -0.05568 |

| | |
|---|---|
| ADRB3~SUSD1 | -0.0556 |
| CYR61~MSC | -0.05559 |
| LINC01956~CA3 | -0.05557 |
| IGFN1~CD69 | -0.05556 |
| TWIST1~KCNB1 | -0.05545 |
| IGFN1~PKP3 | -0.05544 |
| LINC01956~TM4SF19 | -0.05543 |
| LUCAT1~MLIP | -0.05536 |
| RP11-592B15.9~PDIA2 | -0.05536 |
| ADAM28~GRIN2C | -0.05529 |
| GABRD~VNN1 | -0.0552 |
| HPCA~CA3 | -0.0552 |
| SUSD1~CYP2A6 | -0.05519 |
| CSRP2~GAS2L2 | -0.05516 |
| ID4~TRIM55 | -0.05513 |
| LUCAT1~AC005696.4 | -0.05512 |
| CRTAM~WFDC1 | -0.0551 |
| GABRD~ADGRL3 | -0.05508 |
| TMIGD3~CECR2 | -0.05497 |
| AC004233.3~F8A2 | -0.05493 |
| IGHD~MT-ND3 | -0.05491 |
| C20orf204~IGSF1 | -0.05489 |
| ID4~CLDN9 | -0.05474 |
| MIR4458HG~HS3ST2 | -0.05473 |
| TF~AC015688.5 | -0.05466 |
| FKBP5~MSC | -0.05457 |
| POTEF~FGF22 | -0.05453 |
| VNN1~F8A2 | -0.0545 |
| HCN2~CYP2A6 | -0.05443 |
| PEBP4~PDIA2 | -0.05442 |
| P2RY12~AL358334.2 | -0.05439 |
| CSRP2~MATN4 | -0.0543 |
| HTRA3~TRIM55 | -0.05424 |
| ART4~OXT | -0.05421 |
| SPATA18~AP000785.2 | -0.0542 |
| CHIT1~AC092720.1 | -0.05419 |
| TRIM55~CERS1 | -0.05419 |
| TRIM55~GDF1 | -0.05407 |
| SPRR2F~CHIT1 | -0.05404 |
| LINC01230~ADGRE2 | -0.05398 |
| CCR1~GAS2L2 | -0.05393 |
| RORB~HMCN2 | -0.05385 |
| SOWAHA~GAS2L2 | -0.0538 |
| FCGR2C~TREML1 | -0.05375 |
| SOWAHA~ALPK3 | -0.05372 |
| CA3~PTPRO | -0.05369 |
| SUSD1~MAB21L1 | -0.05363 |
| TMEM52~VNN1 | -0.05362 |

| | |
|---|---|
| SPATA18~TMEM88 | -0.05356 |
| SPRR2F~AC015688.5 | -0.0535 |
| GSDMB~IGSF1 | -0.0535 |
| FCGR2C~RASL11B | -0.05349 |
| TNFAIP3~PHACTR3 | -0.05347 |
| IGHD~SOX10 | -0.05339 |
| DHRS9~PDIA2 | -0.05335 |
| ITK~VNN1 | -0.05334 |
| SPRR2F~FOS | -0.05333 |
| PKP3~TOP2A | -0.05333 |
| MACROD1~PDIA2 | -0.05329 |
| ADRB3~CDKN2B | -0.05327 |
| LINC01230~GALNT6 | -0.05327 |
| ADORA3~CD300E | -0.05326 |
| SPRR2F~UCHL1 | -0.05316 |
| HMCN2~CRTAM | -0.05312 |
| TRIM55~SUSD1 | -0.05305 |
| POTEF~LRRC46 | -0.05293 |
| IL2RA~NRIP3 | -0.0529 |
| OGDHL~GLYAT | -0.0529 |
| HOXA9~Z99774.1 | -0.05289 |
| DUSP1~IGHD | -0.05288 |
| ATP8A2~CCL18 | -0.05286 |
| SPX~C20orf204 | -0.05285 |
| IGHD~SGCA | -0.05283 |
| FCGR2B~PTCHD4 | -0.05282 |
| RASL11B~SNHG9 | -0.05282 |
| RP11-592B15.9~HCN2 | -0.0528 |
| PTPRO~KCNB1 | -0.0528 |
| AC131097.2~GAS2L2 | -0.05276 |
| SPRR2F~TF | -0.05275 |
| GAPT~LINC01230 | -0.05273 |
| AC005696.4~TCF15 | -0.05272 |
| CNFN~AL049634.2 | -0.05272 |
| SPRR2F~TRIM54 | -0.05269 |
| LINC01956~AC004233.3 | -0.05266 |
| GJC3~CLDN9 | -0.05262 |
| IGFN1~AC131097.2 | -0.05255 |
| KCNA2~FCN2 | -0.05253 |
| SUSD1~AP000785.2 | -0.05252 |
| ABCC8~CXorf21 | -0.05251 |
| ATP8A2~IGHD | -0.05249 |
| AC010997.5~THBS1 | -0.05243 |
| GLYAT~NRTN | -0.05243 |
| NEXN~THBS1 | -0.05238 |
| CYP4B1~SPX | -0.05236 |
| RASSF10~IGHD | -0.05228 |
| TUBB2A~AC092720.1 | -0.05225 |

| | |
|---|---|
| ENHO~CECR2 | -0.05221 |
| AC010997.5~CYP2A6 | -0.05221 |
| DPP10~F2RL3 | -0.05219 |
| UBASH3B~CSRP2 | -0.05213 |
| LRRC73~SGCA | -0.05208 |
| SPRR2F~IGSF1 | -0.05207 |
| VNN1~CSRP2 | -0.05205 |
| MAB21L1~IGSF1 | -0.05203 |
| SPATA18~TBXA2R | -0.052 |
| VNN1~IGSF1 | -0.052 |
| BCAT1~SOX10 | -0.052 |
| COL9A1~MATN4 | -0.05189 |
| SPP1~AL035446.1 | -0.05187 |
| C6~NQO1 | -0.0518 |
| ATP8A2~AC092118.1 | -0.05179 |
| NEXN~MSC | -0.05178 |
| CKMT2~PTCHD4 | -0.05175 |
| PLA2G7~BIRC3 | -0.05172 |
| DACT2~WFDC1 | -0.0517 |
| AC131097.2~CLDN9 | -0.05169 |
| SGCA~AL049634.2 | -0.0516 |
| ADGRL3~IGHD | -0.05157 |
| ADORA3~CYP2A6 | -0.05156 |
| TFCP2L1~DHRS9 | -0.05145 |
| LINC00968~SLC4A8 | -0.05136 |
| SPRR2F~F8A2 | -0.05135 |
| ADORA3~UBASH3B | -0.05129 |
| SUSD1~ATP8A2 | -0.05124 |
| GPR171~AC005696.4 | -0.05123 |
| CYP4B1~AC010997.5 | -0.05122 |
| IGFN1~TUBB2A | -0.0512 |
| AL035446.1~CNN1 | -0.05118 |
| PKP3~CLEC4E | -0.05116 |
| TRIM54~FOS | -0.05113 |
| TENM4~IGHD | -0.05113 |
| CDKN2B~IGHD | -0.0511 |
| HMCN2~LIPA | -0.05108 |
| IGFN1~SGCA | -0.05105 |
| AP000785.2~SNHG9 | -0.05105 |
| CD300E~CECR2 | -0.05105 |
| FCGR2C~AC069368.1 | -0.05104 |
| PKP3~FMN1 | -0.05104 |
| HTRA3~UBASH3B | -0.05102 |
| AC107976.1~CYP2A6 | -0.05102 |
| ELFN1~CECR2 | -0.05099 |
| ADRB3~TGFBR3L | -0.05096 |
| TF~DCSTAMP | -0.05091 |
| TRPA1~DRD4 | -0.0509 |

| | |
|---|---|
| HEY2~KCNE1 | -0.05088 |
| PTGDR2~ART4 | -0.05084 |
| DHRS9~AC131097.2 | -0.05075 |
| AKR1C3~EXOC3L2 | -0.05074 |
| NRIP3~LINC00942 | -0.05069 |
| CECR2~IGSF1 | -0.05068 |
| MS4A6E~HRC | -0.05066 |
| RASSF10~CYP2A6 | -0.05062 |
| IGFN1~MSC | -0.05057 |
| AKR1C3~PKP3 | -0.05052 |
| KCNK3~TRIM55 | -0.05048 |
| BDH1~ID4 | -0.05048 |
| IGFN1~EXOC3L2 | -0.0504 |
| CARMN~PHLDA1 | -0.05039 |
| GABRD~NRIP3 | -0.0503 |
| CD52~CSRP2 | -0.05027 |
| CYP4B1~ARL4C | -0.05023 |
| THEMIS2~AC005696.4 | -0.05021 |
| HTRA3~Z99774.1 | -0.05019 |
| PEBP4~IGSF6 | -0.05018 |
| CD300E~PHACTR3 | -0.05018 |
| C5orf58~AL358334.2 | -0.05006 |
| MIR4458HG~TGFBR3L | -0.05004 |
| CSRP2~CCL22 | -0.05004 |
| CYR61~LY6K | -0.05003 |
| C6~SOX10 | -0.04996 |
| ATP8A2~FGF22 | -0.04996 |
| SPX~CLDN9 | -0.04995 |
| TRIM54~DMRT2 | -0.04993 |
| DPP10~HRC | -0.04993 |
| UBASH3B~TOP2A | -0.04993 |
| SPRR2F~PRDM1 | -0.04991 |
| KCNK3~SNHG9 | -0.04991 |
| MUC4~AKR1B15 | -0.04991 |
| RANBP3L~CA3 | -0.04987 |
| AL358334.2~RASL10B | -0.04984 |
| C6~FAT2 | -0.04983 |
| HEY2~PSAT1 | -0.04982 |
| LY6K~HCN2 | -0.04982 |
| RCOR2~AC107976.1 | -0.04982 |
| TRPA1~TMIGD2 | -0.04978 |
| SPATA18~BIRC3 | -0.04976 |
| GABRD~PLA2G7 | -0.04974 |
| P2RY12~AC004233.3 | -0.04973 |
| F13A1~TREML1 | -0.04969 |
| MUC4~AL358334.2 | -0.04967 |
| PTPRO~FOS | -0.04963 |
| SLC46A2~FOS | -0.04962 |

| | |
|---|---|
| C6~CCL22 | -0.04955 |
| CRTAM~ALOX15B | -0.04953 |
| ADAM28~FCN2 | -0.04951 |
| IGHD~TNMD | -0.04948 |
| AC004233.3~OXT | -0.04948 |
| NRIP3~IGSF1 | -0.04946 |
| LINC01703~TMIGD2 | -0.04945 |
| IGFN1~AP000785.2 | -0.04942 |
| TF~NRIP3 | -0.04942 |
| CYP4B1~RCOR2 | -0.04941 |
| AKR1C3~AC004233.3 | -0.04941 |
| HTRA3~ADRB3 | -0.04937 |
| AC131097.2~STRC | -0.04933 |
| OLR1~FOS | -0.0493 |
| CTLA4~PDIA2 | -0.04927 |
| LUCAT1~TRIM55 | -0.04927 |
| GPR171~HS3ST2 | -0.04926 |
| TRIM54~ARL11 | -0.04925 |
| PLA2G7~HMCN2 | -0.04918 |
| RANBP3L~FCN2 | -0.04917 |
| COL9A1~SUSD1 | -0.04917 |
| IGFN1~MAB21L1 | -0.04914 |
| TRIM54~PRDM1 | -0.04912 |
| VNN1~AL049634.2 | -0.04909 |
| RANBP3L~SLC2A4 | -0.04908 |
| ALOX5AP~STRC | -0.04907 |
| TMEM88~GFAP | -0.04902 |
| AC015688.5~HRC | -0.04902 |
| PSAT1~PKP3 | -0.049 |
| AC110285.6~SIRPB1 | -0.04895 |
| UCHL1~SOX10 | -0.04892 |
| AL035406.1~PDIA2 | -0.04891 |
| TFCP2L1~LINC00942 | -0.04886 |
| PTCHD4~CLEC7A | -0.04885 |
| PEBP4~TBX1 | -0.04884 |
| GABRD~HS3ST2 | -0.04878 |
| AC010997.5~BIRC3 | -0.04876 |
| ABCC8~GAS2L2 | -0.04874 |
| PDIA2~GRIN2C | -0.04873 |
| STRC~IGSF1 | -0.04868 |
| GABRD~FCGR2C | -0.04867 |
| PSAT1~CDKN1C | -0.04861 |
| AC010997.5~FOS | -0.04861 |
| AC131097.2~OGDHL | -0.04853 |
| TF~RASSF10 | -0.04851 |
| PDIA2~TLR8 | -0.04851 |
| AL035446.1~ADAM28 | -0.04848 |
| SPATA18~CD180 | -0.04846 |

| | |
|---|---|
| MUC20~SLC6A12 | -0.04844 |
| SPP1~WFDC1 | -0.04843 |
| COL9A1~HOXA9 | -0.04841 |
| GAS2L2~CNTD2 | -0.0484 |
| LINC01770~MAPK8IP2 | -0.04838 |
| BDH1~HS3ST2 | -0.04838 |
| IGHD~FMN1 | -0.04837 |
| CD69~AC110285.6 | -0.04833 |
| IL2RA~CRTAM | -0.04832 |
| FAM78B~THBS1 | -0.04828 |
| ADRB3~MATN4 | -0.04828 |
| RASL11B~IL2RA | -0.04827 |
| MUC4~GCAT | -0.04824 |
| NRTN~CYP2A6 | -0.04822 |
| HSPA6~IGFN1 | -0.0482 |
| MUC4~CCL18 | -0.0482 |
| VNN1~CD300E | -0.04819 |
| FOS~F2RL3 | -0.04817 |
| IL24~MAB21L1 | -0.04815 |
| MIR4458HG~MLIP | -0.04815 |
| TRIM54~POTEF | -0.04814 |
| FHOD3~ANKRD24 | -0.04811 |
| CDKN2B~C20orf204 | -0.04809 |
| SPX~THBS1 | -0.04809 |
| SH2D7~SGCA | -0.04806 |
| NEXN~CLDN9 | -0.04804 |
| AL358334.2~GAS2L2 | -0.04802 |
| AC092720.1~TMIGD2 | -0.04802 |
| PSAT1~TBXA2R | -0.04798 |
| DHRS9~GAS2L2 | -0.04797 |
| TFCP2L1~DMRT2 | -0.04793 |
| GLYAT~MYOCD | -0.04793 |
| MIR4458HG~DCSTAMP | -0.04792 |
| HEY2~CSRP2 | -0.04785 |
| HSPA6~STRC | -0.04783 |
| AKR1B15~PKP3 | -0.04783 |
| PLA2G7~ENHO | -0.0478 |
| VNN1~EXOC3L2 | -0.0478 |
| LINC01703~LINC00942 | -0.04776 |
| PDIA2~AC004233.3 | -0.04776 |
| AKR1C3~SLC27A2 | -0.04774 |
| TRPA1~CLDN9 | -0.04772 |
| OLR1~ALPK3 | -0.04772 |
| LINC01770~PHACTR3 | -0.04771 |
| SPATA18~NRTN | -0.04771 |
| SLC27A2~PDIA2 | -0.04771 |
| KCNK3~OLR1 | -0.04768 |
| WFDC1~GCAT | -0.04768 |

| | |
|---|---|
| AP000785.2~AL358334.2 | -0.04767 |
| AL358334.2~KCNE1 | -0.04765 |
| TRIM54~LINC01230 | -0.04763 |
| IGFN1~PTGDR2 | -0.04756 |
| ADORA3~AL049634.2 | -0.04754 |
| SPX~MAB21L1 | -0.04753 |
| GABRD~EVI2A | -0.0475 |
| ACTG2~HRC | -0.0475 |
| BIRC3~PHACTR3 | -0.0475 |
| NCAPH~AC107976.1 | -0.04749 |
| MUC4~SLC6A12 | -0.04749 |
| IGFN1~Z99774.1 | -0.04747 |
| SLC6A12~SOX10 | -0.04747 |
| ENHO~GDF15 | -0.04745 |
| SDS~AC005696.4 | -0.04744 |
| HPCA~GJC3 | -0.04742 |
| IGFN1~TRIM55 | -0.0474 |
| CA3~PKP3 | -0.04739 |
| SMIM1~POTEF | -0.04738 |
| AC010997.5~PHLDA1 | -0.04737 |
| CYP4B1~GAPT | -0.0473 |
| SPRR2F~HSPA6 | -0.04729 |
| NCAPH~SOX10 | -0.04728 |
| HPCA~LIPA | -0.04721 |
| AC010997.5~ATP8A2 | -0.0472 |
| CYP4B1~AC092118.1 | -0.04714 |
| FCMR~P2RY12 | -0.04706 |
| DPP10~PKP3 | -0.04705 |
| FKBP5~ELFN1 | -0.04704 |
| PEBP4~AC107976.1 | -0.04703 |
| SUSD1~CCL22 | -0.04701 |
| ADAM28~CCL18 | -0.047 |
| CA3~RCOR2 | -0.04699 |
| PTCHD4~GSTZ1 | -0.04698 |
| GABRD~TFCP2L1 | -0.04697 |
| CXCR6~HMCN2 | -0.04697 |
| CA3~TNMD | -0.04697 |
| GFAP~SEMA6B | -0.04694 |
| DPP10~CLDN9 | -0.04691 |
| POTEF~MIR4458HG | -0.04687 |
| TWIST1~Z99774.1 | -0.04684 |
| GABRD~KCNA2 | -0.04679 |
| OGDHL~GAS2L2 | -0.04676 |
| TMIGD3~ABCC8 | -0.04675 |
| MUC20~LINC00968 | -0.04675 |
| RERGL~C5AR2 | -0.04674 |
| FCGR2C~TWIST1 | -0.04672 |
| CA3~TMEM88 | -0.04669 |

| | |
|---|---|
| SLC46A2~CRTAM | -0.04669 |
| ARL4C~KCNE1 | -0.04668 |
| ST14~ATP8A2 | -0.04668 |
| PRDM1~HMCN2 | -0.04667 |
| NCAPH~ALOX5AP | -0.04666 |
| AC131097.2~AKR1B15 | -0.04665 |
| MLIP~IPCEF1 | -0.04665 |
| TREML1~OSBPL3 | -0.04664 |
| IGFN1~LINC01956 | -0.04663 |
| GABRD~MATN4 | -0.04662 |
| AC005696.4~PHACTR3 | -0.04662 |
| RASL11B~RP11-592B15.9 | -0.04658 |
| RASL11B~ITK | -0.04658 |
| HRC~F8A2 | -0.04658 |
| KCNB1~C20orf204 | -0.04657 |
| KCNA2~SLC46A2 | -0.04652 |
| TUBB2A~CD69 | -0.0465 |
| LINC01703~GRIN2C | -0.04648 |
| ELFN1~ATP8A2 | -0.04647 |
| C6~ATP8A2 | -0.04646 |
| AC092720.1~TLR7 | -0.04642 |
| RASL11B~MATN4 | -0.0464 |
| CA3~CLDN9 | -0.0464 |
| FCN2~CCL22 | -0.0464 |
| PRDM1~OLR1 | -0.04638 |
| MYCL~OLR1 | -0.04636 |
| BDH1~PHACTR3 | -0.04636 |
| FOS~SRPX2 | -0.04635 |
| AKR1B15~SLC6A12 | -0.04632 |
| SPRR2F~CDKN2B | -0.0463 |
| RASSF10~CRTAM | -0.0463 |
| SPP1~SOX10 | -0.04629 |
| TFCP2L1~RASL11B | -0.04627 |
| FCMR~AKR1B15 | -0.04626 |
| TRPA1~CD300E | -0.04626 |
| CYP2A6~C20orf204 | -0.04625 |
| TRPA1~C5AR2 | -0.04624 |
| MUC4~GSTZ1 | -0.04619 |
| GLYAT~GRIN2C | -0.04616 |
| TRIM54~GSDMB | -0.04615 |
| SPATA18~SLC27A2 | -0.04611 |
| TGFBR3L~GDF15 | -0.04611 |
| HTRA3~GDF15 | -0.0461 |
| SMIM1~AL035446.1 | -0.04608 |
| FAT2~SLC27A2 | -0.04608 |
| RANBP3L~THBS1 | -0.04607 |
| VNN1~GAS2L2 | -0.04606 |
| ADORA3~SIRPB1 | -0.04604 |

| | |
|---|---|
| SPRR2F~CD69 | -0.04604 |
| GSTZ1~PHACTR3 | -0.04603 |
| SPATA18~CNTD2 | -0.04601 |
| RASSF10~GCAT | -0.046 |
| MIR4458HG~TWIST1 | -0.04595 |
| DACT2~TRPA1 | -0.04593 |
| SOWAHA~WFDC1 | -0.04591 |
| RCOR2~HS3ST2 | -0.04591 |
| VNN1~ELFN1 | -0.04587 |
| ELFN1~LINC00942 | -0.04587 |
| SLC4A8~CNFN | -0.04586 |
| TF~MSR1 | -0.04579 |
| FCGR1A~PHACTR3 | -0.04576 |
| CRTAM~AC092720.1 | -0.04576 |
| PKP3~CLEC7A | -0.04571 |
| IL10~AC004233.3 | -0.04569 |
| NEXN~AC004233.3 | -0.04563 |
| PKP3~MS4A6E | -0.04559 |
| ADRB3~FGF22 | -0.04558 |
| TRIM55~TGFBR3L | -0.04557 |
| IGFN1~TTC36 | -0.04556 |
| AL035446.1~AC092720.1 | -0.04556 |
| AL035446.1~HOXA9 | -0.04556 |
| ADGRL3~AL358334.2 | -0.04555 |
| PHLDA1~KCNB1 | -0.04555 |
| IL1RN~ENHO | -0.04554 |
| FKBP5~ABCC3 | -0.04554 |
| CYP4B1~FCGR1A | -0.0455 |
| CNTD2~GCAT | -0.04549 |
| TMIGD2~CNFN | -0.04548 |
| AC131097.2~PHACTR3 | -0.04547 |
| GABRD~NCEH1 | -0.04545 |
| TRIM54~ZFPM1 | -0.04543 |
| IL1RN~ADGRL3 | -0.04541 |
| NRIP3~AP000785.2 | -0.04541 |
| DACT2~CA3 | -0.0454 |
| DCSTAMP~SGCA | -0.0454 |
| RANBP3L~GALNT6 | -0.04538 |
| AC090181.1~GDF15 | -0.04536 |
| NCAPH~TMIGD2 | -0.04534 |
| POTEF~WDR38 | -0.04534 |
| GFAP~BMP2 | -0.04533 |
| CLDN9~SGCA | -0.04531 |
| FCGR2C~ST14 | -0.04527 |
| AKR1B15~GDF15 | -0.04527 |
| ADAM28~AC110285.6 | -0.04527 |
| POTEF~UCHL1 | -0.04526 |
| LRRC73~ABCC8 | -0.04525 |

| | |
|---|---|
| LINC00968~CA3 | -0.04525 |
| SPATA18~IL2RA | -0.04524 |
| STRC~F8A2 | -0.04523 |
| RRM2~ABCC8 | -0.04521 |
| FAM181B~SLC2A4 | -0.0452 |
| IL24~KCNK3 | -0.04516 |
| AKR1B15~CIB2 | -0.04514 |
| DPP10~AC015688.5 | -0.04511 |
| F13A1~LRRC73 | -0.04511 |
| GAPT~UBASH3B | -0.04509 |
| TWIST1~MAB21L1 | -0.04508 |
| PDIA2~AC134669.2 | -0.04508 |
| PARP15~SGCA | -0.04507 |
| SPX~ADGRE2 | -0.04507 |
| C5AR2~HRC | -0.04504 |
| ATP8A2~SH2D7 | -0.04503 |
| ADAM12~TBX1 | -0.04502 |
| FAM78B~IL24 | -0.04499 |
| C6~SAMD4A | -0.04496 |
| UBASH3B~AC107976.1 | -0.04495 |
| RANBP3L~ATP8A2 | -0.04494 |
| AL035406.1~AL358334.2 | -0.0449 |
| SPATA18~CCL22 | -0.0449 |
| AL035446.1~FASN | -0.0449 |
| DPP10~TREML1 | -0.04484 |
| TREML1~FCN2 | -0.04484 |
| IPCEF1~FOXS1 | -0.04484 |
| PTGDR2~FAM181B | -0.04483 |
| AL035446.1~TOP2A | -0.04482 |
| SOWAHA~C20orf204 | -0.04481 |
| GABRD~TF | -0.04476 |
| IL24~P2RY12 | -0.04476 |
| TRIM54~TUBB2A | -0.04476 |
| CD28~AKR1C3 | -0.04476 |
| LINC00942~CSRP2 | -0.04476 |
| TRIM55~PTGDR2 | -0.04475 |
| MAB21L1~AC004233.3 | -0.04475 |
| IGFN1~GLYCTK | -0.04474 |
| COL9A1~TWIST1 | -0.04472 |
| CA3~PDIA2 | -0.04469 |
| RHOU~PTCHD4 | -0.04467 |
| LINC01956~SLC6A12 | -0.04465 |
| RCOR2~F8A2 | -0.04462 |
| FCGR1A~ENHO | -0.04461 |
| NRIP3~MS4A4A | -0.0446 |
| ERRFI1~EXOC3L2 | -0.04456 |
| DMRT2~HS3ST2 | -0.04455 |
| NUDT8~PHACTR3 | -0.04455 |

| | |
|---|---|
| TF~ANKRD24 | -0.04452 |
| UCHL1~GFAP | -0.04449 |
| MSC~AC004233.3 | -0.04448 |
| SOWAHA~TOP2A | -0.04447 |
| RP11-592B15.9~PHACTR3 | -0.04446 |
| PLEK~SPATA18 | -0.04445 |
| RRM2~ADGRL3 | -0.0444 |
| ATP8A2~TLR7 | -0.0444 |
| TWIST1~CSRP2 | -0.0443 |
| TRPA1~AC004233.3 | -0.0443 |
| AC010997.5~TCF15 | -0.04429 |
| AC005696.4~GFAP | -0.04429 |
| LINC01230~RP11-592B15.9 | -0.04425 |
| CYR61~SPATA18 | -0.04424 |
| CR1~RRM2 | -0.04424 |
| IGFN1~C6 | -0.04423 |
| PTCHD4~CNFN | -0.04422 |
| GJC3~PDIA2 | -0.0442 |
| LINC01956~CDKN1A | -0.04419 |
| MS4A6E~CD69 | -0.04417 |
| HOXA9~GRIN2C | -0.04416 |
| GAPT~CNTD2 | -0.04413 |
| SAMD4A~SGCA | -0.04412 |
| ADAM12~FAM181B | -0.04411 |
| SPX~ALOX15B | -0.04411 |
| CSRNP1~AC010997.5 | -0.0441 |
| FAT2~CA3 | -0.0441 |
| COMTD1~ABCC8 | -0.0441 |
| TRIM54~MTRNR2L12 | -0.04406 |
| ADGRL3~CES1 | -0.04404 |
| CLDN9~KCNB1 | -0.04403 |
| G0S2~AC005696.4 | -0.04401 |
| MATN4~CXorf21 | -0.04401 |
| C6~PTCHD4 | -0.04398 |
| LINC01956~SH2D7 | -0.04397 |
| SLC6A12~OXT | -0.04395 |
| F2RL3~TLR8 | -0.04394 |
| ARL11~CECR2 | -0.04391 |
| CLEC7A~SOX10 | -0.04389 |
| CTLA4~MATN4 | -0.04387 |
| MUC4~SGK1 | -0.04382 |
| PTGDR2~GAS2L2 | -0.04381 |
| PSAT1~CKB | -0.0438 |
| CYP2A6~SOX10 | -0.0438 |
| TFCP2L1~LILRB3 | -0.04379 |
| MS4A6E~CCR10 | -0.04378 |
| TF~CDKN1C | -0.04377 |
| COL9A1~AP000785.2 | -0.04376 |

| | |
|---|---|
| AL035406.1~SIRPB1 | -0.04373 |
| SLC46A2~SLC4A8 | -0.04373 |
| PRR7~DACT2 | -0.04372 |
| RASL11B~CCL22 | -0.04371 |
| NRIP3~ABCC8 | -0.04371 |
| FAT2~AL049634.2 | -0.0437 |
| CLDN9~SIRPB1 | -0.04369 |
| LINC01094~STRC | -0.04367 |
| CSRP2~PHACTR3 | -0.04367 |
| ADAM28~MT-ND3 | -0.04366 |
| TF~SRPX2 | -0.04364 |
| RANBP3L~AL358334.2 | -0.04364 |
| SOWAHA~TIGAR | -0.04364 |
| EVI2A~GRIN2C | -0.04364 |
| CKMT2~SLC27A2 | -0.04361 |
| AP000785.2~LDHD | -0.04359 |
| WDR38~AL049634.2 | -0.04357 |
| RRM2~TRIM55 | -0.04352 |
| CD52~HSPA6 | -0.04351 |
| ADRB3~PKP3 | -0.04351 |
| THBS1~AC110285.6 | -0.0435 |
| FCGR2C~UCHL1 | -0.04349 |
| CYTIP~CERS1 | -0.04349 |
| GAPT~PKP3 | -0.04349 |
| TCF15~TLR8 | -0.04347 |
| AC131097.2~SUSD1 | -0.04345 |
| ALOX5AP~AC004233.3 | -0.04345 |
| TFCP2L1~CXorf21 | -0.04344 |
| NLRC4~PTCHD4 | -0.04342 |
| KCNA2~HOXA9 | -0.04341 |
| KCNA2~MSC | -0.04341 |
| AMPD3~ABCC8 | -0.0434 |
| CNN1~F2RL3 | -0.04339 |
| RORB~RASL10B | -0.04337 |
| MIR4458HG~SLC4A8 | -0.04334 |
| ERRFI1~PHACTR3 | -0.04333 |
| TMIGD3~GDF15 | -0.04333 |
| AC131097.2~PKP3 | -0.04332 |
| CYR61~RANBP3L | -0.04331 |
| G0S2~TLR2 | -0.04331 |
| CA3~CASS4 | -0.04331 |
| CXCR4~PKP3 | -0.04329 |
| TF~LRRC73 | -0.04328 |
| FAT2~FKBP5 | -0.04328 |
| COL9A1~PSAT1 | -0.04328 |
| IL10~SPATA18 | -0.04326 |
| SPATA18~MMP9 | -0.04324 |
| PTCHD4~TIGAR | -0.04323 |

| | |
|---|---|
| MYCL~CSRP2 | -0.04321 |
| AC010997.5~PLIN5 | -0.0432 |
| F13A1~CRTAM | -0.04319 |
| OLR1~SMOC1 | -0.04319 |
| FCMR~TF | -0.04317 |
| SPATA18~MLIP | -0.04317 |
| GDF15~EXOC3L2 | -0.04317 |
| GABRD~C9 | -0.04315 |
| HSPA6~ID4 | -0.04315 |
| FKBP5~APLNR | -0.04314 |
| LINC01956~CNN1 | -0.04312 |
| SLC27A2~ABCC3 | -0.04312 |
| GABRD~TM4SF19 | -0.04311 |
| IGHD~SRPX2 | -0.04308 |
| RRM2~CYP2A6 | -0.04307 |
| FAM78B~CSRP2 | -0.04306 |
| IGFN1~TM7SF2 | -0.04306 |
| PTCHD4~ALPK3 | -0.04306 |
| FCGR2B~LRRC73 | -0.04302 |
| ADAM28~STRC | -0.04301 |
| RASL10B~CD300E | -0.04301 |
| G0S2~TFCP2L1 | -0.043 |
| RCOR2~GDF15 | -0.04297 |
| MS4A6E~HCN2 | -0.04296 |
| AZGP1~HCN2 | -0.04295 |
| HSPA6~SGCA | -0.04293 |
| CNTD2~IGSF1 | -0.04291 |
| LINC01230~IGHD | -0.0429 |
| PHLDA1~MAB21L1 | -0.04289 |
| ADORA3~HOXA9 | -0.04281 |
| HSPA6~COL9A1 | -0.04281 |
| CYTIP~GDF1 | -0.04277 |
| SGCA~NRTN | -0.04277 |
| NCAPH~PKP3 | -0.04276 |
| FGF22~NRTN | -0.04276 |
| IL7R~HS3ST2 | -0.04271 |
| AC010997.5~AC011498.4 | -0.04271 |
| MSC~ENHO | -0.04269 |
| SDS~F8A2 | -0.04264 |
| CHIT1~ID4 | -0.04263 |
| HPCA~LINC01230 | -0.0426 |
| ARL4C~GFAP | -0.0426 |
| AC131097.2~MUC20 | -0.0426 |
| TM4SF19~SMOC1 | -0.0426 |
| LY6K~AC025262.1 | -0.0426 |
| SPATA18~GDF1 | -0.04259 |
| PRDM1~AC010997.5 | -0.04259 |
| PLA2G7~TMIGD2 | -0.04258 |

| | |
|---|---|
| AL035406.1~IGSF6 | -0.04257 |
| KCNA2~BIRC3 | -0.04257 |
| FHOD3~HRC | -0.04257 |
| LINC01703~AKR1B15 | -0.04256 |
| DMRT2~IGHD | -0.04256 |
| AL358334.2~CCL22 | -0.04255 |
| GABRD~GCNT1 | -0.04254 |
| KANK4~SNHG9 | -0.04254 |
| TREML1~CDKN1C | -0.04253 |
| PHACTR3~C20orf204 | -0.04253 |
| POTEF~OXT | -0.04252 |
| MUC20~HOXA9 | -0.04252 |
| AP000785.2~SOX10 | -0.04252 |
| PTCHD4~LINC01230 | -0.04251 |
| RERGL~AC092720.1 | -0.04251 |
| HOXA9~CERS1 | -0.04248 |
| LRRC73~HS3ST2 | -0.04247 |
| SPRR2F~ART4 | 0.04247 |
| TF~OXT | 0.04247 |
| STRC~TMEM88 | 0.04247 |
| CYR61~SMOC1 | 0.04248 |
| DCSTAMP~CCL22 | 0.04248 |
| SPX~HS3ST2 | 0.04248 |
| NCAPH~TREM2 | 0.0425 |
| SOWAHA~ADGRE2 | 0.0425 |
| FCGR2B~ENHO | 0.04251 |
| RGS1~SNHG9 | 0.04252 |
| TLR2~CLEC4E | 0.04252 |
| GAPT~SGCA | 0.04253 |
| DUSP1~GPR183 | 0.04253 |
| SMAP2~CLEC4E | 0.04254 |
| ELFN1~HCN2 | 0.04254 |
| OGDHL~NRIP3 | 0.04256 |
| ADAM28~CRTAM | 0.04257 |
| RASSF10~SGCA | 0.04257 |
| ADRB3~FOS | 0.04259 |
| LDHD~REEP6 | 0.0426 |
| MSC~SIRPB1 | 0.04261 |
| AC092720.1~TGFBR3L | 0.04261 |
| IQGAP2~F13A1 | 0.04263 |
| AKR1B15~IGSF6 | 0.04266 |
| AKR1B15~ALOX15B | 0.04269 |
| CXCR4~IL7R | 0.0427 |
| LILRB3~OXT | 0.0427 |
| MYCL~BDH1 | 0.04271 |
| RORB~ALPK3 | 0.04271 |
| CLDN9~AC005696.4 | 0.04272 |
| MUC4~TGFBR3L | 0.04273 |

| | |
|---|---|
| RANBP3L~SRPX2 | 0.04273 |
| CKMT2~NUDT8 | 0.04273 |
| DACT2~OXT | 0.04273 |
| GLIS1~LINC01703 | 0.04274 |
| DACT2~ELFN1 | 0.04274 |
| IPCEF1~AC092720.1 | 0.04275 |
| SLC46A2~SLC6A12 | 0.04275 |
| IL24~OLR1 | 0.04276 |
| IL7R~PRDM1 | 0.04276 |
| PRR7~TMEM88 | 0.04276 |
| TRPA1~SLC6A12 | 0.04276 |
| CA3~WFDC1 | 0.04276 |
| CXCR4~CASS4 | 0.04278 |
| PTGDR2~HS3ST2 | 0.04278 |
| CHIT1~MMP9 | 0.04279 |
| THBS1~SAMSN1 | 0.04279 |
| SPATA18~AC010997.5 | 0.0428 |
| FAM181B~EXOC3L2 | 0.0428 |
| AC131097.2~CSRP2 | 0.04281 |
| AKR1B15~AKR1C3 | 0.04281 |
| OGDHL~TGFBR3L | 0.04282 |
| AP000785.2~CRTAM | 0.04283 |
| CD52~SNHG9 | 0.04284 |
| HOXA9~SIRPB1 | 0.04284 |
| RP11-592B15.9~SLC27A2 | 0.04284 |
| TRPA1~RERGL | 0.04285 |
| THEMIS2~C5AR1 | 0.04286 |
| HMCN2~MATN4 | 0.04286 |
| HS3ST2~MMP9 | 0.04286 |
| MYCL~NRIP3 | 0.04287 |
| ST14~AC005696.4 | 0.04287 |
| ATP8A2~AL358334.2 | 0.04288 |
| AC110285.6~HCN2 | 0.04289 |
| CTLA4~ADAM28 | 0.0429 |
| MIR4458HG~TIGAR | 0.0429 |
| NRTN~CERS1 | 0.0429 |
| THEMIS2~SDS | 0.04291 |
| HTRA3~GDF1 | 0.04291 |
| GFAP~REEP6 | 0.04292 |
| CHIT1~AC005696.4 | 0.04293 |
| GJC3~SNHG9 | 0.04293 |
| GABRD~NLRC4 | 0.04294 |
| IGFN1~PARP15 | 0.04294 |
| CHIT1~TMIGD2 | 0.04294 |
| POTEF~IGSF1 | 0.04295 |
| TRIM55~RP11-592B15.9 | 0.04295 |
| MYCL~GFI1 | 0.04299 |
| PLEK~CRTAM | 0.04299 |

| | |
|---|---|
| IL1RN~DHRS9 | 0.043 |
| LRRC46~CNFN | 0.043 |
| MTRNR2L12~SUSD1 | 0.04302 |
| TF~CLEC7A | 0.04305 |
| CD52~FAM181B | 0.04307 |
| CYP4B1~SLAMF6 | 0.04307 |
| MLIP~ALPK3 | 0.04309 |
| LINC01703~CD300E | 0.0431 |
| UBASH3B~LILRB3 | 0.0431 |
| BCAT1~IGHD | 0.0431 |
| SNHG9~LRRC46 | 0.0431 |
| SLC2A4~TMIGD2 | 0.0431 |
| MIR4458HG~CCL22 | 0.04311 |
| PTCHD4~KCNB1 | 0.04311 |
| SPP1~AC015688.5 | 0.04312 |
| PRDM1~SIRPB1 | 0.04312 |
| CYP4B1~MUC4 | 0.04313 |
| G0S2~MLIP | 0.04313 |
| HOXA9~RERGL | 0.04313 |
| SLAMF6~IL7R | 0.04314 |
| AC131097.2~SNHG9 | 0.04315 |
| IGHD~Z99774.1 | 0.04315 |
| LINC01770~TFCP2L1 | 0.04316 |
| UBASH3B~AC005696.4 | 0.04316 |
| SMIM1~DACT2 | 0.04317 |
| HSPA6~SPATA18 | 0.04317 |
| LINC01703~ENHO | 0.04317 |
| F13A1~MS4A7 | 0.04318 |
| CA3~SOX10 | 0.0432 |
| LRRC73~CRTAM | 0.04321 |
| SMOC1~NDRG4 | 0.04321 |
| PLA2G7~DCSTAMP | 0.04322 |
| CXCR4~MYOCD | 0.04323 |
| LRRC73~ELFN1 | 0.04324 |
| UBASH3B~SIRPB1 | 0.04324 |
| LINC01230~APLNR | 0.04328 |
| SLC6A12~GDF15 | 0.04329 |
| RGS2~FOSL2 | 0.0433 |
| DMRT2~AC025262.1 | 0.0433 |
| HCN2~NRTN | 0.04331 |
| GLYCTK~SLC6A12 | 0.04335 |
| CARMN~LINC01230 | 0.04335 |
| IL1RN~DCSTAMP | 0.04336 |
| BDH1~IGHD | 0.04336 |
| HS3ST2~F2RL3 | 0.04336 |
| HTRA3~GAPT | 0.0434 |
| PLEK~LINC00942 | 0.04342 |
| CD28~PTCHD4 | 0.04342 |

| | |
|---|---|
| ADGRL3~RASSF10 | 0.04342 |
| KCNK3~ALPK3 | 0.04344 |
| UCHL1~LINC00942 | 0.04344 |
| CYP4B1~PHACTR3 | 0.04345 |
| ADGRL3~RORB | 0.04347 |
| TF~HTRA3 | 0.04349 |
| RASL11B~ART4 | 0.04349 |
| PTGDR2~SNHG9 | 0.04349 |
| CXCR6~Z99774.1 | 0.04352 |
| CYP2A6~SMIM25 | 0.04352 |
| ARL4C~GPR183 | 0.04353 |
| FCN2~SLC2A4 | 0.04354 |
| PTGDR2~WFDC1 | 0.04354 |
| AC011498.4~SOX10 | 0.04355 |
| GAPT~F2RL3 | 0.04356 |
| CSRP2~SLC2A4 | 0.04356 |
| PHLDA1~KCNE1 | 0.04357 |
| RASSF10~GFAP | 0.04358 |
| DUSP1~C5AR1 | 0.04361 |
| PTCHD4~AKR1B15 | 0.04361 |
| NCAPH~ENHO | 0.04362 |
| SPRR2F~IQGAP2 | 0.04363 |
| LIPA~ADGRE2 | 0.04363 |
| CHIT1~CNTD2 | 0.04364 |
| POTEF~RP11-592B15.9 | 0.04364 |
| KLHL6~ALPK3 | 0.04365 |
| PTCHD4~FMN1 | 0.04365 |
| SMOC1~ALPK3 | 0.04365 |
| SGCA~ACP5 | 0.04365 |
| ERRFI1~BIRC3 | 0.04368 |
| FAM181B~FOXS1 | 0.04368 |
| COL9A1~Z99774.1 | 0.04369 |
| RASSF10~HS3ST2 | 0.04369 |
| RCOR2~NQO1 | 0.04369 |
| P2RY12~SOX10 | 0.0437 |
| SOWAHA~NXNL1 | 0.04371 |
| CARMN~WFDC1 | 0.04371 |
| TREML1~AC005696.4 | 0.04371 |
| FCGR1A~ATP8A2 | 0.04372 |
| HSPA6~ARL4C | 0.04372 |
| ADGRL3~LY6K | 0.04373 |
| TIGAR~AC004233.3 | 0.04373 |
| SAMD4A~CCL22 | 0.04373 |
| LINC01770~ID4 | 0.04374 |
| UCHL1~CNFN | 0.04374 |
| SGK1~PHLDA1 | 0.04375 |
| KCNA2~ACP5 | 0.04377 |
| ADAM28~MAPK8IP2 | 0.04377 |

| | |
|---|---|
| ADAM12~TENM4 | 0.04377 |
| ADORA3~EXOC3L2 | 0.0438 |
| SPRR2F~CARMN | 0.04381 |
| DPP10~MT-ATP6 | 0.04382 |
| KCNA2~VNN1 | 0.04383 |
| LINC01230~AC025262.1 | 0.04383 |
| HSPA6~CA3 | 0.04384 |
| NLRC4~AC005696.4 | 0.04384 |
| DPP10~Z99774.1 | 0.04387 |
| SDS~GPR183 | 0.04387 |
| RGS2~CASS4 | 0.04391 |
| GALNT6~MAB21L1 | 0.04392 |
| SPRR2F~CYP2A6 | 0.04393 |
| ABCC8~SLC6A12 | 0.04393 |
| CTLA4~PARP15 | 0.04394 |
| TIGAR~AC069368.1 | 0.04394 |
| PLEK~CCR1 | 0.04395 |
| MLIP~SIK1 | 0.04396 |
| MLIP~CU639417.2 | 0.04396 |
| ARL4C~LILRB3 | 0.04397 |
| HTRA3~THBS1 | 0.04401 |
| LUCAT1~HMCN2 | 0.04401 |
| GLYAT~CES1 | 0.04403 |
| FCGR3A~KYNU | 0.04404 |
| GPT~SLC2A4 | 0.04405 |
| CDKN2B~PHACTR3 | 0.04406 |
| SUSD1~AL049634.2 | 0.04406 |
| TRIM55~SMOC1 | 0.04408 |
| FCGR2C~LYZ | 0.04411 |
| MIR4458HG~ANKRD24 | 0.04411 |
| ALDH4A1~BMP2 | 0.04412 |
| AL035406.1~AC134669.2 | 0.04413 |
| ITK~C5orf58 | 0.04413 |
| RORB~AP000785.2 | 0.04413 |
| PKP3~UBASH3B | 0.04413 |
| MTRNR2L12~MT-CO2 | 0.04414 |
| IGFN1~FAT2 | 0.04415 |
| CYTIP~CSRNP1 | 0.04415 |
| AZGP1~RASL10B | 0.04415 |
| PTGDR2~CKB | 0.04415 |
| G0S2~SPX | 0.04416 |
| OSBPL3~LILRB3 | 0.04416 |
| ITGAX~CD300LB | 0.04416 |
| ZC3H12D~PHLDA1 | 0.04419 |
| PEBP4~REEP6 | 0.04421 |
| GALNT6~IGSF6 | 0.04423 |
| SNHG9~CNFN | 0.04423 |
| AL035406.1~HS3ST2 | 0.04424 |

| | |
|---|---|
| TMEM88~CYP2A6 | 0.04424 |
| GCNT1~NRIP3 | 0.04425 |
| CTLA4~PLA2G7 | 0.04426 |
| IGSF1~MT-CYB | 0.04426 |
| UCHL1~GDF1 | 0.04428 |
| PKP3~SMOC1 | 0.04428 |
| MYCL~CXCR4 | 0.04429 |
| FAM78B~GDF1 | 0.04429 |
| LINC00968~PSAT1 | 0.0443 |
| GFI1~DPP10 | 0.04431 |
| AC004233.3~IGSF1 | 0.04433 |
| HSPA6~MSC | 0.04434 |
| LY6K~TIGAR | 0.04434 |
| CTLA4~CD69 | 0.04435 |
| F13A1~ALOX5AP | 0.04435 |
| HMCN2~CNN1 | 0.04435 |
| FCGR3A~ENHO | 0.04436 |
| FCGR1A~ADAM28 | 0.04438 |
| FAM78B~HTRA3 | 0.04438 |
| AKR1B15~THBS1 | 0.04438 |
| CTSB~HS3ST2 | 0.04438 |
| FCGR2B~ATP8A2 | 0.04439 |
| MLIP~CKB | 0.04439 |
| PTGDR2~STRC | 0.04439 |
| IGHD~BMP2 | 0.04439 |
| DPP10~CD300E | 0.0444 |
| WDR38~CNFN | 0.0444 |
| DRD4~CLDN9 | 0.0444 |
| MLIP~TM7SF2 | 0.04441 |
| DUSP1~CDKN1A | 0.04444 |
| ABCC8~HS3ST2 | 0.04445 |
| AC131097.2~TUBB2A | 0.04446 |
| UBASH3B~ALOX15B | 0.04446 |
| FCN2~LILRB3 | 0.04448 |
| RERGL~ABCC3 | 0.04448 |
| NCAPH~SLC6A12 | 0.04449 |
| CMTM8~WFDC1 | 0.04449 |
| APLNR~TMEM88 | 0.04452 |
| HSPA6~IGHD | 0.04453 |
| ID4~FCN2 | 0.04454 |
| CLDN9~TLR7 | 0.04455 |
| SUSD1~AL358334.2 | 0.04456 |
| MIR4458HG~TBXA2R | 0.04457 |
| CXCR4~GPR171 | 0.04458 |
| NCEH1~NRIP3 | 0.04458 |
| VNN1~CLDN9 | 0.04458 |
| DHRS9~AL358334.2 | 0.04459 |
| FAT2~CD300E | 0.0446 |

| | |
|---|---|
| GLYCTK~AL035446.1 | 0.04461 |
| DMRT2~CRTAM | 0.04463 |
| COL9A1~SOX10 | 0.04464 |
| TFCP2L1~MATN4 | 0.04465 |
| GLYCTK~SGCA | 0.04465 |
| F13A1~CD163L1 | 0.04465 |
| PSAT1~AC004233.3 | 0.04465 |
| HSPA6~SOWAHA | 0.04466 |
| MTRNR2L12~MT-ND3 | 0.04466 |
| SUSD1~AC015688.5 | 0.04467 |
| CYR61~PDIA2 | 0.04469 |
| FCGR2A~LINC01956 | 0.04469 |
| TBX1~IGSF1 | 0.04469 |
| LUCAT1~SLC46A2 | 0.04473 |
| TRPA1~BIRC3 | 0.04473 |
| GSTZ1~TGFBR3L | 0.04474 |
| ERRFI1~CSRNP1 | 0.04475 |
| LY6K~AC005696.4 | 0.04475 |
| AP000785.2~HS3ST2 | 0.04475 |
| SH2D7~CD300E | 0.04476 |
| RGS2~CNN1 | 0.04477 |
| TRIM54~TGFBR3L | 0.04477 |
| SUSD1~AC107976.1 | 0.04477 |
| HTRA3~CCL18 | 0.04478 |
| LY6K~PHACTR3 | 0.04478 |
| P2RY12~MUC4 | 0.04479 |
| SOWAHA~SOX10 | 0.04479 |
| SPX~CCL22 | 0.0448 |
| AP000785.2~GSTZ1 | 0.04481 |
| STRC~AC004233.3 | 0.04481 |
| TRIM54~PDIA2 | 0.04482 |
| SLC6A12~HS3ST2 | 0.04484 |
| TRIM54~CRTAM | 0.04485 |
| DPP10~CCL18 | 0.04485 |
| FCGR2C~OSBPL3 | 0.04486 |
| RASSF10~NDRG4 | 0.04486 |
| LINC01770~C5orf58 | 0.04488 |
| RASL10B~OXT | 0.04488 |
| GLIS1~FHOD3 | 0.04489 |
| UCHL1~CERS1 | 0.0449 |
| AL358334.2~CLDN9 | 0.0449 |
| TF~RASL11B | 0.04492 |
| ADRB3~CLEC7A | 0.04492 |
| STRC~MAPK8IP2 | 0.04493 |
| FASN~HCN2 | 0.04493 |
| WDR38~LRRC46 | 0.04496 |
| AC107976.1~PDIA2 | 0.04496 |
| LYZ~CD300E | 0.04498 |

| | |
|---|---|
| IGSF6~MATN4 | 0.04498 |
| CYP2A6~MATN4 | 0.04498 |
| ATP8A2~TBXA2R | 0.04499 |
| SUSD1~ABCC8 | 0.045 |
| RGS1~CXCR4 | 0.04502 |
| KLHL6~UBASH3B | 0.04502 |
| CTLA4~MSC | 0.04503 |
| CD180~CRTAM | 0.04503 |
| TRIM54~THBS1 | 0.04504 |
| SDS~CD300E | 0.04504 |
| AC131097.2~TRIM55 | 0.04506 |
| NCAPH~RASL11B | 0.04507 |
| IPCEF1~CECR2 | 0.04508 |
| ADGRL3~CASS4 | 0.04509 |
| LINC01230~CES1 | 0.04509 |
| LEP~GLYAT | 0.0451 |
| AC131097.2~CCR5 | 0.04515 |
| CCL22~ADGRE2 | 0.04515 |
| TRIM54~ATP8A2 | 0.04517 |
| SDS~PHACTR3 | 0.04519 |
| CHIT1~NCAPH | 0.04523 |
| FCN2~PHLDA1 | 0.04527 |
| ART4~TBX1 | 0.04528 |
| AC004233.3~MT-CO2 | 0.04529 |
| NRIP3~WFDC1 | 0.04531 |
| CARMN~HRC | 0.04532 |
| HMCN2~RERGL | 0.04536 |
| ANKRD24~F2RL3 | 0.04536 |
| HMCN2~MAB21L1 | 0.04538 |
| KCNK3~TIGAR | 0.04539 |
| LY6K~FHOD3 | 0.04539 |
| LINC01703~CLDN9 | 0.04542 |
| IGHD~TGFBR3L | 0.04542 |
| DHRS9~HS3ST2 | 0.04543 |
| PRDM1~ADAM28 | 0.04543 |
| LINC00942~F2RL3 | 0.04545 |
| IFI30~SMIM25 | 0.04546 |
| GAPT~SIRPB1 | 0.04548 |
| DACT2~GRIN2C | 0.04549 |
| NDRG4~AC015688.5 | 0.04549 |
| SPATA18~AL035446.1 | 0.0455 |
| CYR61~TREML1 | 0.04552 |
| IGFN1~MMP9 | 0.04554 |
| DACT2~AL358334.2 | 0.04554 |
| HMCN2~GRIN2C | 0.04554 |
| FHOD3~PHACTR3 | 0.04554 |
| ST14~AL049634.2 | 0.04556 |
| FKBP5~IL2RA | 0.04558 |

| | |
|---|---|
| TENM4~GAS2L2 | 0.04558 |
| TREM2~ALOX15B | 0.04559 |
| GABRD~CIB2 | 0.0456 |
| FCGR2B~CD69 | 0.04561 |
| SPRR2F~STRC | 0.04562 |
| FCMR~CERS1 | 0.04562 |
| ABCC8~CECR2 | 0.04562 |
| LINC00942~PDIA2 | 0.04562 |
| CD28~HEY2 | 0.04563 |
| ADGRL3~CNN1 | 0.04563 |
| ADGRL3~CKMT2 | 0.04563 |
| PKP3~CIB2 | 0.04563 |
| LINC01956~PTGDR2 | 0.04564 |
| SLC4A8~AC110285.6 | 0.04564 |
| PROB1~TMEM88 | 0.04566 |
| ABCC8~ANKRD24 | 0.04567 |
| KCNB1~PHACTR3 | 0.04567 |
| CD52~NCAPH | 0.04569 |
| LDHD~CHCHD10 | 0.04569 |
| TRIM54~ADRB3 | 0.0457 |
| LINC01094~CRTAM | 0.0457 |
| PRR7~TUBB2A | 0.0457 |
| CTLA4~AP000785.2 | 0.04572 |
| CD69~AC005696.4 | 0.04572 |
| CYP4B1~P2RY12 | 0.04573 |
| ZC3H12D~CLDN9 | 0.04573 |
| WFDC1~PHACTR3 | 0.04573 |
| DUSP1~SGK1 | 0.04575 |
| MYCL~TMIGD3 | 0.04576 |
| HOXA9~AL049634.2 | 0.04576 |
| CSRP2~GCAT | 0.04578 |
| NQO1~SRPX2 | 0.04578 |
| RRM2~MTRNR2L12 | 0.0458 |
| RASL11B~HOXA9 | 0.04582 |
| CCL22~GAS2L2 | 0.04582 |
| DPP10~TOP2A | 0.04583 |
| HEY2~RERGL | 0.04584 |
| CYP4B1~AL035446.1 | 0.04588 |
| CCL22~MMP9 | 0.04588 |
| ID4~PHLDA1 | 0.0459 |
| GAPT~ID4 | 0.04592 |
| AC131097.2~PSAT1 | 0.04594 |
| HEY2~TWIST1 | 0.04596 |
| SLAMF8~IGFN1 | 0.04599 |
| LY6K~AKR1C3 | 0.04599 |
| FAT2~STRC | 0.04601 |
| HPCA~MAB21L1 | 0.04602 |
| PTGDR2~PHACTR3 | 0.04603 |

| | |
|---|---|
| LINC01956~UBASH3B | 0.04604 |
| TUBB2A~TBXA2R | 0.04606 |
| RGS2~GAPT | 0.04607 |
| CCR5~GPR171 | 0.04607 |
| MUSTN1~TBX1 | 0.04608 |
| C6~FCN2 | 0.04613 |
| AKR1C3~CES1 | 0.04613 |
| LINC00942~CES1 | 0.04613 |
| MUC4~PTGDR2 | 0.04614 |
| TREML1~CRTAM | 0.04614 |
| MSC~MAB21L1 | 0.04614 |
| HEY2~MYOCD | 0.04615 |
| AKR1B15~MSC | 0.04615 |
| TGFBR3L~TBX1 | 0.04615 |
| HTRA3~AP000785.2 | 0.04616 |
| TNFAIP3~CD69 | 0.04618 |
| LINC00942~CXorf21 | 0.04618 |
| GDF15~SIK1 | 0.04619 |
| DPP10~AL049634.2 | 0.0462 |
| GDF15~CU639417.2 | 0.0462 |
| VNN1~GLYAT | 0.04621 |
| SPRR2F~TMEM88 | 0.04622 |
| TENM4~REEP6 | 0.04623 |
| IL10~SOX10 | 0.04624 |
| PTGDR2~SLC2A4 | 0.04624 |
| FCGR1A~NRTN | 0.04626 |
| CHIT1~ST14 | 0.04626 |
| CD52~SPP1 | 0.04628 |
| GPR171~IL7R | 0.04628 |
| FAM78B~DACT2 | 0.0463 |
| RASL10B~AC011498.4 | 0.0463 |
| SOWAHA~AC011498.4 | 0.04634 |
| ADRB3~ENHO | 0.04635 |
| SLC4A8~CYP2A6 | 0.04635 |
| SPATA18~SIRPB1 | 0.04639 |
| PXMP2~AC135586.2 | 0.04639 |
| LINC01956~RORB | 0.0464 |
| FCGR1A~CCR1 | 0.04642 |
| DHRS9~GDF15 | 0.04642 |
| KCNK3~TBXA2R | 0.04644 |
| CTLA4~TRIM55 | 0.04644 |
| ADRB3~LIPA | 0.04645 |
| HSPA6~BCAT1 | 0.04646 |
| TUBB2A~CNN1 | 0.04647 |
| IGHD~OXT | 0.04647 |
| VNN1~ANKRD24 | 0.04649 |
| AC090181.1~GAS2L2 | 0.04649 |
| FCGR2C~CCR10 | 0.04652 |

| | |
|---|---|
| LINC01703~SOWAHA | 0.04655 |
| CD69~AL049634.2 | 0.04655 |
| KANK4~FAT2 | 0.04656 |
| MT-ND3~MT-CYB | 0.04656 |
| NEXN~AC131097.2 | 0.04657 |
| ENHO~TBX1 | 0.04657 |
| LINC01770~FCGR1A | 0.0466 |
| HSPA6~GPR65 | 0.04661 |
| CD28~RCOR2 | 0.04662 |
| LINC01956~SLC27A2 | 0.04663 |
| AC131097.2~ST14 | 0.04664 |
| CD28~FCN2 | 0.04666 |
| CTLA4~TF | 0.04667 |
| CTLA4~DUSP1 | 0.0467 |
| NPL~VNN1 | 0.04672 |
| RASL10B~CCL18 | 0.04673 |
| AC131097.2~MAB21L1 | 0.04674 |
| NCAPH~SNHG9 | 0.04675 |
| AC092118.1~EXOC3L2 | 0.04675 |
| LINC00942~BMP2 | 0.04677 |
| KCNK3~TNFSF8 | 0.04678 |
| ADGRL3~SMOC1 | 0.04679 |
| CYP4B1~KCNE1 | 0.0468 |
| TUBB2A~CRTAM | 0.04681 |
| LIPA~UBASH3B | 0.04681 |
| TMIGD3~GAPT | 0.04682 |
| ALDH4A1~CYP4B1 | 0.04683 |
| TMIGD3~CHIT1 | 0.04683 |
| ADRB3~ALPK3 | 0.04683 |
| RCOR2~STRC | 0.04685 |
| CD28~AMPD3 | 0.04686 |
| CYP4B1~FCGR2B | 0.04687 |
| HOXA9~SUSD1 | 0.04687 |
| NEXN~TUBB2A | 0.04689 |
| NCAPH~AC004233.3 | 0.0469 |
| DACT2~CLDN9 | 0.0469 |
| TRIM54~GFAP | 0.04693 |
| DACT2~HCN2 | 0.04696 |
| IGHD~MYOCD | 0.04697 |
| TRIM54~TMIGD2 | 0.04699 |
| HMCN2~GCAT | 0.04699 |
| FCGR1A~SPP1 | 0.047 |
| MT-CO3~MT-ND3 | 0.047 |
| RGS1~KCNE1 | 0.04701 |
| ACTG2~HOXA9 | 0.04703 |
| P2RY12~ATP8A2 | 0.04703 |
| CYP4B1~SESN1 | 0.04704 |
| PTCHD4~GFAP | 0.04704 |

| | |
|---|---|
| PRDM1~CLEC4E | 0.04705 |
| SLC46A2~ATP8A2 | 0.04706 |
| SLC6A12~ABCC3 | 0.04707 |
| DACT2~AC010997.5 | 0.04709 |
| TRIM54~CECR2 | 0.0471 |
| SMOC1~TBX1 | 0.0471 |
| EVI2A~TLR8 | 0.0471 |
| GABRD~ST14 | 0.04712 |
| WFDC1~AC110285.6 | 0.04712 |
| SPRR2F~PTPRO | 0.04714 |
| MSC~SAMD4A | 0.04714 |
| MUC4~SNHG9 | 0.04717 |
| LUCAT1~TIGAR | 0.04717 |
| LINC01703~CIB2 | 0.04719 |
| LRRC73~GCAT | 0.04721 |
| ST14~CCL18 | 0.04721 |
| IGHD~RASL10B | 0.04723 |
| P2RY12~RANBP3L | 0.04724 |
| CD52~ATP8A2 | 0.04725 |
| PRDM1~SUSD1 | 0.04726 |
| TRIM54~AL358334.2 | 0.04727 |
| ADGRL3~CECR2 | 0.04727 |
| FCN2~GFAP | 0.04728 |
| ADAM28~FGF22 | 0.04729 |
| POTEF~HS3ST2 | 0.0473 |
| TM4SF19~DCSTAMP | 0.04731 |
| LINC01703~CYP2A6 | 0.04732 |
| C6~EXOC3L2 | 0.04732 |
| NRIP3~GSTZ1 | 0.04733 |
| ACTG2~PKP3 | 0.04734 |
| APLNR~ANKRD24 | 0.04735 |
| AP000785.2~OXT | 0.04735 |
| TFCP2L1~HCN2 | 0.04737 |
| DRD4~AC092720.1 | 0.04737 |
| LINC01956~CLEC4E | 0.0474 |
| MTRNR2L12~MT-CYB | 0.04744 |
| CHIT1~MS4A6E | 0.04745 |
| HPCA~GCAT | 0.04746 |
| SPRR2F~MATN4 | 0.04747 |
| IL10~DUSP1 | 0.04747 |
| ITGAX~CD300E | 0.04747 |
| CCL22~CCR10 | 0.04748 |
| C5AR1~PHACTR3 | 0.04749 |
| NCAPH~AMPD3 | 0.04751 |
| AL035446.1~CCL18 | 0.04753 |
| AL358334.2~PDIA2 | 0.04754 |
| MT-ND3~MT-ND4 | 0.04754 |
| HTRA3~FHOD3 | 0.04755 |

| | |
|---|---|
| AC090181.1~CES1 | 0.04755 |
| POTEF~Z99774.1 | 0.04756 |
| CD28~ABCC8 | 0.04758 |
| TRIM54~LRRC73 | 0.04759 |
| HSPA6~POTEF | 0.0476 |
| ACTG2~TBX1 | 0.0476 |
| MT-ATP6~MT-ND3 | 0.04761 |
| MLIP~ATP8A2 | 0.04762 |
| SOWAHA~PLIN5 | 0.04764 |
| FCGR2B~CCL22 | 0.04767 |
| SOWAHA~FCN2 | 0.04767 |
| MTRNR2L12~MT-ND1 | 0.04768 |
| TLR2~MIR4458HG | 0.04768 |
| NRIP3~HS3ST2 | 0.04768 |
| TMEM52~LINC01703 | 0.04769 |
| POTEF~TIGAR | 0.04771 |
| AMPD3~SIRPB1 | 0.04771 |
| FAM78B~FAM181B | 0.04772 |
| SLC46A2~CLDN9 | 0.04776 |
| GFI1~ART4 | 0.04777 |
| HPCA~PDIA2 | 0.04778 |
| CYP4B1~AC131097.2 | 0.04778 |
| KCNA2~LIPA | 0.04778 |
| FAM78B~DPP10 | 0.04779 |
| SPATA18~GSTZ1 | 0.04779 |
| GLIS1~GAPT | 0.0478 |
| PRR7~ANKRD24 | 0.0478 |
| CDKN2B~TENM4 | 0.0478 |
| CA3~GFAP | 0.04781 |
| UBASH3B~LINC00942 | 0.04782 |
| FCMR~AL035446.1 | 0.04783 |
| ACTG2~HTRA3 | 0.04784 |
| NCAPH~UBASH3B | 0.04785 |
| GLIS1~SOWAHA | 0.04788 |
| GPR171~TMIGD2 | 0.04788 |
| PRDM1~AL049634.2 | 0.04792 |
| CMTM8~AC092720.1 | 0.04794 |
| MUC4~AC107976.1 | 0.04795 |
| NCAPH~ACP5 | 0.04796 |
| GFI1~ITK | 0.04799 |
| NCAPH~ITGAX | 0.04802 |
| TNFAIP3~PHLDA1 | 0.04802 |
| RCOR2~SOX10 | 0.04808 |
| FAT2~RCOR2 | 0.04809 |
| GCNT1~OGDHL | 0.04811 |
| FAM181B~IGHD | 0.04815 |
| MUC20~CRTAM | 0.04817 |
| FCGR2B~SPATA18 | 0.0482 |

| | |
|---|---|
| TM4SF19~TRPA1 | 0.04821 |
| ADGRL3~MMP9 | 0.04821 |
| CHIT1~SOWAHA | 0.04822 |
| BDH1~MSC | 0.04822 |
| IGHD~ALPK3 | 0.04822 |
| GABRD~FCGR1A | 0.04823 |
| CSRP2~GSDMB | 0.04823 |
| VNN1~FGF22 | 0.04826 |
| CA3~CNN1 | 0.04827 |
| LY6K~MMP9 | 0.04827 |
| DACT2~ENHO | 0.04829 |
| FCGR1A~MLIP | 0.04831 |
| NEXN~CYR61 | 0.04833 |
| CHIT1~ITGAX | 0.04834 |
| ATP8A2~SNHG9 | 0.04834 |
| FASN~TMIGD2 | 0.04835 |
| HCN2~PHACTR3 | 0.04836 |
| LINC01230~CRTAM | 0.04838 |
| CTLA4~APLNR | 0.0484 |
| CSRP2~GRIN2C | 0.04841 |
| WFDC1~MYOCD | 0.04842 |
| KCNK3~PRDM1 | 0.04843 |
| DHRS9~VNN1 | 0.04844 |
| TF~SPATA18 | 0.04845 |
| COL9A1~AL035446.1 | 0.04845 |
| CKMT2~PEBP4 | 0.04846 |
| CYP4B1~GFAP | 0.04847 |
| PRDM1~HOXA9 | 0.04847 |
| TF~FHOD3 | 0.04849 |
| BDH1~ADRB3 | 0.04849 |
| ADAM28~F2RL3 | 0.04851 |
| CYP4B1~AC025262.1 | 0.04852 |
| HMCN2~TMIGD2 | 0.04853 |
| TMIGD3~TFCP2L1 | 0.04854 |
| GFI1~ABCC8 | 0.04857 |
| LINC00942~TNMD | 0.04861 |
| TM4SF19~SPP1 | 0.04865 |
| GFI1~IL7R | 0.04866 |
| ADAM28~RCOR2 | 0.04866 |
| RGS2~ADAM28 | 0.04867 |
| AC004233.3~MT-ND3 | 0.04868 |
| FCGR2C~IGSF6 | 0.04869 |
| TF~MUC4 | 0.04869 |
| WFDC1~TOP2A | 0.04871 |
| SLAMF6~SGCA | 0.04872 |
| POTEF~AC069368.1 | 0.04876 |
| AL035446.1~GAS2L2 | 0.04877 |
| DPP10~LRRC73 | 0.04878 |

| | |
|---|---|
| KCNK3~ADGRE2 | 0.04882 |
| FCGR2C~CRTAM | 0.04883 |
| FCN2~TENM4 | 0.04886 |
| CXCR6~IL7R | 0.04887 |
| TRIM55~UBASH3B | 0.04887 |
| DMRT2~AC010997.5 | 0.04887 |
| ADAM28~CASS4 | 0.04891 |
| MSC~LY6K | 0.04891 |
| TRIM54~HMCN2 | 0.04893 |
| MIR4458HG~SOX10 | 0.04894 |
| TMIGD3~PHACTR3 | 0.04896 |
| VNN1~HRC | 0.049 |
| LRRC73~PDIA2 | 0.04901 |
| PKP3~HS3ST2 | 0.04902 |
| NRIP3~ITGAX | 0.04903 |
| CXCR4~PRDM1 | 0.04904 |
| CYP4B1~GDF15 | 0.04906 |
| CR1~GRIN2C | 0.04906 |
| MACROD1~AP000785.2 | 0.04907 |
| ATP8A2~IGSF1 | 0.04907 |
| DMRT2~AP000785.2 | 0.04908 |
| RCOR2~FHOD3 | 0.04911 |
| TUBB2A~CD300E | 0.04912 |
| SUSD1~IL2RA | 0.04912 |
| SLC4A8~AC092720.1 | 0.04913 |
| G0S2~RORB | 0.04914 |
| TRIM54~MAB21L1 | 0.04915 |
| AC110285.6~KCNB1 | 0.04915 |
| ERRFI1~SIK1 | 0.04918 |
| ERRFI1~CU639417.2 | 0.04919 |
| CKMT2~AC004233.3 | 0.04922 |
| C6~SGCA | 0.04923 |
| AL035446.1~STRC | 0.04923 |
| HSPA6~OLR1 | 0.04926 |
| CXCR4~CTLA4 | 0.04926 |
| F2RL3~TCF15 | 0.0493 |
| TUBB2A~TWIST1 | 0.04932 |
| SMIM1~PHACTR3 | 0.04933 |
| TF~BDH1 | 0.04935 |
| CTLA4~ITK | 0.04936 |
| TREML1~GPR65 | 0.04938 |
| IGFN1~LRRC73 | 0.04939 |
| CES1~TMIGD2 | 0.0494 |
| TRIM54~STRC | 0.04942 |
| AC004233.3~CCL22 | 0.04943 |
| TMEM52~ATP8A2 | 0.04944 |
| FAM181B~RERGL | 0.04948 |
| SPATA18~SLC6A12 | 0.04949 |

| | |
|---|---|
| LINC01230~AP000785.2 | 0.04949 |
| AC010997.5~HCN2 | 0.0495 |
| CSRNP1~GDF15 | 0.04951 |
| P2RY12~IGHD | 0.04954 |
| PTCHD4~NQO1 | 0.04955 |
| PRDM1~PHLDA1 | 0.04955 |
| PKP3~AP000785.2 | 0.04958 |
| CLDN9~WFDC1 | 0.04961 |
| LINC00942~IGSF1 | 0.04962 |
| GPR171~CRTAM | 0.04963 |
| ABCC8~ATP8A2 | 0.04966 |
| COL9A1~GDF1 | 0.04972 |
| ADRB3~IGHD | 0.04972 |
| CYP4B1~PTGDR2 | 0.04973 |
| KCNK3~ADGRL3 | 0.04973 |
| CYP4B1~PKP3 | 0.04974 |
| SPRR2F~LRRC73 | 0.04974 |
| CKMT2~CHCHD10 | 0.04977 |
| GLYCTK~C6 | 0.04978 |
| IGSF1~MT-ND2 | 0.04978 |
| TENM4~SIRPB1 | 0.04979 |
| SPATA18~ABCC3 | 0.04982 |
| CTLA4~CNFN | 0.04986 |
| OLR1~CD300E | 0.04986 |
| NCAPH~CLDN9 | 0.04987 |
| FCN2~TMIGD2 | 0.04988 |
| GFI1~SLAMF6 | 0.04989 |
| CHIT1~AKR1C3 | 0.04989 |
| AKR1B15~SLC4A8 | 0.0499 |
| CYP4B1~HOXA9 | 0.04991 |
| TRPA1~AC069368.1 | 0.04992 |
| ALDH4A1~LEP | 0.04993 |
| IGFN1~F8A2 | 0.04993 |
| FCGR1A~CA3 | 0.04996 |
| MUC20~TREML1 | 0.04996 |
| CHIT1~AC015688.5 | 0.04998 |
| RRM2~LRRC73 | 0.05 |
| POTEF~TREML1 | 0.05 |
| CHIT1~DCSTAMP | 0.05001 |
| ELFN1~EXOC3L2 | 0.05002 |
| AL035446.1~IL2RA | 0.05003 |
| ACTG2~SMOC1 | 0.05004 |
| LRRC73~TGFBR3L | 0.05004 |
| LINC01956~PEBP4 | 0.05005 |
| ABCC8~PTPRO | 0.05006 |
| NCAPH~THBS1 | 0.05008 |
| C6~AC005696.4 | 0.05008 |
| RRM2~MLIP | 0.05011 |

| | |
|---|---|
| PRDM1~TNFAIP3 | 0.05011 |
| SMIM1~SNHG9 | 0.05012 |
| ABCC8~PTGDR2 | 0.05013 |
| TRPA1~ADGRE2 | 0.05016 |
| LY6K~ALPK3 | 0.05016 |
| LINC00968~SRPX2 | 0.05018 |
| WFDC1~CNFN | 0.05018 |
| PRDM1~GDF15 | 0.05019 |
| COL9A1~CD300E | 0.05021 |
| SPATA18~TLR8 | 0.05022 |
| CYR61~SGK1 | 0.05024 |
| SPATA18~ADGRL3 | 0.05024 |
| CKMT2~REEP6 | 0.05025 |
| LINC00968~CDKN2B | 0.05027 |
| RASL11B~PDIA2 | 0.05029 |
| EXOC3L2~AL049634.2 | 0.0503 |
| FAT2~GRIN2C | 0.05031 |
| LINC01770~PTCHD4 | 0.05033 |
| LY6K~SPX | 0.05034 |
| IGFN1~C8orf82 | 0.05036 |
| SLAMF6~ITK | 0.05038 |
| DHRS9~AC004233.3 | 0.05038 |
| CD180~AC005696.4 | 0.05038 |
| IGFN1~TOP2A | 0.05039 |
| AL358334.2~CNTD2 | 0.05039 |
| CDKN1C~TMEM88 | 0.05043 |
| PTCHD4~IGSF6 | 0.05044 |
| CYR61~ID4 | 0.05046 |
| TREML1~GDF15 | 0.05046 |
| GABRD~LILRB3 | 0.05047 |
| SLC2A4~CYP2A6 | 0.05053 |
| DPP10~LUCAT1 | 0.05054 |
| AC010997.5~ABCC8 | 0.05056 |
| PDIA2~CNN1 | 0.05057 |
| SEMA6B~EXOC3L2 | 0.0506 |
| AL035446.1~NDRG4 | 0.05061 |
| TBXA2R~CNN1 | 0.05062 |
| CD28~HCN2 | 0.05066 |
| CCR5~CRTAM | 0.05066 |
| AL358334.2~AC005696.4 | 0.05066 |
| TRIM54~LIPA | 0.05067 |
| IGHD~CLDN9 | 0.05067 |
| EXOC3L2~SIRPB1 | 0.05067 |
| SLAMF6~SPATA18 | 0.0507 |
| SLAMF6~CXCR6 | 0.05073 |
| LINC01703~AL358334.2 | 0.05073 |
| LINC01956~HCN2 | 0.05073 |
| DHRS9~SUSD1 | 0.05073 |

| | |
|---|---|
| OLR1~CSRP2 | 0.05073 |
| GDF1~MATN4 | 0.05073 |
| NPL~CD28 | 0.05079 |
| MARCH1~HOXA9 | 0.05081 |
| CD28~VNN1 | 0.05082 |
| SOWAHA~MYOCD | 0.05082 |
| ADGRL3~FAT2 | 0.05085 |
| TREM2~OLR1 | 0.05086 |
| MT-CO2~MT-ND3 | 0.05087 |
| AC131097.2~GDF15 | 0.05088 |
| LY6K~OGDHL | 0.05088 |
| TOP2A~MATN4 | 0.05088 |
| SPRR2F~AC005696.4 | 0.0509 |
| TREML1~GALNT6 | 0.05091 |
| MUC4~C20orf204 | 0.05093 |
| LINC01703~PDIA2 | 0.05094 |
| FCGR2C~CD69 | 0.05095 |
| TM4SF19~MS4A6E | 0.05097 |
| RRM2~CRTAM | 0.05098 |
| TF~WFDC1 | 0.05098 |
| OGDHL~GSTZ1 | 0.051 |
| PSAT1~AC090181.1 | 0.05104 |
| GPR171~SOX10 | 0.05106 |
| TF~SOX10 | 0.05115 |
| LRRC73~TMIGD2 | 0.05115 |
| TRIM55~TBX1 | 0.05116 |
| SLC46A2~GFAP | 0.05116 |
| ADRB3~CIB2 | 0.05117 |
| DMRT2~GFAP | 0.05122 |
| CTLA4~FOS | 0.05127 |
| FCGR1A~LINC00942 | 0.05129 |
| TUBB2A~WFDC1 | 0.05135 |
| TF~FAT2 | 0.05136 |
| DHRS9~AC015688.5 | 0.0514 |
| NCAPH~CECR2 | 0.05146 |
| AL358334.2~GRIN2C | 0.05152 |
| DPP10~CKMT2 | 0.05154 |
| NEXN~F2RL3 | 0.05156 |
| SGCA~TBXA2R | 0.05156 |
| CERS1~MATN4 | 0.05156 |
| LRRC73~OGDHL | 0.05157 |
| CD300LB~CD300E | 0.05158 |
| CYR61~CU639417.2 | 0.05161 |
| COL9A1~PEBP4 | 0.05161 |
| CYR61~SIK1 | 0.05162 |
| MLIP~CSRP2 | 0.05163 |
| SPATA18~IPCEF1 | 0.05164 |
| CSRNP1~C5AR2 | 0.05166 |

| | |
|---|---|
| FAM181B~TBX1 | 0.05169 |
| TWIST1~HOXA9 | 0.05173 |
| IGFN1~CERS1 | 0.05175 |
| TF~MYOCD | 0.05175 |
| RASSF10~MAB21L1 | 0.05175 |
| SPRR2F~PHLDA1 | 0.05179 |
| FCMR~NRIP3 | 0.05179 |
| CRTAM~CASS4 | 0.0518 |
| SPRR2F~TRIM55 | 0.05182 |
| RORB~PDIA2 | 0.05182 |
| AP000785.2~IGSF6 | 0.05183 |
| AKR1B15~LINC00942 | 0.05185 |
| SLC6A12~GFAP | 0.05186 |
| TRIM54~RANBP3L | 0.05189 |
| SLAMF6~CCR5 | 0.05192 |
| TMIGD2~C20orf204 | 0.05192 |
| LRRC73~SIRPB1 | 0.05203 |
| FCGR2C~MUC20 | 0.05204 |
| AC092720.1~AC011498.4 | 0.05204 |
| AC005696.4~CYP2A6 | 0.05205 |
| HPCA~POTEF | 0.05207 |
| FCGR2B~HRC | 0.05208 |
| RCOR2~MATN4 | 0.05209 |
| PRR7~ELFN1 | 0.05211 |
| AKR1B15~PTGDR2 | 0.05214 |
| TRPA1~CCL22 | 0.05216 |
| RASL11B~AC005696.4 | 0.05224 |
| OGDHL~TOP2A | 0.05225 |
| TMEM88~TCF15 | 0.05226 |
| CMTM8~EXOC3L2 | 0.05228 |
| CHIT1~GAPT | 0.05229 |
| AC131097.2~AC010997.5 | 0.0523 |
| KANK4~CD300E | 0.05236 |
| NCAPH~CYP2A6 | 0.05236 |
| COL9A1~CERS1 | 0.05237 |
| GLIS1~ABCC8 | 0.05238 |
| MTRNR2L12~SPX | 0.05238 |
| FCN2~FASN | 0.05238 |
| GABRD~HRC | 0.05239 |
| DPP10~GCNT1 | 0.0524 |
| CXCR6~GPR171 | 0.0524 |
| ADGRL3~ABCC8 | 0.05241 |
| ARL4C~F13A1 | 0.05242 |
| MIR4458HG~MT-ND3 | 0.05242 |
| RORB~LINC00942 | 0.05244 |
| FHOD3~CNN1 | 0.05245 |
| CXCR4~CD69 | 0.05247 |
| RP11-592B15.9~CD300E | 0.05247 |

| | |
|---|---|
| LINC01770~MLIP | 0.05249 |
| MLIP~RASSF10 | 0.05251 |
| ADGRE2~MMP9 | 0.05252 |
| SPRR2F~LINC01956 | 0.05254 |
| LUCAT1~SNHG9 | 0.05257 |
| VNN1~FMN1 | 0.0526 |
| LUCAT1~AKR1C3 | 0.05262 |
| FAM78B~MATN4 | 0.05264 |
| MLIP~SMOC1 | 0.05264 |
| SPRR2F~CTLA4 | 0.05266 |
| HEY2~WFDC1 | 0.05267 |
| LY6K~AC107976.1 | 0.05268 |
| GRIN2C~HCN2 | 0.05268 |
| LINC01703~MUC4 | 0.05269 |
| ACTG2~HEY2 | 0.05269 |
| PROB1~SEMA6B | 0.05271 |
| HPCA~BDH1 | 0.05273 |
| MT-ND1~MT-CO2 | 0.05277 |
| ID4~HS3ST2 | 0.05279 |
| LY6K~HMCN2 | 0.05279 |
| PKP3~EXOC3L2 | 0.05279 |
| GAPT~IPCEF1 | 0.0528 |
| GFAP~GDF15 | 0.05284 |
| GLIS1~FCN2 | 0.05285 |
| KCNA2~BMP2 | 0.05289 |
| CDKN2B~HS3ST2 | 0.05289 |
| AL035446.1~GALNT6 | 0.0529 |
| MUC20~CNTD2 | 0.05293 |
| GABRD~LY6K | 0.05296 |
| ARL4C~SDS | 0.05296 |
| RGS2~ID4 | 0.05298 |
| IGFN1~SEMA6B | 0.05301 |
| CHIT1~TM4SF19 | 0.05302 |
| MUSTN1~CLDN9 | 0.05304 |
| IL10~ARL4C | 0.05305 |
| SPP1~AKR1B15 | 0.05307 |
| HSPA6~TIGAR | 0.05308 |
| SUSD1~SGCA | 0.05309 |
| IL7R~ITK | 0.05311 |
| ID4~PTCHD4 | 0.05312 |
| ADAM28~SDS | 0.05312 |
| LINC01230~FAM181B | 0.05312 |
| TUBB2A~MMP9 | 0.05318 |
| ADRB3~SH2D7 | 0.05319 |
| LINC00942~CCL22 | 0.0532 |
| PARP15~SNHG9 | 0.05323 |
| TMEM88~EXOC3L2 | 0.05325 |
| FCGR2C~TF | 0.05332 |

| | |
|---|---|
| CD28~F13A1 | 0.05337 |
| TMEM52~MUC4 | 0.05339 |
| APLNR~MYOCD | 0.05339 |
| GFAP~NRTN | 0.0534 |
| CSRP2~IGHD | 0.05343 |
| IGFN1~AC134669.2 | 0.05347 |
| MUSTN1~CNN1 | 0.05347 |
| ATP8A2~AC005696.4 | 0.05347 |
| CXCR4~CLEC4E | 0.05348 |
| AZGP1~AC011498.4 | 0.05354 |
| TREML1~NRIP3 | 0.05355 |
| BDH1~GCAT | 0.05361 |
| TREML1~LIPA | 0.05361 |
| PHLDA1~AL049634.2 | 0.05361 |
| TF~RCOR2 | 0.05364 |
| GAPT~ADGRE2 | 0.05366 |
| CD69~SIRPB1 | 0.05369 |
| AC010997.5~CCL22 | 0.05371 |
| SDS~TCF15 | 0.05371 |
| PLEK~GRIN2C | 0.05373 |
| RANBP3L~TMIGD2 | 0.05373 |
| IL2RA~AC004233.3 | 0.05378 |
| GABRD~FCGR3A | 0.05382 |
| G0S2~CARMN | 0.05389 |
| TF~TENM4 | 0.0539 |
| STRC~AC110285.6 | 0.0539 |
| ARL4C~OSBPL3 | 0.05399 |
| WFDC1~SGCA | 0.05399 |
| ADGRL3~TBX1 | 0.054 |
| IL1RN~LINC00942 | 0.05404 |
| TFCP2L1~PARP15 | 0.05407 |
| TREML1~SMIM25 | 0.05409 |
| FCN2~HS3ST2 | 0.05411 |
| RORB~SOX10 | 0.05414 |
| SPP1~PLA2G7 | 0.05419 |
| KLHL6~SGCA | 0.05423 |
| CCL18~OXT | 0.05425 |
| FCGR2C~RP11-592B15.9 | 0.05428 |
| MSC~CD69 | 0.05428 |
| AL035446.1~LINC00942 | 0.05432 |
| FAM78B~ADGRL3 | 0.05433 |
| ADAM28~OGDHL | 0.05438 |
| HMCN2~TBX1 | 0.05439 |
| IGFN1~CLDN9 | 0.05441 |
| MTRNR2L12~MT-CO3 | 0.05441 |
| WFDC1~FOXS1 | 0.05444 |
| MUC20~RASSF10 | 0.05446 |
| MTRNR2L12~MT-ND4 | 0.0545 |

| | |
|---|---|
| IGFN1~TRIM54 | 0.05451 |
| FCN2~SGCA | 0.05453 |
| TRIM54~GSTZ1 | 0.05454 |
| SPX~ALPK3 | 0.05454 |
| DHRS9~NRIP3 | 0.05457 |
| FOSL2~CDKN1A | 0.05459 |
| RASL11B~DAPP1 | 0.0546 |
| AC131097.2~OLR1 | 0.05461 |
| MIR4458HG~SNHG9 | 0.05468 |
| MYOCD~NRTN | 0.05469 |
| FCN2~CCL18 | 0.05472 |
| ITK~IPCEF1 | 0.05474 |
| SUSD1~SIRPB1 | 0.05474 |
| F2RL3~IGSF1 | 0.05475 |
| TBXA2R~EXOC3L2 | 0.05476 |
| IL10~AC005696.4 | 0.05477 |
| ABCC8~SMOC1 | 0.05478 |
| TMEM52~MUC20 | 0.05493 |
| ADGRL3~PSAT1 | 0.05494 |
| WFDC1~CCR10 | 0.05494 |
| MTRNR2L12~SNHG9 | 0.05496 |
| TF~REEP6 | 0.05496 |
| CCL22~HRC | 0.05497 |
| LY6K~AP000785.2 | 0.05503 |
| PEBP4~GFAP | 0.05506 |
| AL035446.1~AP000785.2 | 0.05512 |
| CARMN~AC110285.6 | 0.05518 |
| ATP8A2~SOX10 | 0.05518 |
| MLIP~MAB21L1 | 0.05519 |
| HSPA6~ADAM28 | 0.0552 |
| PTCHD4~NXNL1 | 0.05521 |
| CARMN~PDIA2 | 0.05522 |
| SPATA18~LYZ | 0.05525 |
| CR1~F13A1 | 0.05528 |
| TMIGD3~P2RY12 | 0.05532 |
| POTEF~MLIP | 0.05533 |
| SPP1~ALOX15B | 0.05535 |
| CYP4B1~TREML1 | 0.05538 |
| BDH1~CLDN9 | 0.05543 |
| PTGDR2~FHOD3 | 0.05543 |
| TRIM54~TBXA2R | 0.05544 |
| CD28~SGCA | 0.05546 |
| SPP1~OLR1 | 0.05546 |
| BDH1~TRIM55 | 0.05548 |
| AKR1B15~SUSD1 | 0.0555 |
| AC107976.1~AC110285.6 | 0.0555 |
| TF~VNN1 | 0.05551 |
| UCHL1~SRPX2 | 0.05552 |

| | |
|---|---|
| FOS~GDF15 | 0.05553 |
| CLEC4E~F2RL3 | 0.05555 |
| MYCL~Z99774.1 | 0.05556 |
| VNN1~TRPA1 | 0.05559 |
| COL9A1~SGCA | 0.05562 |
| MUC4~TREML1 | 0.05566 |
| TRPA1~EXOC3L2 | 0.0557 |
| RASL11B~UBASH3B | 0.05572 |
| AKR1B15~CES1 | 0.05575 |
| PTPN22~SGCA | 0.05576 |
| PDIA2~AC110285.6 | 0.05576 |
| CSRNP1~SGK1 | 0.05577 |
| SPP1~TREM2 | 0.0558 |
| FCN2~GLYAT | 0.05583 |
| AC131097.2~FASN | 0.05584 |
| GFI1~AL358334.2 | 0.05586 |
| SPRR2F~TOP2A | 0.05592 |
| CES1~NQO1 | 0.05595 |
| FCN2~CES1 | 0.05604 |
| RASSF10~IGSF1 | 0.05607 |
| CLEC4E~PHLDA1 | 0.05607 |
| CA3~CCL18 | 0.05608 |
| HEY2~HRC | 0.05611 |
| FOSL2~CSRNP1 | 0.05612 |
| HS3ST2~AC015688.5 | 0.05612 |
| RRM2~AL049634.2 | 0.05615 |
| CDKN1C~AC092720.1 | 0.05617 |
| SGCA~GDF15 | 0.05617 |
| MUC20~MLIP | 0.05619 |
| GPR171~CD300E | 0.05622 |
| CES1~GAS2L2 | 0.05626 |
| KCNK3~HCN2 | 0.05627 |
| ENHO~AC004233.3 | 0.05627 |
| SPRR2F~CNN1 | 0.05628 |
| TUBB2A~RP11-592B15.9 | 0.05631 |
| HSPA6~PKP3 | 0.05633 |
| LINC01703~ADRB3 | 0.05633 |
| MERTK~FKBP5 | 0.05636 |
| LRRC73~MAPK8IP2 | 0.0564 |
| RANBP3L~TNMD | 0.05642 |
| PKP3~PDIA2 | 0.05643 |
| CXCR6~CCR5 | 0.05647 |
| ID4~TGFBR3L | 0.05652 |
| GJC3~AZGP1 | 0.05653 |
| CKMT2~LDHD | 0.05659 |
| MTRNR2L12~MT-ATP6 | 0.05661 |
| LINC01703~GDF15 | 0.05662 |
| SLAMF6~CRTAM | 0.05664 |

| | |
|---|---|
| NEXN~ATP8A2 | 0.05666 |
| HS3ST2~RASL10B | 0.05667 |
| PTCHD4~OXT | 0.05668 |
| PRDM1~F2RL3 | 0.05669 |
| TRIM54~HCN2 | 0.0567 |
| GLIS1~CYP2A6 | 0.0568 |
| IL10~CD180 | 0.05683 |
| WFDC1~TBXA2R | 0.05683 |
| AC005696.4~MATN4 | 0.05683 |
| GLYAT~CYP2A6 | 0.05685 |
| MUSTN1~HEY2 | 0.05687 |
| VNN1~SGCA | 0.05688 |
| PDIA2~KCNB1 | 0.05691 |
| C5AR1~C5AR2 | 0.05696 |
| TWIST1~CA3 | 0.05699 |
| MT-ND2~MT-CO2 | 0.05699 |
| GSTZ1~AC005696.4 | 0.057 |
| NEXN~RASL11B | 0.05701 |
| LINC01956~HOXA9 | 0.05707 |
| SUSD1~LINC00942 | 0.05707 |
| AL035406.1~AC015688.5 | 0.0571 |
| IGHD~GCAT | 0.0571 |
| AL358334.2~GDF15 | 0.05712 |
| GABRD~IGFN1 | 0.05715 |
| HSPA6~THBS1 | 0.05716 |
| MT-CO2~MT-CYB | 0.0572 |
| SPATA18~GAS2L2 | 0.05724 |
| TF~CA3 | 0.05729 |
| LINC01703~RORB | 0.05732 |
| ABCC8~CSRP2 | 0.05736 |
| HS3ST2~ANKRD24 | 0.05738 |
| RRM2~PTCHD4 | 0.0574 |
| TUBB2A~TREML1 | 0.05742 |
| CSRP2~C20orf204 | 0.05742 |
| LDHD~SLC2A4 | 0.05744 |
| RGS1~CTLA4 | 0.05745 |
| KCNK3~MYOCD | 0.05748 |
| CDKN1A~GDF15 | 0.05755 |
| LINC00942~GAS2L2 | 0.05755 |
| FKBP5~RASL10B | 0.05759 |
| RRM2~NRTN | 0.05761 |
| AC005696.4~GRIN2C | 0.05768 |
| NCAPH~CD300LB | 0.0578 |
| ACP5~GDF15 | 0.05781 |
| CHIT1~ADGRE2 | 0.05784 |
| MYCL~MSC | 0.05785 |
| FCGR2C~TENM4 | 0.05789 |
| IGFN1~WFDC1 | 0.0579 |

| | |
|---|---|
| MT-ND1~MT-ND3 | 0.05792 |
| ATP8A2~TOP2A | 0.05793 |
| CLDN9~NRTN | 0.05796 |
| SPRR2F~HMCN2 | 0.058 |
| LINC01094~CLEC7A | 0.05809 |
| GFAP~SOX10 | 0.05811 |
| LINC00968~SOX10 | 0.05812 |
| HSPA6~LINC01956 | 0.05813 |
| DRD4~AC005696.4 | 0.05817 |
| LINC00968~SLC46A2 | 0.05818 |
| AC092720.1~CYP2A6 | 0.05819 |
| TRIM54~CNN1 | 0.05821 |
| TFCP2L1~FKBP5 | 0.05823 |
| IL10~AC131097.2 | 0.05824 |
| SLC27A2~RASL10B | 0.05824 |
| ENHO~GAS2L2 | 0.05837 |
| C6~RASL10B | 0.05842 |
| TMIGD3~HSPA6 | 0.05848 |
| SOWAHA~AL358334.2 | 0.05848 |
| CCL18~LILRB3 | 0.05849 |
| FCGR1A~SOWAHA | 0.05851 |
| ACTG2~TF | 0.05851 |
| CCL22~AC110285.6 | 0.05859 |
| ADGRL3~SGCA | 0.05864 |
| KCNK3~NRTN | 0.05868 |
| IGFN1~OGDHL | 0.05874 |
| TTC36~PHACTR3 | 0.05877 |
| CSRNP1~AL358334.2 | 0.05885 |
| MLIP~RORB | 0.05886 |
| RORB~IGSF1 | 0.05901 |
| CTLA4~ADRB3 | 0.05911 |
| AC131097.2~CECR2 | 0.05911 |
| BIRC3~SNHG9 | 0.05915 |
| MSC~TRPA1 | 0.05935 |
| IGHD~HCN2 | 0.05936 |
| MT-ND2~MT-ND3 | 0.05937 |
| CYR61~IL10 | 0.0594 |
| IL1RN~MUC4 | 0.05944 |
| GABRD~WFDC1 | 0.05945 |
| DUSP1~CU639417.2 | 0.05947 |
| DUSP1~SIK1 | 0.05947 |
| FCGR2C~CLEC7A | 0.0595 |
| MT-ND1~MT-CO3 | 0.05952 |
| TF~AC010997.5 | 0.05959 |
| DPP10~AC131097.2 | 0.05967 |
| MLIP~LY6K | 0.05967 |
| OLR1~F2RL3 | 0.05967 |
| TRIM55~SLC27A2 | 0.05979 |

| | |
|---|---|
| ACTG2~AL049634.2 | 0.05982 |
| RASL11B~ADAM12 | 0.05982 |
| OPLAH~CLDN9 | 0.05984 |
| TRIM55~AC010997.5 | 0.05986 |
| GAPT~CASS4 | 0.05998 |
| CKMT2~PSAT1 | 0.05999 |
| CHIT1~PHACTR3 | 0.06008 |
| G0S2~DMRT2 | 0.06015 |
| AC107976.1~TBX1 | 0.06016 |
| IL10~FOS | 0.06022 |
| HOXA9~SMOC1 | 0.06023 |
| AZGP1~SGCA | 0.06035 |
| IL1RN~IGHD | 0.06047 |
| MT-CO3~MT-CYB | 0.06047 |
| SOWAHA~IGHD | 0.06053 |
| FAT2~PSAT1 | 0.06058 |
| CRTAM~TMIGD2 | 0.0606 |
| MLIP~HMCN2 | 0.06061 |
| IGFN1~GDF1 | 0.06068 |
| TRIM54~ADGRL3 | 0.06072 |
| IL1RN~TM4SF19 | 0.06074 |
| PARP15~PTCHD4 | 0.06075 |
| COL9A1~ADRB3 | 0.06079 |
| HMCN2~CCL22 | 0.06091 |
| CD52~GAPT | 0.06094 |
| TMEM52~LY6K | 0.06097 |
| MUC20~PDIA2 | 0.061 |
| LINC01230~PKP3 | 0.06101 |
| AC131097.2~GPR65 | 0.0611 |
| RERGL~TBX1 | 0.06118 |
| MT-ND2~MT-CO3 | 0.06119 |
| POTEF~HMCN2 | 0.06129 |
| HTRA3~RCOR2 | 0.06129 |
| CYR61~CSRNP1 | 0.06141 |
| ADGRE2~SIRPB1 | 0.06145 |
| FAM181B~ART4 | 0.06146 |
| CYR61~GDF15 | 0.06152 |
| TF~PKP3 | 0.06152 |
| APLNR~TCF15 | 0.06152 |
| TRIM55~OLR1 | 0.06155 |
| TF~SOWAHA | 0.06162 |
| CLEC7A~FOS | 0.06162 |
| AC131097.2~BDH1 | 0.06163 |
| IGFN1~ADGRL3 | 0.06171 |
| KCNK3~CKMT2 | 0.06171 |
| CLEC7A~SIRPB1 | 0.06171 |
| AC134669.2~EVI2A | 0.06171 |
| MT-CO2~MT-ND4 | 0.06179 |

| | |
|---|---|
| NEXN~KCNB1 | 0.06181 |
| MIR4458HG~PKP3 | 0.06185 |
| LINC01770~FCN2 | 0.0619 |
| VNN1~FCN2 | 0.06191 |
| IGFN1~LY6K | 0.06199 |
| THEMIS2~ARL4C | 0.06203 |
| TFCP2L1~AC110285.6 | 0.0621 |
| HOXA9~ADGRE2 | 0.06211 |
| HEY2~CNN1 | 0.06216 |
| ID4~AC092720.1 | 0.06224 |
| IL10~C5AR2 | 0.06243 |
| ACTG2~SOX10 | 0.06245 |
| G0S2~LINC01230 | 0.06249 |
| PROB1~AC092720.1 | 0.06252 |
| TMEM88~SEMA6B | 0.06257 |
| OGDHL~HS3ST2 | 0.06261 |
| MYOCD~FHOD3 | 0.06262 |
| HOXA9~ADRB3 | 0.06264 |
| SLAMF6~GPR171 | 0.06265 |
| CD69~CLEC7A | 0.06267 |
| CARMN~STRC | 0.06277 |
| HTRA3~HMCN2 | 0.0628 |
| TNFAIP3~TRPA1 | 0.0628 |
| KCNK3~PEBP4 | 0.06284 |
| RASL11B~CNN1 | 0.06285 |
| MT-CO2~MT-ATP6 | 0.06285 |
| CA3~DMRT2 | 0.06289 |
| TREML1~TREM2 | 0.06293 |
| FHOD3~MATN4 | 0.06297 |
| TMIGD3~CLDN9 | 0.06309 |
| AP000785.2~CD69 | 0.06316 |
| GABRD~COL9A1 | 0.06321 |
| MT-CO3~MT-ND4 | 0.06321 |
| SPP1~ABCC8 | 0.06324 |
| CSRNP1~FOS | 0.06325 |
| SOWAHA~CNTD2 | 0.06327 |
| THBS1~AC004233.3 | 0.06327 |
| TNFAIP3~CLDN9 | 0.06328 |
| LIPA~NRIP3 | 0.06329 |
| NEXN~LINC01956 | 0.06333 |
| MAB21L1~CECR2 | 0.06335 |
| TMIGD3~ADORA3 | 0.06336 |
| CD300E~LILRB3 | 0.06337 |
| HSPA6~IL10 | 0.06339 |
| MIR4458HG~ATP8A2 | 0.06347 |
| IGFN1~FGF22 | 0.06354 |
| COL9A1~F2RL3 | 0.06357 |
| CTLA4~SOX10 | 0.06361 |

| | |
|---|---|
| RERGL~HRC | 0.06367 |
| PKP3~GAS2L2 | 0.06376 |
| PRR7~SEMA6B | 0.06399 |
| ADRB3~AC004233.3 | 0.06399 |
| MUSTN1~WFDC1 | 0.06401 |
| CSRNP1~DUSP1 | 0.06407 |
| KCNK3~REEP6 | 0.0641 |
| HSPA6~MUC20 | 0.06419 |
| ARL4C~HS3ST2 | 0.0642 |
| AC092720.1~AL121845.2 | 0.0642 |
| IGHD~CYP2A6 | 0.06429 |
| GFI1~CXCR6 | 0.0644 |
| MT-CO2~MT-CO3 | 0.06451 |
| AL080251.1~ALDH4A1 | 0.06452 |
| IGFN1~MIR4458HG | 0.06454 |
| CYP4B1~FKBP5 | 0.06455 |
| LY6K~WFDC1 | 0.06455 |
| MT-ND1~MT-CYB | 0.06461 |
| BDH1~VNN1 | 0.06465 |
| LINC00942~HS3ST2 | 0.0647 |
| SGCA~CD300E | 0.06473 |
| HEY2~TBXA2R | 0.06484 |
| RASL11B~FCN2 | 0.06486 |
| CYR61~ADRB3 | 0.06494 |
| DUSP1~PHACTR3 | 0.06494 |
| FKBP5~KCNB1 | 0.06499 |
| CDKN1A~CU639417.2 | 0.06502 |
| CDKN1A~SIK1 | 0.06503 |
| CTLA4~ATP8A2 | 0.06508 |
| AC131097.2~RASL10B | 0.06522 |
| DPP10~MAPK8IP2 | 0.06534 |
| PKP3~SH2D7 | 0.0654 |
| AKR1B15~LIPA | 0.06551 |
| PHACTR3~IGSF1 | 0.06552 |
| CYP4B1~RANBP3L | 0.06554 |
| SPATA18~GDF15 | 0.06562 |
| APLNR~HRC | 0.06567 |
| IL24~FCMR | 0.06575 |
| GAS2L2~TMIGD2 | 0.06576 |
| CXCR4~F2RL3 | 0.0658 |
| GFAP~PHACTR3 | 0.0658 |
| LINC00942~AC090181.1 | 0.06595 |
| CYP4B1~NEXN | 0.06596 |
| HMCN2~AL049634.2 | 0.066 |
| TRIM54~PKP3 | 0.0661 |
| PKP3~CD300E | 0.06622 |
| PDIA2~GAS2L2 | 0.06629 |
| TF~LUCAT1 | 0.06634 |

| | |
|---|---|
| AZGP1~PLIN5 | 0.06636 |
| FCN2~LDHD | 0.06645 |
| TMIGD3~RANBP3L | 0.06649 |
| ACTG2~MYOCD | 0.06651 |
| RASL11B~ID4 | 0.06654 |
| IGFN1~AL049634.2 | 0.0666 |
| COL9A1~RERGL | 0.06682 |
| MT-ND1~MT-ATP6 | 0.06684 |
| IGFN1~HRC | 0.06686 |
| MT-ND1~MT-ND4 | 0.06687 |
| SPRR2F~HOXA9 | 0.06688 |
| AL035446.1~ANKRD24 | 0.06694 |
| GRIN2C~SOX10 | 0.06699 |
| WFDC1~AC005696.4 | 0.06701 |
| SPRR2F~OGDHL | 0.06702 |
| ENHO~HMCN2 | 0.06721 |
| LINC01230~GFAP | 0.06723 |
| SPP1~CCL22 | 0.06736 |
| CD52~ADGRE2 | 0.06747 |
| G0S2~CRTAM | 0.06749 |
| RASL11B~ATP8A2 | 0.06751 |
| ABCC8~AC005696.4 | 0.06759 |
| SPP1~ATP8A2 | 0.06769 |
| LRRC73~AL049634.2 | 0.06776 |
| MUSTN1~MYOCD | 0.06781 |
| TNMD~SRPX2 | 0.06788 |
| SGCA~LILRB3 | 0.0679 |
| HSPA6~CLDN9 | 0.06792 |
| MUC4~IGHD | 0.06792 |
| RORB~ATP8A2 | 0.06792 |
| LINC01703~MUC20 | 0.06799 |
| FCGR2B~PKP3 | 0.06803 |
| ENHO~CD300E | 0.06817 |
| HSPA6~CD180 | 0.06818 |
| TUBB2A~MATN4 | 0.06824 |
| POTEF~TUBB2A | 0.06828 |
| AKR1B15~AL358334.2 | 0.06831 |
| RERGL~MYOCD | 0.06831 |
| NRIP3~Z99774.1 | 0.06834 |
| AC004233.3~WFDC1 | 0.0684 |
| PTCHD4~F2RL3 | 0.06842 |
| MUC4~OXT | 0.06881 |
| TF~TRPA1 | 0.06884 |
| PLA2G7~MMP9 | 0.06913 |
| LINC01703~MLIP | 0.0694 |
| ITGAX~ADGRE2 | 0.06946 |
| OGDHL~STRC | 0.06958 |
| TRIM54~NRIP3 | 0.06961 |

| | |
|---|---|
| AL035446.1~SIRPB1 | 0.06961 |
| ART4~AL358334.2 | 0.06976 |
| PDIA2~SIRPB1 | 0.06987 |
| NRIP3~AC005696.4 | 0.06988 |
| HS3ST2~GDF15 | 0.06988 |
| MT-ATP6~MT-CO3 | 0.06989 |
| IGFN1~HMCN2 | 0.06997 |
| MT-ND4~MT-CYB | 0.06997 |
| TM4SF19~MMP9 | 0.07012 |
| HPCA~SGCA | 0.07014 |
| MT-ATP6~MT-CYB | 0.07022 |
| HTRA3~OXT | 0.07031 |
| UBASH3B~ADGRE2 | 0.07036 |
| MT-ND2~MT-ATP6 | 0.0704 |
| DCSTAMP~MMP9 | 0.07053 |
| PKP3~OLR1 | 0.07055 |
| TREML1~SOX10 | 0.07062 |
| RGS1~DUSP1 | 0.07064 |
| COL9A1~SLC27A2 | 0.07073 |
| LINC00968~TNMD | 0.07082 |
| RGS1~FOS | 0.07083 |
| COL9A1~MYOCD | 0.07113 |
| CD69~GPR183 | 0.07119 |
| IGFN1~PROB1 | 0.07139 |
| AC068987.5~CYP2A6 | 0.0714 |
| UCHL1~NQO1 | 0.07142 |
| PTCHD4~GALNT6 | 0.07157 |
| LINC01956~SGCA | 0.07169 |
| NRIP3~TGFBR3L | 0.07169 |
| LY6K~ATP8A2 | 0.07182 |
| ADGRL3~MLIP | 0.07204 |
| ATP8A2~CCL22 | 0.07209 |
| LY6K~CCL22 | 0.07212 |
| COL9A1~SRPX2 | 0.07216 |
| FCGR2C~LINC01094 | 0.07228 |
| SPATA18~MSC | 0.0723 |
| LINC01956~TMIGD2 | 0.0724 |
| HPCA~MSC | 0.0725 |
| MT-ND2~MT-CYB | 0.07258 |
| ADRB3~SLC27A2 | 0.07268 |
| GJC3~ABCC8 | 0.0727 |
| F13A1~AC092720.1 | 0.07302 |
| SOWAHA~IGSF1 | 0.07307 |
| TRPA1~ATP8A2 | 0.07333 |
| MT-ND2~MT-ND4 | 0.07334 |
| HSPA6~SAMSN1 | 0.07362 |
| AC004233.3~EXOC3L2 | 0.07397 |
| PKP3~CCL22 | 0.07398 |

| | |
|---|---|
| COL9A1~PHACTR3 | 0.07418 |
| RERGL~WFDC1 | 0.07432 |
| MYCL~ATP8A2 | 0.07434 |
| TRIM54~HEY2 | 0.07444 |
| COL9A1~HRC | 0.07461 |
| NEXN~FKBP5 | 0.07477 |
| CHIT1~SPP1 | 0.07513 |
| CSRNP1~CU639417.2 | 0.07555 |
| CSRNP1~SIK1 | 0.07556 |
| TF~GFAP | 0.07556 |
| SPATA18~NRIP3 | 0.07557 |
| SPX~IGHD | 0.07561 |
| IGFN1~DHRS9 | 0.07567 |
| TRPA1~CD69 | 0.07601 |
| CYR61~DUSP1 | 0.07605 |
| TF~CLDN9 | 0.07612 |
| LINC01956~AKR1B15 | 0.07622 |
| MT-ATP6~MT-ND4 | 0.07635 |
| MT-ND1~MT-ND2 | 0.07646 |
| F2RL3~SIK1 | 0.07648 |
| F2RL3~CU639417.2 | 0.0765 |
| CD28~IGHD | 0.07655 |
| RORB~SPX | 0.0766 |
| PTCHD4~AC110285.6 | 0.07663 |
| FHOD3~CYP2A6 | 0.07666 |
| ACTG2~RORB | 0.07676 |
| HMCN2~GAS2L2 | 0.07679 |
| RANBP3L~LINC00968 | 0.07682 |
| STRC~NDRG4 | 0.07684 |
| LY6K~SH2D7 | 0.0771 |
| IPCEF1~ABCC8 | 0.07712 |
| AC090181.1~AC004233.3 | 0.07733 |
| MUC20~IGHD | 0.07745 |
| MTRNR2L12~MIR4458HG | 0.07752 |
| DDTL~DDT | 0.07786 |
| ABCC8~GRIN2C | 0.07791 |
| IGHD~HS3ST2 | 0.07798 |
| FCGR2B~HMCN2 | 0.07805 |
| PLIN5~AC011498.4 | 0.07844 |
| GABRD~GRIN2C | 0.07846 |
| PSAT1~IGSF1 | 0.07858 |
| ACTG2~WFDC1 | 0.07859 |
| LINC01956~AC131097.2 | 0.07863 |
| HMCN2~SNHG9 | 0.07871 |
| AC005696.4~AL049634.2 | 0.07905 |
| AC004233.3~NRTN | 0.07921 |
| RASL11B~CLEC4E | 0.07931 |
| FCGR1A~ADGRE2 | 0.07956 |

| | |
|---|---|
| FCGR1A~UBASH3B | 0.07971 |
| LY6K~CIB2 | 0.07982 |
| SUSD1~SNHG9 | 0.07986 |
| MUC4~STRC | 0.08 |
| PKP3~AC107976.1 | 0.08021 |
| ATP8A2~SGCA | 0.08029 |
| FOS~PHACTR3 | 0.08035 |
| PTCHD4~TMIGD2 | 0.08038 |
| UBASH3B~AC092720.1 | 0.0805 |
| PTGDR2~CSRP2 | 0.08069 |
| FAM78B~PKP3 | 0.08071 |
| ABCC8~SGCA | 0.08082 |
| MTRNR2L12~LINC00942 | 0.08101 |
| TNFAIP3~BIRC3 | 0.08108 |
| DUSP1~CD69 | 0.08115 |
| LINC01956~IGHD | 0.08116 |
| IGFN1~RERGL | 0.08163 |
| ACTG2~MUSTN1 | 0.082 |
| SPRR2F~VNN1 | 0.08204 |
| TRIM54~NDRG4 | 0.08213 |
| EXOC3L2~TCF15 | 0.08269 |
| CA3~LINC01230 | 0.0828 |
| RGS1~CD69 | 0.08335 |
| SPRR2F~RASL11B | 0.08336 |
| APLNR~EXOC3L2 | 0.08404 |
| CSRNP1~CDKN1A | 0.08412 |
| TRPA1~LY6K | 0.08432 |
| PKP3~TENM4 | 0.08483 |
| IL1RN~CCL22 | 0.08504 |
| TRIM54~TBX1 | 0.08514 |
| TREML1~ADAM28 | 0.08518 |
| LINC00942~SPX | 0.08519 |
| OSBPL3~CLDN9 | 0.08529 |
| TM4SF19~PLA2G7 | 0.08542 |
| TREML1~UBASH3B | 0.08553 |
| GAPT~ADAM28 | 0.08556 |
| RASL11B~TWIST1 | 0.08567 |
| MUC20~PKP3 | 0.08591 |
| SPRR2F~PARP15 | 0.08628 |
| TF~LINC01094 | 0.08727 |
| PTCHD4~PHACTR3 | 0.08785 |
| PRDM1~CD69 | 0.08804 |
| ACTG2~ADGRL3 | 0.0881 |
| CSRP2~AC005696.4 | 0.08813 |
| OGDHL~IGHD | 0.0884 |
| PDIA2~AC005696.4 | 0.0887 |
| KCNK3~HS3ST2 | 0.08913 |
| RRM2~TOP2A | 0.08961 |

| | |
|---|---|
| MUC4~CNTD2 | 0.0898 |
| DPP10~OGDHL | 0.0901 |
| LINC01956~RASL11B | 0.09014 |
| MUSTN1~ADGRL3 | 0.09042 |
| CD300E~ADGRE2 | 0.09106 |
| HMCN2~AC005696.4 | 0.09124 |
| CYP4B1~RASL10B | 0.09128 |
| HS3ST2~PHACTR3 | 0.09157 |
| HTRA3~MATN4 | 0.09227 |
| IGFN1~TBX1 | 0.09249 |
| MYOCD~CNN1 | 0.0926 |
| NRIP3~TMIGD2 | 0.09326 |
| SOWAHA~CYP2A6 | 0.09567 |
| CLDN9~PHACTR3 | 0.09759 |
| CLEC7A~IGHD | 0.09799 |
| AP000785.2~AC005696.4 | 0.09825 |
| OGDHL~MAPK8IP2 | 0.09949 |
| SIRPB1~PHACTR3 | 0.09959 |
| CD69~FOS | 0.10077 |
| AC010997.5~AC110285.6 | 0.10147 |
| SPATA18~PKP3 | 0.1026 |
| CA3~FCN2 | 0.10558 |
| SPRR2F~DPP10 | 0.10652 |
| TF~AC004233.3 | 0.10669 |
| SPRR2F~ABCC8 | 0.10725 |
| DACT2~AC005696.4 | 0.10729 |
| HS3ST2~AC005696.4 | 0.10823 |
| PDIA2~PHACTR3 | 0.11067 |
| WFDC1~CNN1 | 0.11407 |
| CYR61~FOS | 0.11424 |
| SH2D7~CIB2 | 0.11852 |
| TRIM54~ACTG2 | 0.11911 |
| ADGRL3~MYOCD | 0.12437 |
| TF~IGHD | 0.12615 |
| DUSP1~FOS | 0.13615 |
| SIK1~CU639417.2 | 0.14249 |
| ACTG2~CNN1 | 0.15658 |
| LINC01956~ATP8A2 | 0.17982 |
| GDF1~CERS1 | 0.18616 |
| AC004233.3~CLDN9 | 0.19225 |
| AL049634.2~SIRPB1 | 0.21884 |
| LINC01230~DMRT2 | 0.2845 |
| MUC20~MUC4 | 0.31048 |

Table S3. Output from Centiscape showing network centrality measures for network constructed in Genenet.

| name |
|---|
| IGFN1 |
| IGHD |
| PHACTR3 |
| PKP3 |
| CLDN9 |
| SGCA |
| TRIM54 |
| GABRD |
| AC005696.4 |
| CYP4B1 |
| TF |
| ATP8A2 |
| HS3ST2 |
| AC004233.3 |
| SPATA18 |
| SPRR2F |
| LINC01956 |
| CYP2A6 |
| HSPA6 |
| PTCHD4 |
| RASL11B |
| ABCC8 |
| TREML1 |
| NRIP3 |
| AC131097.2 |
| WFDC1 |
| IGSF1 |
| LINC00942 |
| DPP10 |
| PDIA2 |
| AC092720.1 |
| CRTAM |
| AL358334.2 |
| GDF15 |
| F13A1 |
| MLIP |
| HOXA9 |
| CD300E |
| VNN1 |
| SNHG9 |
| CD69 |
| FCN2 |
| MTRNR2L12 |

SOX10
ADAM28
ADGRL3
LY6K
SOWAHA
HMCN2
POTEF
EXOC3L2
AL035446.1
KCNK3
CA3
TMIGD2
MIR4458HG
GRIN2C
NCAPH
GAS2L2
AC010997.5
GFAP
ADRB3
F2RL3
CCL22
MUC4
AP000785.2
CSRP2
TRPA1
FCGR2C
STRC
MT-ND3
AL049634.2
LINC01230
FKBP5
CLEC4E
LRRC73
CTLA4
PTGDR2
CHIT1
AKR1B15
COL9A1
ALDH4A1
FCGR1A
UBASH3B
SLC2A4
FOS
CD28
MATN4

CKMT2
GLYAT
HCN2
SUSD1
FCGR3A
SMIM25
ENHO
RCOR2
CSRNP1
RASL10B
GAPT
ARL4C
TFCP2L1
PRDM1
SPX
OLR1
MSC
CYR61
PHLDA1
SIRPB1
TRIM55
OGDHL
ID4
TBX1
MAB21L1
RANBP3L
DUSP1
C6
LINC01703
TUBB2A
SPP1
FAM181B
RRM2
CNN1
PSAT1
HRC
MUC20
FAT2
ADGRE2
THBS1
RERGL
ACTG2
MYOCD
NRTN
HTRA3

HEY2
CECR2
DACT2
LDHD
NEXN
AC110285.6
SLC6A12
OXT
SLAMF6
GDF1
TMEM88
CERS1
KCNA2
DMRT2
SDS
RASSF10
TWIST1
BMP2
GPR171
CNFN
ALPK3
IL10
UCHL1
G0S2
ANKRD24
RORB
TENM4
MT-ATP6
P2RY12
FHOD3
TGFBR3L
TIGAR
KCNB1
IPCEF1
LINC01770
TMIGD3
MT-CO2
CXCR4
Z99774.1
PEBP4
SMOC1
AC107976.1
CCL18
BDH1
SLC27A2

SLC46A2
CES1
FAM78B
LILRB3
ELFN1
DHRS9
TBXA2R
MMP9
HPCA
GFI1
APLNR
IGSF6
LUCAT1
ITK
MYCL
BIRC3
RP11-592B15.9
AC015688.5
FCGR2B
SLC4A8
TCF15
DCSTAMP
C20orf204
TM4SF19
FCMR
MT-CYB
PLA2G7
IL1RN
MT-ND2
AKR1C3
TNFAIP3
IL2RA
CLEC7A
ALOX5AP
LINC00968
SRPX2
MS4A6E
LIPA
AC011498.4
KCNE1
CD52
CASS4
ART4
TOP2A
CARMN

CXCR6
GALNT6
C5AR2
SIK1
CU639417.2
REEP6
GJC3
AZGP1
PRR7
IL7R
TLR8
ERRFI1
GCAT
CNTD2
F8A2
PLEK
RGS1
GSTZ1
RGS2
ALOX15B
CDKN1A
AL035406.1
ITGAX
CDKN2B
ADORA3
AC134669.2
PARP15
C5AR1
CDKN1C
ST14
OSBPL3
FGF22
ABCC3
IL24
EVI2A
NQO1
SEMA6B
SH2D7
PROB1
NDRG4
THEMIS2
LINC01094
LEP
GLIS1
MUSTN1

CR1
FOSL2
CCR5
TREM2
SGK1
AC090181.1
IQGAP2
SMIM1
AMPD3
PTPRO
TNMD
CIB2
ACP5
GPR183
C5orf58
CCR10
ADAM12
TLR2
TMEM52
MAPK8IP2
CYTIP
NUDT8
GLYCTK
FOXS1
DRD4
NLRC4
AC069368.1
LRRC46
FASN
CMTM8
WDR38
TLR7
PLIN5
AC025262.1
CD180
FMN1
KANK4
GCNT1
KLHL6
LYZ
CKB
AC092118.1
CXorf21
CCR1
SAMD4A

GSDMB
NXNL1
GPR65
ZC3H12D
MT-ND1
MT-CO3
MT-ND4
TTC36
TM7SF2
CD300LB
BCAT1
MACROD1
NCEH1
SLAMF8
SAMSN1
NPL
ARL11
CHCHD10
C9
AL080251.1
SMAP2
SESN1
PTPN22
FCGR2A
KYNU
C8orf82
RHOU
TNFSF8
ZFPM1
MERTK
MSR1
DAPP1
Mar-01
MS4A7
CD163L1
CTSB
OPLAH
GPT
COMTD1
MS4A4A
AC068987.5
PXMP2
AC135586.2
AL121845.2
IFI30

DDTL
DDT

| shared name | **Degree unDir** | **Betweenness unDir** |
|---|---|---|
| IGFN1 | **56** | **5721.198022** |
| IGHD | **57** | **4878.911318** |
| PHACTR3 | **47** | **4266.723023** |
| PKP3 | **52** | **3909.320135** |
| CLDN9 | **45** | **3859.271792** |
| SGCA | **44** | **3856.617988** |
| TRIM54 | **45** | **3753.370403** |
| GABRD | **35** | **3703.20153** |
| AC005696.4 | **46** | **3403.468823** |
| CYP4B1 | **29** | **3302.156631** |
| TF | **46** | **3253.128579** |
| ATP8A2 | **48** | **3018.943654** |
| HS3ST2 | **43** | **2989.067061** |
| AC004233.3 | **39** | **2945.053889** |
| SPATA18 | **43** | **2921.503903** |
| SPRR2F | **42** | **2904.323464** |
| LINC01956 | **32** | **2813.091647** |
| CYP2A6 | **31** | **2785.455615** |
| HSPA6 | **38** | **2604.842865** |
| PTCHD4 | **33** | **2574.869183** |
| RASL11B | **33** | **2548.384428** |
| ABCC8 | **34** | **2408.351302** |
| TREML1 | **26** | **2250.685317** |
| NRIP3 | **29** | **2204.64335** |
| AC131097.2 | **39** | **2163.677435** |
| WFDC1 | **33** | **2149.772136** |
| IGSF1 | **29** | **2062.860059** |
| LINC00942 | **34** | **1998.837108** |
| DPP10 | **29** | **1976.538993** |
| PDIA2 | **34** | **1928.467059** |
| AC092720.1 | **23** | **1877.473021** |
| CRTAM | **31** | **1843.576378** |
| AL358334.2 | **35** | **1813.956423** |
| GDF15 | **31** | **1715.388826** |
| F13A1 | **11** | **1702.322924** |
| MLIP | **33** | **1682.870368** |
| HOXA9 | **29** | **1669.73564** |
| CD300E | **27** | **1632.741937** |
| VNN1 | **28** | **1614.568475** |
| SNHG9 | **24** | **1556.130844** |
| CD69 | **30** | **1532.318236** |
| FCN2 | **29** | **1500.090683** |
| MTRNR2L12 | **15** | **1462.855007** |

| | | |
|---|---|---|
| SOX10 | **35** | **1442.234455** |
| ADAM28 | **27** | **1440.965138** |
| ADGRL3 | **30** | **1423.399825** |
| LY6K | **34** | **1419.872199** |
| SOWAHA | **28** | **1413.073218** |
| HMCN2 | **33** | **1330.952128** |
| POTEF | **24** | **1314.278571** |
| EXOC3L2 | **25** | **1260.29505** |
| AL035446.1 | **29** | **1251.057861** |
| KCNK3 | **19** | **1244.437601** |
| CA3 | **30** | **1238.984517** |
| TMIGD2 | **25** | **1174.442145** |
| MIR4458HG | **23** | **1163.31892** |
| GRIN2C | **21** | **1149.583354** |
| NCAPH | **22** | **1148.06643** |
| GAS2L2 | **29** | **1142.896305** |
| AC010997.5 | **23** | **1093.64833** |
| GFAP | **25** | **1092.787638** |
| ADRB3 | **33** | **1092.750027** |
| F2RL3 | **24** | **1091.993719** |
| CCL22 | **31** | **1089.57105** |
| MUC4 | **29** | **1039.159324** |
| AP000785.2 | **26** | **1022.284807** |
| CSRP2 | **25** | **1016.611239** |
| TRPA1 | **24** | **1016.049257** |
| FCGR2C | **23** | **986.1723774** |
| STRC | **27** | **973.6706828** |
| MT-ND3 | **12** | **971.9158535** |
| AL049634.2 | **25** | **929.6647562** |
| LINC01230 | **26** | **928.760184** |
| FKBP5 | **13** | **911.0414943** |
| CLEC4E | **9** | **876.6845637** |
| LRRC73 | **22** | **872.2046706** |
| CTLA4 | **23** | **869.296656** |
| PTGDR2 | **20** | **860.2580466** |
| CHIT1 | **25** | **859.6690544** |
| AKR1B15 | **24** | **850.0767843** |
| COL9A1 | **26** | **836.7910894** |
| ALDH4A1 | **4** | **823.9228181** |
| FCGR1A | **24** | **817.8401333** |
| UBASH3B | **23** | **817.6351026** |
| SLC2A4 | **9** | **810.4273681** |
| FOS | **22** | **796.1160165** |
| CD28 | **15** | **793.8553941** |
| MATN4 | **25** | **790.9249699** |

| | | |
|---|---|---|
| CKMT2 | **12** | **765.8821991** |
| GLYAT | **11** | **754.7247999** |
| HCN2 | **20** | **753.6291556** |
| SUSD1 | **26** | **724.2152433** |
| FCGR3A | **3** | **711.3003238** |
| SMIM25 | **3** | **709.483132** |
| ENHO | **17** | **685.0604548** |
| RCOR2 | **20** | **664.1451404** |
| CSRNP1 | **15** | **637.1679613** |
| RASL10B | **17** | **612.1660011** |
| GAPT | **20** | **608.8724266** |
| ARL4C | **13** | **607.9683845** |
| TFCP2L1 | **20** | **607.4646412** |
| PRDM1 | **20** | **594.1723961** |
| SPX | **19** | **593.6205848** |
| OLR1 | **19** | **583.6575641** |
| MSC | **20** | **583.6173583** |
| CYR61 | **19** | **583.1435562** |
| PHLDA1 | **16** | **579.7974907** |
| SIRPB1 | **22** | **579.4294131** |
| TRIM55 | **23** | **563.0736571** |
| OGDHL | **20** | **555.1543436** |
| ID4 | **19** | **552.4346249** |
| TBX1 | **18** | **544.7951334** |
| MAB21L1 | **22** | **532.6569712** |
| RANBP3L | **18** | **519.7589464** |
| DUSP1 | **15** | **493.0823621** |
| C6 | **17** | **491.2653914** |
| LINC01703 | **21** | **483.9118285** |
| TUBB2A | **20** | **462.3374989** |
| SPP1 | **17** | **462.2935689** |
| FAM181B | **14** | **451.3000126** |
| RRM2 | **13** | **444.6960733** |
| CNN1 | **20** | **442.0663918** |
| PSAT1 | **15** | **434.8127573** |
| HRC | **17** | **432.277169** |
| MUC20 | **22** | **430.3133001** |
| FAT2 | **15** | **424.9134117** |
| ADGRE2 | **19** | **421.5467161** |
| THBS1 | **16** | **411.9904237** |
| RERGL | **14** | **390.1948438** |
| ACTG2 | **22** | **382.2191347** |
| MYOCD | **17** | **370.6491398** |
| NRTN | **15** | **356.3641329** |
| HTRA3 | **21** | **340.396625** |

| | | |
|---|---|---|
| HEY2 | **17** | **340.3896806** |
| CECR2 | **14** | **339.2245602** |
| DACT2 | **16** | **338.7962031** |
| LDHD | **7** | **327.859013** |
| NEXN | **16** | **325.2844312** |
| AC110285.6 | **18** | **323.7885905** |
| SLC6A12 | **16** | **323.5100813** |
| OXT | **18** | **319.6817301** |
| SLAMF6 | **10** | **314.5021631** |
| GDF1 | **11** | **304.0586274** |
| TMEM88 | **13** | **294.7293376** |
| CERS1 | **11** | **290.4755901** |
| KCNA2 | **12** | **289.32593** |
| DMRT2 | **16** | **272.6298218** |
| SDS | **10** | **269.4107012** |
| RASSF10 | **17** | **269.2139465** |
| TWIST1 | **17** | **268.3469055** |
| BMP2 | **5** | **263.8953988** |
| GPR171 | **11** | **262.3626853** |
| CNFN | **10** | **262.2199997** |
| ALPK3 | **13** | **257.3074515** |
| IL10 | **12** | **256.5784304** |
| UCHL1 | **12** | **255.5867051** |
| G0S2 | **11** | **255.4686144** |
| ANKRD24 | **15** | **245.4165381** |
| RORB | **17** | **244.5550152** |
| TENM4 | **12** | **244.4568338** |
| MT-ATP6 | **9** | **236.6521191** |
| P2RY12 | **12** | **228.690905** |
| FHOD3 | **16** | **226.5700319** |
| TGFBR3L | **16** | **226.333867** |
| TIGAR | **11** | **223.6787755** |
| KCNB1 | **11** | **222.0459159** |
| IPCEF1 | **8** | **221.0524378** |
| LINC01770 | **11** | **220.9730839** |
| TMIGD3 | **14** | **216.8058356** |
| MT-CO2 | **9** | **211.8831898** |
| CXCR4 | **12** | **209.9864664** |
| Z99774.1 | **12** | **209.9580804** |
| PEBP4 | **11** | **206.6633188** |
| SMOC1 | **14** | **202.0757051** |
| AC107976.1 | **15** | **198.0770377** |
| CCL18 | **14** | **197.7384296** |
| BDH1 | **16** | **191.4398093** |
| SLC27A2 | **12** | **190.2986658** |

| | | |
|---|---|---|
| SLC46A2 | **13** | **187.0187274** |
| CES1 | **11** | **184.2230587** |
| FAM78B | **11** | **175.0930121** |
| LILRB3 | **11** | **174.1843409** |
| ELFN1 | **11** | **169.1525586** |
| DHRS9 | **15** | **169.1363366** |
| TBXA2R | **12** | **162.7840401** |
| MMP9 | **13** | **162.1278085** |
| HPCA | **12** | **153.4680134** |
| GFI1 | **9** | **152.9670045** |
| APLNR | **10** | **151.7955162** |
| IGSF6 | **10** | **150.566871** |
| LUCAT1 | **12** | **148.9608057** |
| ITK | **8** | **147.3625159** |
| MYCL | **12** | **147.2369671** |
| BIRC3 | **9** | **145.6063391** |
| RP11-592B15.9 | **13** | **145.2660936** |
| AC015688.5 | **11** | **142.6900643** |
| FCGR2B | **11** | **137.6037851** |
| SLC4A8 | **9** | **137.5902286** |
| TCF15 | **9** | **130.5168808** |
| DCSTAMP | **10** | **128.0558236** |
| C20orf204 | **11** | **124.7393573** |
| TM4SF19 | **11** | **121.2068884** |
| FCMR | **8** | **120.6274752** |
| MT-CYB | **9** | **116.4371171** |
| PLA2G7 | **10** | **113.6022157** |
| IL1RN | **11** | **113.3529622** |
| MT-ND2 | **8** | **112.1166071** |
| AKR1C3 | **10** | **110.6263293** |
| TNFAIP3 | **10** | **109.0553997** |
| IL2RA | **8** | **106.9479215** |
| CLEC7A | **12** | **106.8226655** |
| ALOX5AP | **5** | **106.6417771** |
| LINC00968 | **11** | **102.0197657** |
| SRPX2 | **9** | **101.874351** |
| MS4A6E | **8** | **99.47475132** |
| LIPA | **10** | **98.29753697** |
| AC011498.4 | **7** | **96.93943937** |
| KCNE1 | **6** | **91.89490777** |
| CD52 | **10** | **91.83366955** |
| CASS4 | **8** | **89.6073685** |
| ART4 | **9** | **88.31578928** |
| TOP2A | **12** | **88.06627143** |
| CARMN | **9** | **82.54956358** |

| | | |
|---|---|---|
| CXCR6 | **8** | **77.27694602** |
| GALNT6 | **9** | **76.94881747** |
| C5AR2 | **6** | **76.2605876** |
| SIK1 | **9** | **74.69291847** |
| CU639417.2 | **9** | **74.69291847** |
| REEP6 | **7** | **73.57524946** |
| GJC3 | **6** | **73.03208978** |
| AZGP1 | **6** | **72.66568842** |
| PRR7 | **8** | **70.65092057** |
| IL7R | **8** | **70.07881487** |
| TLR8 | **5** | **66.52429843** |
| ERRFI1 | **6** | **66.13902338** |
| GCAT | **10** | **65.44005717** |
| CNTD2 | **10** | **63.86034732** |
| F8A2 | **8** | **63.08975354** |
| PLEK | **5** | **62.68570725** |
| RGS1 | **7** | **62.08782741** |
| GSTZ1 | **10** | **61.32311349** |
| RGS2 | **6** | **60.48101726** |
| ALOX15B | **7** | **58.38503653** |
| CDKN1A | **7** | **56.13730366** |
| AL035406.1 | **7** | **55.48768087** |
| ITGAX | **6** | **53.57005731** |
| CDKN2B | **8** | **53.12029948** |
| ADORA3 | **8** | **53.00842601** |
| AC134669.2 | **4** | **52.39528139** |
| PARP15 | **7** | **51.14986685** |
| C5AR1 | **4** | **51.13585356** |
| CDKN1C | **5** | **48.41363262** |
| ST14 | **8** | **47.83024116** |
| OSBPL3 | **6** | **47.4231873** |
| FGF22 | **8** | **46.77516129** |
| ABCC3 | **6** | **44.3938301** |
| IL24 | **6** | **42.73338384** |
| EVI2A | **4** | **38.1449577** |
| NQO1 | **6** | **38.13310661** |
| SEMA6B | **6** | **35.88952215** |
| SH2D7 | **8** | **34.07307924** |
| PROB1 | **5** | **33.8682164** |
| NDRG4 | **7** | **33.20698845** |
| THEMIS2 | **4** | **32.60965864** |
| LINC01094 | **7** | **31.58323674** |
| LEP | **2** | **31.48204152** |
| GLIS1 | **7** | **31.34753423** |
| MUSTN1 | **8** | **29.31240091** |

| | | |
|---|---|---|
| CR1 | **3** | **28.95096133** |
| FOSL2 | **4** | **27.42790816** |
| CCR5 | **5** | **27.35919314** |
| TREM2 | **5** | **26.77059344** |
| SGK1 | **5** | **24.75084952** |
| AC090181.1 | **6** | **22.6926655** |
| IQGAP2 | **2** | **21.46438162** |
| SMIM1 | **5** | **21.17166624** |
| AMPD3 | **5** | **19.74731298** |
| PTPRO | **5** | **19.5444571** |
| TNMD | **6** | **18.11459419** |
| CIB2 | **7** | **17.66393115** |
| ACP5 | **4** | **16.91021485** |
| GPR183 | **4** | **16.53148017** |
| C5orf58 | **4** | **16.45648479** |
| CCR10 | **4** | **15.84485387** |
| ADAM12 | **4** | **14.33354054** |
| TLR2 | **3** | **14.2870153** |
| TMEM52 | **6** | **14.19007807** |
| MAPK8IP2 | **6** | **14.15806546** |
| CYTIP | **3** | **13.99794804** |
| NUDT8 | **2** | **13.71798517** |
| GLYCTK | **5** | **13.46497154** |
| FOXS1 | **3** | **13.32116863** |
| DRD4 | **4** | **13.27808131** |
| NLRC4 | **3** | **12.94179988** |
| AC069368.1 | **4** | **12.67943762** |
| LRRC46 | **4** | **12.61631979** |
| FASN | **5** | **11.89056645** |
| CMTM8 | **3** | **11.86632335** |
| WDR38 | **4** | **10.63475944** |
| TLR7 | **3** | **10.089376** |
| PLIN5 | **4** | **9.920080895** |
| AC025262.1 | **4** | **9.670828929** |
| CD180 | **5** | **9.094107442** |
| FMN1 | **4** | **8.709292111** |
| KANK4 | **3** | **8.572585206** |
| GCNT1 | **4** | **8.222491989** |
| KLHL6 | **3** | **7.96565191** |
| LYZ | **3** | **7.714521199** |
| CKB | **3** | **7.673468863** |
| AC092118.1 | **3** | **6.947549629** |
| CXorf21 | **4** | **6.120585752** |
| CCR1 | **3** | **5.328787998** |
| SAMD4A | **4** | **5.139231296** |

| | | |
|---|---|---|
| GSDMB | **3** | **3.552509937** |
| NXNL1 | **2** | **3.199043703** |
| GPR65 | **3** | **2.974850101** |
| ZC3H12D | **2** | **2.367150456** |
| MT-ND1 | **8** | **2.13771092** |
| MT-CO3 | **8** | **2.13771092** |
| MT-ND4 | **8** | **2.13771092** |
| TTC36 | **2** | **1.912761614** |
| TM7SF2 | **2** | **1.620156613** |
| CD300LB | **3** | **1.050474899** |
| BCAT1 | **3** | **0.521031746** |
| MACROD1 | **2** | **0.43977591** |
| NCEH1 | **2** | **0** |
| SLAMF8 | **2** | **0** |
| SAMSN1 | **2** | **0** |
| NPL | **2** | **0** |
| ARL11 | **2** | **0** |
| CHCHD10 | **2** | **0** |
| C9 | **1** | **0** |
| AL080251.1 | **1** | **0** |
| SMAP2 | **1** | **0** |
| SESN1 | **1** | **0** |
| PTPN22 | **1** | **0** |
| FCGR2A | **1** | **0** |
| KYNU | **1** | **0** |
| C8orf82 | **1** | **0** |
| RHOU | **1** | **0** |
| TNFSF8 | **1** | **0** |
| ZFPM1 | **1** | **0** |
| MERTK | **1** | **0** |
| MSR1 | **1** | **0** |
| DAPP1 | **1** | **0** |
| Mar-01 | **1** | **0** |
| MS4A7 | **1** | **0** |
| CD163L1 | **1** | **0** |
| CTSB | **1** | **0** |
| OPLAH | **1** | **0** |
| GPT | **1** | **0** |
| COMTD1 | **1** | **0** |
| MS4A4A | **1** | **0** |
| AC068987.5 | **1** | **0** |
| PXMP2 | **1** | **0** |
| AC135586.2 | **1** | **0** |
| AL121845.2 | **1** | **0** |
| IFI30 | **1** | **0** |

| | | |
|---|---|---|
| DDTL | **1** | **0** |
| DDT | **1** | **0** |

Table S4. Output from PANDA analysis of differentially expressed genes showing connectivities to genes and TFs

| ENSEMBL |
|---|
| ENSG00000196664 |
| ENSG00000134516 |
| ENSG00000050438 |
| ENSG00000119457 |
| ENSG00000133805 |
| ENSG00000145703 |
| ENSG00000087589 |
| ENSG00000248905 |
| ENSG00000145416 |
| ENSG00000172578 |
| ENSG00000203710 |
| ENSG00000151490 |
| ENSG00000177575 |
| ENSG00000038945 |
| ENSG00000162614 |
| ENSG00000073737 |
| ENSG00000168685 |
| ENSG00000106952 |
| ENSG00000166926 |
| ENSG00000136167 |
| ENSG00000158714 |
| ENSG00000042980 |
| ENSG00000163131 |
| ENSG00000111339 |
| ENSG00000160791 |
| ENSG00000113600 |
| ENSG00000104321 |
| ENSG00000135069 |
| ENSG00000137801 |
| ENSG00000105967 |
| ENSG00000251442 |
| ENSG00000163599 |
| ENSG00000074706 |
| ENSG00000169508 |
| ENSG00000101916 |
| ENSG00000155629 |
| ENSG00000162894 |
| ENSG00000174946 |
| ENSG00000023445 |
| ENSG00000134242 |
| ENSG00000165168 |
| ENSG00000177675 |
| ENSG00000234511 |
| ENSG00000171860 |
| ENSG00000080546 |
| ENSG00000154127 |

ENSG00000081237
ENSG00000082074
ENSG00000146070
ENSG00000163823
ENSG00000187474
ENSG00000000005
ENSG00000169313
ENSG00000057657
ENSG00000173200
ENSG00000070190
ENSG00000126860
ENSG00000168995
ENSG00000115956
ENSG00000135838
ENSG00000140030
ENSG00000172215
ENSG00000090104
ENSG00000172243
ENSG00000175857
ENSG00000155307
ENSG00000149256
ENSG00000101307
ENSG00000112299
ENSG00000010671
ENSG00000072694
ENSG00000149177
ENSG00000066294
ENSG00000070882
ENSG00000137491
ENSG00000178562
ENSG00000143226
ENSG00000086570
ENSG00000124491
ENSG00000173391
ENSG00000115919
ENSG00000060982
ENSG00000113263
ENSG00000129450
ENSG00000136689
ENSG00000116574
ENSG00000110077
ENSG00000110079
ENSG00000144959
ENSG00000149124
ENSG00000163071
ENSG00000244682
ENSG00000102359
ENSG00000110848

ENSG00000154277
ENSG00000155659
ENSG00000090382
ENSG00000131747
ENSG00000164188
ENSG00000155926
ENSG00000107798
ENSG00000143119
ENSG00000196139
ENSG00000118785
ENSG00000169442
ENSG00000167851
ENSG00000115112
ENSG00000145569
ENSG00000106868
ENSG00000171848
ENSG00000162739
ENSG00000187210
ENSG00000123338
ENSG00000134460
ENSG00000246430
ENSG00000115165
ENSG00000178789
ENSG00000118515
ENSG00000162676
ENSG00000136634
ENSG00000139629
ENSG00000140749
ENSG00000234147
ENSG00000166927
ENSG00000134294
ENSG00000186407
ENSG00000120280
ENSG00000235568
ENSG00000121933
ENSG00000137462
ENSG00000178199
ENSG00000179593
ENSG00000180509
ENSG00000091106
ENSG00000162892
ENSG00000134061
ENSG00000173110
ENSG00000148848
ENSG00000248323
ENSG00000128045
ENSG00000153208
ENSG00000127507

ENSG00000146192
ENSG00000166523
ENSG00000108846
ENSG00000020577
ENSG00000164935
ENSG00000132965
ENSG00000147255
ENSG00000125845
ENSG00000197405
ENSG00000103034
ENSG00000081320
ENSG00000132854
ENSG00000249628
ENSG00000112280
ENSG00000109943
ENSG00000150337
ENSG00000139289
ENSG00000196604
ENSG00000111181
ENSG00000164733
ENSG00000084070
ENSG00000078237
ENSG00000167664
ENSG00000147883
ENSG00000145107
ENSG00000039537
ENSG00000141506
ENSG00000121966
ENSG00000198848
ENSG00000227471
ENSG00000224397
ENSG00000204577
ENSG00000116990
ENSG00000087495
ENSG00000102962
ENSG00000135094
ENSG00000160255
ENSG00000152213
ENSG00000131095
ENSG00000215375
ENSG00000095970
ENSG00000006071
ENSG00000099977
ENSG00000175352
ENSG00000177301
ENSG00000133063
ENSG00000170801
ENSG00000169020

ENSG00000203747
ENSG00000130176
ENSG00000228672
ENSG00000105219
ENSG00000131730
ENSG00000165644
ENSG00000168237
ENSG00000070388
ENSG00000149418
ENSG00000187185
ENSG00000183134
ENSG00000225518
ENSG00000198712
ENSG00000134830
ENSG00000160862
ENSG00000166165
ENSG00000158716
ENSG00000169738
ENSG00000213999
ENSG00000134775
ENSG00000156381
ENSG00000141294
ENSG00000173253
ENSG00000188778
ENSG00000182103
ENSG00000198840
ENSG00000164879
ENSG00000167701
ENSG00000136918
ENSG00000160339
ENSG00000178821
ENSG00000255198
ENSG00000091513
ENSG00000148357
ENSG00000169710
ENSG00000179588
ENSG00000174332
ENSG00000121152
ENSG00000115286
ENSG00000105427
ENSG00000272573
ENSG00000089847
ENSG00000260260
ENSG00000250479
ENSG00000006638
ENSG00000178814
ENSG00000179772
ENSG00000136425

ENSG00000123689
ENSG00000213937
ENSG00000176894
ENSG00000198732
ENSG00000167799
ENSG00000122971
ENSG00000101405
ENSG00000161267
ENSG00000150471
ENSG00000125652
ENSG00000235169
ENSG00000149809
ENSG00000163395
ENSG00000145113
ENSG00000181856
ENSG00000188859
ENSG00000176402
ENSG00000122691
ENSG00000183476
ENSG00000136383
ENSG00000184363
ENSG00000167969
ENSG00000198763
ENSG00000247516
ENSG00000099624
ENSG00000069696
ENSG00000166823
ENSG00000134020
ENSG00000216921
ENSG00000170293
ENSG00000133315
ENSG00000100577
ENSG00000100116
ENSG00000249669
ENSG00000175183
ENSG00000073605
ENSG00000258711
ENSG00000129757
ENSG00000171303
ENSG00000197444
ENSG00000172201
ENSG00000255974
ENSG00000198888
ENSG00000128309
ENSG00000184451
ENSG00000146147
ENSG00000198944
ENSG00000214456

ENSG00000160886
ENSG00000225285
ENSG00000152082
ENSG00000187730
ENSG00000175497
ENSG00000140284
ENSG00000132932
ENSG00000161509
ENSG00000131188
ENSG00000103175
ENSG00000100146
ENSG00000269028
ENSG00000134548
ENSG00000176945
ENSG00000115255
ENSG00000260001
ENSG00000204052
ENSG00000099822
ENSG00000233493
ENSG00000140678
ENSG00000142973
ENSG00000124159
ENSG00000223802
ENSG00000196421
ENSG00000166816
ENSG00000099795
ENSG00000260750
ENSG00000111404
ENSG00000130528
ENSG00000213563
ENSG00000189431
ENSG00000242866
ENSG00000171119
ENSG00000125878
ENSG00000008735
ENSG00000078399
ENSG00000258910
ENSG00000116285
ENSG00000167680
ENSG00000163017
ENSG00000135547
ENSG00000108823
ENSG00000184058
ENSG00000181019
ENSG00000167874
ENSG00000134817
ENSG00000147573
ENSG00000225968

ENSG00000172425
ENSG00000075426
ENSG00000167771
ENSG00000130775
ENSG00000127533
ENSG00000130513
ENSG00000168913
ENSG00000120129
ENSG00000099954
ENSG00000138100
ENSG00000198963
ENSG00000244094
ENSG00000100985
ENSG00000141052
ENSG00000137267
ENSG00000122254
ENSG00000198727
ENSG00000102575
ENSG00000012779
ENSG00000121905
ENSG00000185615
ENSG00000244694
ENSG00000118503
ENSG00000164488
ENSG00000270017
ENSG00000124762
ENSG00000159423
ENSG00000180660
ENSG00000178860
ENSG00000142178
ENSG00000142871
ENSG00000158445
ENSG00000161911
ENSG00000116741
ENSG00000170345
ENSG00000144655
ENSG00000096060
ENSG00000174697
ENSG00000188042
ENSG00000211898

| SYMBOL | Freq_Gene | Freq_TF |
|---|---|---|
| TLR7 | 216 | 305 |
| DOCK2 | 214 | 305 |
| SLC4A8 | 220 | 302 |
| SLC46A2 | 209 | 299 |
| AMPD3 | 215 | 296 |
| IQGAP2 | 213 | 295 |
| CASS4 | 210 | 293 |
| FMN1 | 212 | 291 |
| 1-Mar | 213 | 290 |
| KLHL6 | 216 | 289 |
| CR1 | 205 | 289 |
| PTPRO | 214 | 287 |
| CD163 | 211 | 285 |
| MSR1 | 216 | 284 |
| NEXN | 219 | 283 |
| DHRS9 | 213 | 283 |
| IL7R | 209 | 283 |
| TNFSF8 | 207 | 283 |
| MS4A6E | 212 | 282 |
| LCP1 | 213 | 281 |
| SLAMF8 | 209 | 281 |
| ADAM28 | 207 | 281 |
| CTSS | 213 | 280 |
| ART4 | 207 | 279 |
| CCR5 | 216 | 278 |
| C9 | 219 | 277 |
| TRPA1 | 212 | 277 |
| PSAT1 | 209 | 277 |
| THBS1 | 213 | 276 |
| TFEC | 211 | 275 |
| LINC01094 | 210 | 275 |
| CTLA4 | 209 | 274 |
| IPCEF1 | 212 | 273 |
| GPR183 | 213 | 272 |
| TLR8 | 212 | 272 |
| PIK3AP1 | 220 | 271 |
| FCMR | 214 | 271 |
| GPR171 | 214 | 271 |
| BIRC3 | 213 | 271 |
| PTPN22 | 210 | 271 |
| CYBB | 210 | 271 |
| CD163L1 | 210 | 271 |
| C5orf58 | 215 | 270 |
| C3AR1 | 212 | 270 |
| SESN1 | 218 | 269 |
| UBASH3B | 217 | 268 |

| | | |
|---|---|---|
| PTPRC | 215 | 268 |
| FYB1 | 212 | 268 |
| PLA2G7 | 222 | 267 |
| CCR1 | 220 | 267 |
| FPR3 | 219 | 267 |
| TNMD | 211 | 267 |
| P2RY12 | 211 | 267 |
| PRDM1 | 221 | 266 |
| PARP15 | 218 | 266 |
| DAPP1 | 209 | 266 |
| EVI2A | 216 | 265 |
| SIGLEC7 | 216 | 265 |
| PLEK | 215 | 265 |
| NPL | 208 | 265 |
| GPR65 | 218 | 264 |
| CXCR6 | 216 | 264 |
| RGS1 | 212 | 264 |
| CLEC7A | 210 | 264 |
| GAPT | 210 | 264 |
| SAMSN1 | 211 | 263 |
| TENM4 | 216 | 262 |
| SIRPB1 | 215 | 262 |
| VNN1 | 215 | 262 |
| BTK | 212 | 262 |
| FCGR2B | 211 | 262 |
| PTPRJ | 221 | 261 |
| CD84 | 211 | 261 |
| OSBPL3 | 221 | 260 |
| SLCO2B1 | 221 | 260 |
| CD28 | 217 | 260 |
| FCGR2A | 215 | 260 |
| FAT2 | 210 | 260 |
| F13A1 | 218 | 259 |
| OLR1 | 213 | 259 |
| KYNU | 211 | 259 |
| BCAT1 | 218 | 258 |
| ITK | 216 | 258 |
| SIGLEC9 | 216 | 258 |
| IL1RN | 213 | 258 |
| RHOU | 212 | 258 |
| MS4A6A | 216 | 257 |
| MS4A4A | 212 | 257 |
| NCEH1 | 212 | 257 |
| GLYAT | 208 | 257 |
| SPATA18 | 216 | 256 |
| FCGR2C | 217 | 255 |
| SRPX2 | 213 | 255 |
| CD69 | 212 | 255 |

| | | |
|---|---|---|
| UCHL1 | 211 | 255 |
| VSIG4 | 209 | 255 |
| LYZ | 216 | 254 |
| TOP2A | 215 | 254 |
| RANBP3L | 215 | 254 |
| SLA | 214 | 254 |
| LIPA | 219 | 253 |
| CD53 | 215 | 253 |
| AKR1C3 | 211 | 252 |
| SPP1 | 213 | 251 |
| CD52 | 220 | 250 |
| CD300A | 216 | 250 |
| TFCP2L1 | 227 | 249 |
| FAM105A | 226 | 249 |
| SUSD1 | 223 | 249 |
| RRM2 | 216 | 249 |
| SLAMF6 | 214 | 249 |
| GCNT1 | 223 | 248 |
| NCKAP1L | 220 | 248 |
| IL2RA | 213 | 248 |
| LINC00968 | 212 | 248 |
| CYTIP | 210 | 248 |
| CD300LB | 221 | 247 |
| SGK1 | 219 | 247 |
| GFI1 | 216 | 247 |
| IL10 | 212 | 247 |
| GALNT6 | 224 | 246 |
| IGSF6 | 220 | 246 |
| AL035446. | 215 | 246 |
| MS4A7 | 212 | 246 |
| SLC38A2 | 221 | 245 |
| CD300E | 209 | 245 |
| CXorf21 | 219 | 243 |
| NFAM1 | 215 | 243 |
| TMIGD3 | 214 | 243 |
| TLR2 | 210 | 243 |
| ZC3H12D | 221 | 242 |
| ALOX15B | 231 | 240 |
| KCNE1 | 219 | 240 |
| NLRC4 | 221 | 239 |
| IL24 | 219 | 239 |
| CD180 | 214 | 239 |
| HSPA6 | 210 | 239 |
| ADAM12 | 209 | 239 |
| LUCAT1 | 207 | 239 |
| RASL11B | 226 | 237 |
| MERTK | 219 | 235 |
| ADGRE2 | 217 | 235 |

| | | |
|---|---|---|
| FGD2 | 238 | 233 |
| CLEC4E | 213 | 233 |
| ABCC3 | 237 | 231 |
| SAMD4A | 228 | 231 |
| DCSTAMP | 221 | 231 |
| ALOX5AP | 215 | 231 |
| IGSF1 | 214 | 231 |
| BMP2 | 234 | 229 |
| C5AR1 | 225 | 229 |
| NDRG4 | 178 | 228 |
| STK17B | 221 | 226 |
| KANK4 | 224 | 225 |
| LINC00942 | 219 | 225 |
| COL9A1 | 210 | 225 |
| CRTAM | 213 | 224 |
| FCGR1A | 211 | 224 |
| PHLDA1 | 219 | 223 |
| POTEF | 216 | 223 |
| SLC6A12 | 238 | 219 |
| CTSB | 233 | 219 |
| SMAP2 | 220 | 219 |
| TIGAR | 238 | 216 |
| TMIGD2 | 247 | 215 |
| CDKN2B | 220 | 215 |
| TM4SF19 | 219 | 214 |
| C6 | 212 | 213 |
| PIK3R5 | 293 | 209 |
| CXCR4 | 252 | 207 |
| CES1 | 215 | 207 |
| AKR1B15 | 207 | 207 |
| SMIM25 | 237 | 206 |
| LILRB3 | 236 | 206 |
| MYCL | 277 | 205 |
| PHACTR3 | 220 | 202 |
| CCL22 | 216 | 201 |
| SDS | 230 | 200 |
| ITGB2 | 239 | 199 |
| ARL11 | 251 | 197 |
| GFAP | 190 | 196 |
| MYL5 | 163 | 195 |
| TREM2 | 214 | 194 |
| ABCC8 | 182 | 194 |
| DDT | 169 | 194 |
| NRIP3 | 243 | 192 |
| KCNA2 | 259 | 191 |
| CHIT1 | 248 | 187 |
| HTRA3 | 168 | 187 |
| ATP5I | 162 | 187 |

| | | |
|---|---|---|
| FCGR3A | 250 | 186 |
| CNN1 | 198 | 186 |
| PROB1 | 167 | 186 |
| CNTD2 | 184 | 185 |
| CKMT2 | 172 | 185 |
| COMTD1 | 164 | 185 |
| GLYCTK | 171 | 184 |
| FGF22 | 169 | 183 |
| ST14 | 241 | 182 |
| AC092118. | 181 | 182 |
| PTGDR2 | 170 | 182 |
| LINC01703 | 169 | 182 |
| MT-CO2 | 163 | 182 |
| C5AR2 | 253 | 179 |
| AZGP1 | 181 | 179 |
| CKB | 177 | 179 |
| DUSP23 | 174 | 179 |
| DCXR | 171 | 179 |
| MEF2B | 175 | 178 |
| FHOD3 | 192 | 177 |
| ANKRD9 | 168 | 177 |
| LRRC46 | 166 | 177 |
| DMRT2 | 166 | 177 |
| ADRB3 | 173 | 176 |
| FAM181B | 169 | 175 |
| MT-ND3 | 157 | 175 |
| CA3 | 173 | 173 |
| GPT | 171 | 173 |
| WDR38 | 167 | 173 |
| FCN2 | 167 | 173 |
| TMEM52 | 167 | 173 |
| SNHG9 | 164 | 173 |
| TF | 207 | 172 |
| HMCN2 | 179 | 172 |
| FASN | 171 | 172 |
| ZFPM1 | 169 | 172 |
| GLIS1 | 166 | 172 |
| NCAPH | 233 | 171 |
| NDUFS7 | 187 | 171 |
| CNFN | 172 | 171 |
| MUSTN1 | 169 | 170 |
| ANKRD24 | 182 | 169 |
| SNHG19 | 178 | 169 |
| CHCHD10 | 177 | 169 |
| TBXA2R | 176 | 169 |
| OPLAH | 172 | 169 |
| FOXS1 | 170 | 169 |
| CIB2 | 167 | 169 |

| | | |
|---|---|---|
| G0S2 | 163 | 169 |
| CLDN9 | 170 | 167 |
| PXMP2 | 173 | 166 |
| SMOC1 | 170 | 166 |
| NUDT8 | 169 | 166 |
| ACADS | 168 | 166 |
| OXT | 164 | 166 |
| BDH1 | 309 | 165 |
| ADGRL3 | 231 | 165 |
| ALKBH7 | 186 | 165 |
| SMIM1 | 173 | 165 |
| TM7SF2 | 162 | 165 |
| IGFN1 | 174 | 164 |
| MUC4 | 172 | 164 |
| SLC2A4 | 172 | 164 |
| FAM78B | 172 | 164 |
| GJC3 | 165 | 164 |
| TWIST1 | 173 | 163 |
| SH2D7 | 173 | 163 |
| ALPK3 | 172 | 163 |
| PKP3 | 170 | 163 |
| ECI1 | 167 | 163 |
| MT-ND2 | 165 | 163 |
| MIR4458H | 190 | 162 |
| ATP5D | 174 | 162 |
| DRD4 | 172 | 162 |
| MESP1 | 163 | 162 |
| PEBP4 | 173 | 161 |
| AC131097. | 173 | 161 |
| CMTM8 | 172 | 161 |
| MACROD1 | 180 | 160 |
| GSTZ1 | 179 | 160 |
| GCAT | 173 | 160 |
| CARMN | 168 | 160 |
| CSRP2 | 167 | 160 |
| GSDMB | 253 | 159 |
| AL358334. | 160 | 159 |
| CDKN1C | 174 | 158 |
| KCNK3 | 173 | 158 |
| OGDHL | 172 | 158 |
| ID4 | 270 | 157 |
| CYP2A6 | 166 | 157 |
| MT-ND1 | 156 | 157 |
| MPST | 173 | 156 |
| CCR10 | 172 | 156 |
| MLIP | 213 | 155 |
| SOWAHA | 179 | 155 |
| PLIN5 | 173 | 155 |

| | | |
|---|---|---|
| LY6K | 187 | 154 |
| LINC01770 | 175 | 154 |
| MZT2B | 174 | 154 |
| GABRD | 173 | 154 |
| DPP10 | 220 | 153 |
| SLC27A2 | 198 | 153 |
| ATP8A2 | 172 | 153 |
| GRIN2C | 171 | 153 |
| PRR7 | 176 | 152 |
| WFDC1 | 174 | 152 |
| SOX10 | 173 | 152 |
| MTRNR2L1 | 167 | 152 |
| SPX | 200 | 151 |
| MUC20 | 177 | 151 |
| REEP6 | 173 | 151 |
| TGFBR3L | 171 | 151 |
| LRRC73 | 164 | 151 |
| HCN2 | 172 | 150 |
| TMEM238 | 159 | 150 |
| ITGAX | 247 | 149 |
| CYP4B1 | 224 | 149 |
| MATN4 | 180 | 149 |
| CERS1 | 176 | 149 |
| C20orf204 | 173 | 149 |
| LDHD | 184 | 148 |
| NDUFB7 | 173 | 147 |
| AC092720. | 171 | 147 |
| RERGL | 208 | 146 |
| HRC | 176 | 146 |
| C8orf82 | 172 | 146 |
| RASSF10 | 169 | 145 |
| STRC | 196 | 144 |
| NRTN | 184 | 144 |
| TCF15 | 178 | 144 |
| MAPK8IP2 | 170 | 144 |
| HOXA9 | 183 | 142 |
| LINC01956 | 175 | 142 |
| ERRFI1 | 278 | 141 |
| SEMA6B | 234 | 141 |
| ACTG2 | 190 | 141 |
| HEY2 | 172 | 141 |
| SGCA | 195 | 140 |
| TBX1 | 171 | 139 |
| NQO1 | 225 | 136 |
| TMEM88 | 181 | 136 |
| APLNR | 175 | 136 |
| TRIM55 | 203 | 133 |
| ELFN1 | 207 | 132 |

| | | |
|---|---|---|
| TTC36 | 194 | 132 |
| FOSL2 | 307 | 131 |
| RCOR2 | 187 | 128 |
| THEMIS2 | 236 | 125 |
| F2RL3 | 251 | 122 |
| GDF15 | 265 | 116 |
| ENHO | 173 | 116 |
| DUSP1 | 176 | 114 |
| CECR2 | 235 | 111 |
| TRIM54 | 200 | 110 |
| RORB | 274 | 103 |
| SPRR2F | 209 | 102 |
| MMP9 | 191 | 101 |
| MYOCD | 192 | 97 |
| TUBB2A | 176 | 97 |
| HS3ST2 | 173 | 95 |
| MT-CYB | 143 | 93 |
| ACP5 | 322 | 92 |
| ALOX5 | 281 | 91 |
| HPCA | 275 | 90 |
| PDIA2 | 214 | 90 |
| PTCHD4 | 296 | 89 |
| TNFAIP3 | 199 | 84 |
| DACT2 | 195 | 84 |
| AC107976. | 197 | 82 |
| CDKN1A | 267 | 80 |
| ALDH4A1 | 300 | 79 |
| MAB21L1 | 198 | 75 |
| MSC | 242 | 74 |
| SIK1 | 208 | 60 |
| CYR61 | 186 | 60 |
| KCNB1 | 272 | 57 |
| TREML1 | 220 | 55 |
| RGS2 | 300 | 54 |
| FOS | 179 | 54 |
| CSRNP1 | 206 | 52 |
| FKBP5 | 198 | 47 |
| LEP | 188 | 39 |
| ARL4C | 178 | 35 |
| IGHD | 179 | 28 |

Table S5. List of 80 genes, their rank and the analysis datasets

| Gene | TF |
|---|---|
| AC005696.4 | 0 |
| AL121845.2 | 0 |
| CLDN9 | 0 |
| IGFN1 | 0 |
| IGHD | 0 |
| PHACTR3 | 0 |
| PKP3 | 0 |
| SGCA | 0 |
| TRIM54 | 0 |
| GABRD | 0 |
| CYP4B1 | 0 |
| ATP8A2 | 0 |
| TF | 0 |
| MUC20 | 0 |
| MUC4 | 0 |
| DMRT2 | 1 |
| LINC01230 | 0 |
| AL049634.2 | 0 |
| SIRPB1 | 0 |
| AC004233.3 | 0 |
| CERS1 | 0 |
| GDF1 | 0 |
| DDT | 0 |
| DDTL | 0 |
| AC068987.5 | 0 |
| LINC01094 | 0 |
| MT-ATP6 | 0 |
| MT-ND2 | 0 |
| MT-ND4 | 0 |
| AL080251.1 | 0 |
| TNFAIP3 | 0 |
| OXA1L | 0 |
| CCR10 | 0 |
| ACTR2 | 0 |
| AMN | 0 |
| LRRC46 | 0 |
| H2AFY | 0 |
| ANKRD16 | 0 |
| FAM78B | 0 |
| MEF2B | 1 |
| SLC4A8 | 0 |
| OSBPL3 | 0 |
| NEXN | 0 |
| SESN1 | 0 |
| UBASH3B | 0 |
| FCGR2C | 0 |

| | |
|---|---|
| MSR1 | 0 |
| LIPA | 0 |
| MS4A6A | 0 |
| PARP15 | 0 |
| PRDM1 | 1 |
| IRF3 | 1 |
| SOX30 | 1 |
| BCL6 | 1 |
| SOX3 | 1 |
| HOXC6 | 1 |
| SPI1 | 1 |
| FOXA1 | 1 |
| LEF1 | 1 |
| HOXD10 | 1 |
| ARRDC2 | 0 |
| NFIL3 | 0 |
| EIF2B3 | 0 |
| RASSF9 | 0 |
| SEC61A1 | 0 |
| RDH16 | 0 |
| GPRIN2 | 0 |
| MED27 | 0 |
| C2CD4C | 0 |
| GALNT9 | 0 |
| PAX9 | 1 |
| REL | 1 |
| PAX1 | 1 |
| RELA | 1 |
| PAX2 | 1 |
| RELB | 1 |
| NFYC | 1 |
| NFIA | 1 |
| NFIB | 1 |
| CEBPZ | 1 |

| Top10_rank | | Dataset |
|---|---|---|
| | 9 | Centiscape_betweeness, Centiscape_degree |
| | 9 | geneNet_median, Limma |
| | 5 | Centiscape_betweeness, Centiscape_degree, geneNet_max |
| | 1 | Centiscape_betweeness, Centiscape_degree |
| | 2 | Centiscape_betweeness, Centiscape_degree |
| | 3 | Centiscape_betweeness, Centiscape_degree |
| | 4 | Centiscape_betweeness, Centiscape_degree |
| | 6 | Centiscape_betweeness, Centiscape_degree |
| | 7 | Centiscape_betweeness, Centiscape_degree |
| | 8 | Centiscape_betweeness |
| | 10 | Centiscape_betweeness |
| | 4 | Centiscape_degree |
| | 6 | Centiscape_degree |
| | 1 | geneNet_max |
| | 2 | geneNet_max |
| | 3 | geneNet_max |
| | 4 | geneNet_max |
| | 5 | geneNet_max |
| | 6 | geneNet_max |
| | 7 | geneNet_max |
| | 9 | geneNet_max |
| | 10 | geneNet_max |
| | 1 | geneNet_median |
| | 2 | geneNet_median |
| | 3 | geneNet_median |
| | 4 | geneNet_median |
| | 5 | geneNet_median |
| | 6 | geneNet_median |
| | 7 | geneNet_median |
| | 8 | geneNet_median |
| | 10 | geneNet_median |
| | 1 | Limma |
| | 2 | Limma |
| | 3 | Limma |
| | 4 | Limma |
| | 5 | Limma |
| | 6 | Limma |
| | 7 | Limma |
| | 8 | Limma |
| | 9 | Limma |
| | 1 | PANDA_GENE |
| | 2 | PANDA_GENE |
| | 3 | PANDA_GENE |
| | 4 | PANDA_GENE |
| | 5 | PANDA_GENE |
| | 6 | PANDA_GENE |

| | |
|---|---|
| 7 | PANDA_GENE |
| 8 | PANDA_GENE |
| 9 | PANDA_GENE |
| 10 | PANDA_GENE |
| 1 | PANDA_TF |
| 2 | PANDA_TF |
| 3 | PANDA_TF |
| 4 | PANDA_TF |
| 5 | PANDA_TF |
| 6 | PANDA_TF |
| 7 | PANDA_TF |
| 8 | PANDA_TF |
| 9 | PANDA_TF |
| 10 | PANDA_TF |
| 1 | alpaca_gene_Modularity |
| 2 | alpaca_gene_Modularity |
| 3 | alpaca_gene_Modularity |
| 4 | alpaca_gene_Modularity |
| 5 | alpaca_gene_Modularity |
| 6 | alpaca_gene_Modularity |
| 7 | alpaca_gene_Modularity |
| 8 | alpaca_gene_Modularity |
| 9 | alpaca_gene_Modularity |
| 10 | alpaca_gene_Modularity |
| 1 | alpaca_TF_Modularity |
| 2 | alpaca_TF_Modularity |
| 3 | alpaca_TF_Modularity |
| 4 | alpaca_TF_Modularity |
| 5 | alpaca_TF_Modularity |
| 6 | alpaca_TF_Modularity |
| 7 | alpaca_TF_Modularity |
| 8 | alpaca_TF_Modularity |
| 9 | alpaca_TF_Modularity |
| 10 | alpaca_TF_Modularity |

Table S6. Results from Pubmed analysis of 80 genes (top 10 from each analysis)

| Gene | count_pubmed |
|---|---|
| TF | 582 |
| DDT | 280 |
| RELA | 90 |
| LIPA | 78 |
| IRF3 | 72 |
| REL | 72 |
| IGHD | 36 |
| BCL6 | 34 |
| LEF1 | 31 |
| TNFAIP3 | 22 |
| PAX2 | 21 |
| MSR1 | 20 |
| FOXA1 | 20 |
| SPI1 | 17 |
| MUC4 | 15 |
| NFIA | 15 |
| NFIL3 | 14 |
| CERS1 | 13 |
| RELB | 11 |
| PRDM1 | 10 |
| HOXC6 | 8 |
| MT-ND2 | 6 |
| SESN1 | 6 |
| DMRT2 | 6 |
| NFIB | 6 |
| CYP4B1 | 5 |
| MT-ATP6 | 5 |
| MS4A6A | 4 |
| CCR10 | 4 |
| OSBPL3 | 4 |
| SOX3 | 4 |
| AMN | 4 |
| RDH16 | 4 |
| SGCA | 3 |
| NFYC | 3 |
| GALNT9 | 3 |
| PAX1 | 3 |
| SEC61A1 | 3 |
| IGFN1 | 2 |
| ACTR2 | 2 |
| ATP8A2 | 2 |
| FCGR2C | 2 |
| MT-ND4 | 2 |
| MUC20 | 2 |
| FAM78B | 2 |
| GDF1 | 2 |

| | |
|---|---|
| H2AFY | 2 |
| HOXD10 | 2 |
| MEF2B | 2 |
| RASSF9 | 2 |
| SIRPB1 | 2 |
| UBASH3B | 2 |
| CLDN9 | 1 |
| GABRD | 1 |
| LINC01230 | 1 |
| OXA1L | 1 |
| TRIM54 | 1 |
| ARRDC2 | 1 |
| CEBPZ | 1 |
| NEXN | 1 |
| PAX9 | 1 |
| AC004233.3 | 0 |
| AC005696.4 | 0 |
| AC068987.5 | 0 |
| AL049634.2 | 0 |
| AL080251.1 | 0 |
| AL121845.2 | 0 |
| ANKRD16 | 0 |
| C2CD4C | 0 |
| EIF2B3 | 0 |
| GPRIN2 | 0 |
| LINC01094 | 0 |
| LRRC46 | 0 |
| MED27 | 0 |
| PARP15 | 0 |
| PHACTR3 | 0 |
| PKP3 | 0 |
| SLC4A8 | 0 |
| SOX30 | 0 |
| DDTL | 0 |

note
other_term (transcription factor)
other_term (Dichlorodiphenyltrichloroethane)

other_term(light-intensity physical activity (LIPA))

other_term (relaxation training [REL])
other_term (diopathic growth hormone deficiency)

other_term (solid gel coupling agent)

Table S7. Results of Disgenet analysis of 80 genes (top 10 from each analysis)

| symbol | number_disease_disgenet_all |
|---|---|
| RELA | 483 |
| BCL6 | 309 |
| PAX2 | 216 |
| TNFAIP3 | 212 |
| LEF1 | 211 |
| MUC4 | 183 |
| IRF3 | 174 |
| TF | 168 |
| GABRD | 154 |
| SOX3 | 151 |
| FOXA1 | 138 |
| GDF1 | 136 |
| LIPA | 130 |
| PRDM1 | 126 |
| SGCA | 111 |
| MSR1 | 108 |
| UBASH3B | 104 |
| NFIB | 102 |
| NFIA | 95 |
| REL | 90 |
| PAX1 | 89 |
| DDT | 86 |
| HOXD10 | 83 |
| HOXC6 | 80 |
| CEBPZ | 76 |
| OXA1L | 75 |
| RELB | 72 |
| FCGR2C | 71 |
| SPI1 | 58 |
| CCR10 | 57 |
| ATP8A2 | 55 |
| NFIL3 | 54 |
| PAX9 | 53 |
| CERS1 | 49 |
| ACTR2 | 45 |
| AMN | 43 |
| MEF2B | 42 |
| EIF2B3 | 36 |
| NEXN | 36 |
| CYP4B1 | 35 |
| PKP3 | 31 |
| SEC61A1 | 30 |
| SOX30 | 28 |
| IGHD | 21 |
| MUC20 | 20 |
| MED27 | 17 |

| | |
|---|---|
| SESN1 | 13 |
| NFYC | 11 |
| SIRPB1 | 10 |
| DMRT2 | 10 |
| OSBPL3 | 10 |
| MS4A6A | 10 |
| GALNT9 | 8 |
| TRIM54 | 8 |
| IGFN1 | 6 |
| PHACTR3 | 6 |
| CLDN9 | 5 |
| RASSF9 | 5 |
| PARP15 | 5 |
| LINC01094 | 5 |
| RDH16 | 4 |
| SLC4A8 | 4 |
| ARRDC2 | 4 |
| FAM78B | 4 |
| ANKRD16 | 3 |
| C2CD4C | 3 |
| GPRIN2 | 2 |
| LRRC46 | 2 |
| DDTL | 2 |
| LINC01230 | 1 |

| Total diseases | Disease_specificity_index_disgenet_all |
|---|---|
| 4884 | 0.272401636 |
| 4884 | 0.324990536 |
| 4884 | 0.367146726 |
| 4884 | 0.369347426 |
| 4884 | 0.369904089 |
| 4884 | 0.386666119 |
| 4884 | 0.392603547 |
| 4884 | 0.39673499 |
| 4884 | 0.406979192 |
| 4884 | 0.409295346 |
| 4884 | 0.419894489 |
| 4884 | 0.421613265 |
| 4884 | 0.426925476 |
| 4884 | 0.430604965 |
| 4884 | 0.445527956 |
| 4884 | 0.448753748 |
| 4884 | 0.453197069 |
| 4884 | 0.455483239 |
| 4884 | 0.46385365 |
| 4884 | 0.470219202 |
| 4884 | 0.47153468 |
| 4884 | 0.475571671 |
| 4884 | 0.479752018 |
| 4884 | 0.484086276 |
| 4884 | 0.490125243 |
| 4884 | 0.491684657 |
| 4884 | 0.496490795 |
| 4884 | 0.498137452 |
| 4884 | 0.521947617 |
| 4884 | 0.523995217 |
| 4884 | 0.528200451 |
| 4884 | 0.53036077 |
| 4884 | 0.53256147 |
| 4884 | 0.541800251 |
| 4884 | 0.551826224 |
| 4884 | 0.557178693 |
| 4884 | 0.559949034 |
| 4884 | 0.578097817 |
| 4884 | 0.578097817 |
| 4884 | 0.581414489 |
| 4884 | 0.595702793 |
| 4884 | 0.599563272 |
| 4884 | 0.607686082 |
| 4884 | 0.641556056 |
| 4884 | 0.647300319 |
| 4884 | 0.66643433 |

| | |
|---|---|
| 4884 | 0.698018135 |
| 4884 | 0.717686088 |
| 4884 | 0.728907341 |
| 4884 | 0.728907341 |
| 4884 | 0.728907341 |
| 4884 | 0.728907341 |
| 4884 | 0.755178934 |
| 4884 | 0.755178934 |
| 4884 | 0.789048909 |
| 4884 | 0.789048909 |
| 4884 | 0.810514363 |
| 4884 | 0.810514363 |
| 4884 | 0.810514363 |
| 4884 | 0.810514363 |
| 4884 | 0.836785956 |
| 4884 | 0.836785956 |
| 4884 | 0.836785956 |
| 4884 | 0.836785956 |
| 4884 | 0.870655931 |
| 4884 | 0.870655931 |
| 4884 | 0.918392978 |
| 4884 | 0.918392978 |
| 4884 | 0.918392978 |
| 4884 | 1 |

Table S8. Results from GWAS Catalog analysis on 80 genes (top 10 genes from each analysis)

| gene |
|---|
| NFIA |
| NFIB |
| UBASH3B |
| BCL6 |
| MSR1 |
| LINC01094 |
| PAX2 |
| TF |
| MED27 |
| ATP8A2 |
| GALNT9 |
| LIPA |
| PHACTR3 |
| RELB |
| SPI1 |
| SLC4A8 |
| TNFAIP3 |
| HOXC6 |
| NFIL3 |
| FCGR2C |
| DMRT2 |
| LEF1 |
| OSBPL3 |
| SIRPB1 |
| MEF2B |
| PRDM1 |
| ARRDC2 |
| MUC4 |
| SESN1 |
| FAM78B |
| RASSF9 |
| CYP4B1 |
| MS4A6A |
| RDH16 |
| ANKRD16 |
| PAX9 |
| ACTR2 |
| AMN |
| LINC01230 |
| SGCA |
| TRIM54 |
| HOXD10 |
| IRF3 |
| REL |
| PARP15 |
| SOX30 |

FOXA1
IGFN1
IGHD
NEXN
RELA
CERS1
DDTL
NFYC
C2CD4C
EIF2B3
GDF1
PAX1
PKP3
OXA1L

| count_gwas_traits |
| ---: |
| 120 |
| 91 |
| 73 |
| 53 |
| 50 |
| 45 |
| 41 |
| 40 |
| 36 |
| 28 |
| 28 |
| 28 |
| 28 |
| 28 |
| 28 |
| 25 |
| 25 |
| 24 |
| 24 |
| 23 |
| 21 |
| 20 |
| 20 |
| 19 |
| 18 |
| 17 |
| 16 |
| 16 |
| 16 |
| 15 |
| 14 |
| 13 |
| 13 |
| 13 |
| 11 |
| 11 |
| 9 |
| 8 |
| 8 |
| 7 |
| 7 |
| 6 |
| 6 |
| 6 |
| 5 |
| 5 |

4
4
4
4
4
3
3
3
2
2
2
2
2
1
